# Major histocompatibility complex class I-restricted protection against murine cytomegalovirus requires missing-self recognition by the natural killer cell inhibitory Ly49 receptors

**DOI:** 10.1101/753970

**Authors:** Bijal A. Parikh, Michael D. Bern, Sytse J. Piersma, Liping Yang, Diana L. Beckman, Jennifer Poursine-Laurent, Beatrice Plougastel Douglas, Wayne M. Yokoyama

## Abstract

Viruses have evolved strategies that highlight critical, intertwined host immune mechanisms. As postulated by the missing-self hypothesis, natural killer (NK) cells and major histocompatibility complex class I (MHC-I)-restricted cytotoxic T lymphocytes (CTLs) have opposing requirements for ubiquitously expressed MHC-I molecules. Since NK cell MHC-I-specific Ly49 inhibitory receptors prevent killing of cells with normal MHC-I, viruses evading CTLs by down-regulating MHC-I should be vulnerable to NK cells. However, definitive integrated *in vivo* evidence for this interplay has been lacking, in part due to receptor polymorphism and a proposed second function of Ly49 receptors in licensing NK cells via self-MHC-I. Here we generated mice lacking specific Ly49 inhibitory receptors to show their essential role in licensing and controlling murine cytomegalovirus (MCMV) infection *in vivo* in an MHC-restricted manner. When MCMV cannot down-regulate MHC-I, NK cells cannot control infection that instead is mediated by CTLs, as predicted by the missing-self hypothesis.

## Introduction

Viruses and their hosts are engaged in an evolutionary “arms race,” highlighting host immune mechanisms. A critical mechanism involves recognition of virus-infected cells by cytotoxic lymphocytes that trigger exocytic release of perforin and granzymes to kill infected cells. While virus-specific CD8^+^ cytotoxic T lymphocytes (CTLs) recognize viral peptides presented by MHC class I molecules (MHC-I) on infected cells, viruses can specifically down-regulate MHC-I, nearly ubiquitously expressed as self, to evade CTL effector responses [1, 2]. The missing-self hypothesis predicts that NK cells should kill cells that downregulate MHC-I [3], as supported by MHC-I-specific inhibitory NK cell receptors that prevent NK cell killing of cells with normal MHC-I expression *in vitro* [4, 5]. However, the role of inhibitory NK cell receptors during *in vivo* responses is still poorly understood.

In mice, the Ly49 NK cell receptor family is encoded in a gene cluster in the NK gene complex (NKC) on mouse chromosome 6 [6]. While Ly49s display profound allelic polymorphisms with several haplotypes and allelic forms, [7] most Ly49s mediate inhibitory function in effector responses through their cytoplasmic immunoreceptor tyrosine-based inhibitory motifs (ITIMs) [8]. Inhibitory Ly49 receptors bind MHC-I alleles but their specificities are incompletely understood regarding *in vivo* function and are reportedly identical, *e.g.*, Ly49A^B6^, Ly49C^B6^, Ly49G^B6^, and Ly49I^B6^ recognize H2D^d^ [9, 10]. Ly49s are stochastically expressed on overlapping subsets of NK cells with individual NK cells simultaneously expressing two or more Ly49s. Only receptors specific for self-MHC-I *in vivo* appear to provide a second function in conferring the licensed phenotype whereby licensed NK cells exhibit enhanced responsiveness to stimulation through activation receptors *in vitro* [11]. However, due to the potential role of other receptors [12, 13], definitive evidence that MHC-I-dependent licensing plays a role in NK cell function *in vivo* has not been established.

In contrast to inhibitory Ly49s, Ly49 activation receptors lack ITIMs and are coupled to immunoreceptor tyrosine-based activation motif (ITAM)-containing chains, *e.g.* DAP12 [8]. The Ly49H^B6^ activation receptor is responsible for genetic resistance of C57BL/6 (B6) mice to murine cytomegalovirus (MCMV) infection, providing vital early viral control, even in mice with intact adaptive immunity [14]. Ly49H^B6^ recognizes an MCMV-encoded MHC-I-like molecule, m157 [15]. The Ly49P1^NOD/Ltj^, Ly49L^BALB^, and Ly49D2^PWK/Pas^ activation receptors serve similar functions, albeit with ligands distinct from m157 [16]. Additionally, the inhibitory Ly49I^129^ receptor also binds m157 [15, 17], suggesting that inhibitory Ly49 receptors play critical roles in viral control, although effects on licensed NK cells also require consideration.

Indeed, prior studies suggested that licensed NK cells hamper MCMV control through inhibitory receptors for host MHC-I [18]. However, other studies suggest that inhibitory Ly49s may enhance MCMV control, but these approaches have utilized mouse strains with poorly characterized Ly49s and Ly49 depleting antibodies with unclear specificities and that also affect total NK cell number, confounding interpretations [16,19,20]. Thus, the *in vivo* role of licensed NK cells and inhibitory Ly49s in viral infections is incompletely understood.

Herein, we used CRISPR-Cas9 to simultaneously and cleanly target multiple Ly49 receptors, allowing definitive evaluation of the role of Ly49s in missing-self rejection and resistance to MCMV infection. In turn, we studied how MCMV modulates these host responses.

## Results

### MHC-restricted, NK cell-dependent protection against MCMV in mice

To investigate the role of NK cells in MCMV control in different MHC backgrounds, we utilized C57BL/10 (B10, H2^b^) and B10.D2 (H2^d^) MHC-congenic mouse strains that are closely related to B6 (H2^b^) and share Ly49 haplotypes [21]. Susceptibility at day 4 post-infection (d4 p.i.) to wild-type (WT)-MCMV in B10 mice was NK cell-dependent, and antibody blocking studies showed protection against WT-MCMV was dependent upon Ly49H (**Fig 1A**) as shown for B6 mice [14]. To examine other mechanisms of viral resistance applicable to MCMV isolates from the wild lacking m157 [17], we used Δm157-MCMV that contains a single nucleotide deletion in m157 that prevents full-length expression. Although B10 mice were unable to control Δm157-MCMV infection at d4 p.i., MHC-congenic B10.D2 (H2^d^) mice were resistant (**Fig 1B**), demonstrating an MHC-restricted effect.

**Fig 1.**
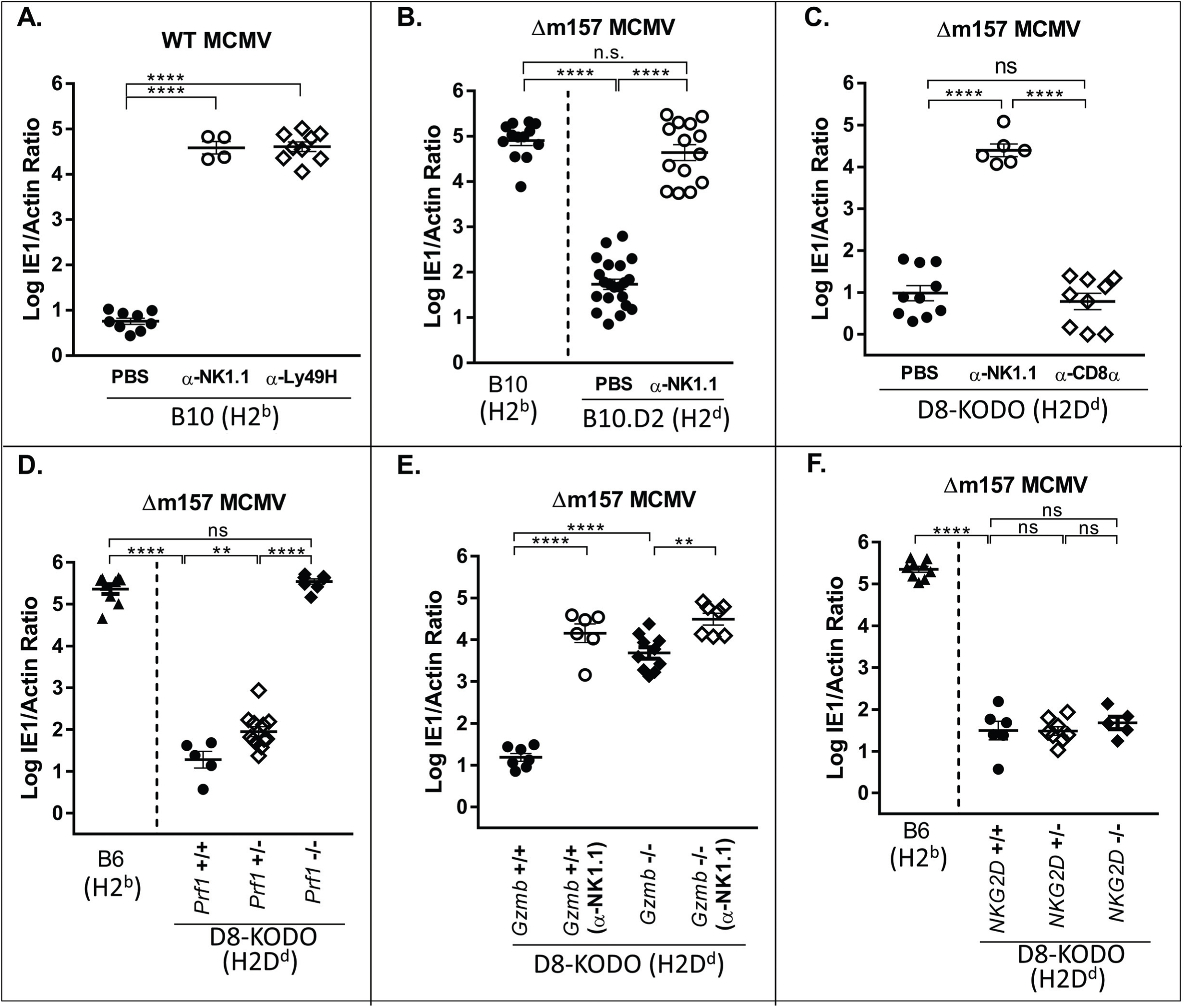
H2^d^-dependent protection against MCMV lacking m157 requires cytotoxic NK cells. Splenic viral titers in mice depleted of total NK cells, CD8^+^ T-cells, or where Ly49H^+^ receptors on NK cells were blocked. Data represent a composite of two independent experiments with 3-7 mice per group with individual points representing a single mouse using (A) H2^b^-expressing B10 mice, (B) B10 and H2^d^-expressing B10.D2 mice, (C) H2D^d^-expressing (D8-KODO) mice, (D-F) D8-KODO mice with wild-type, heterozygous, or no expression of (D) perforin (*Prf1*), (E) granzyme B (*Gzmb*), and (F) NKG2D (*Klrk1*).

To isolate the MHC-restricted effect to a single MHC-I allele, we showed Δm157-MCMV resistance in B6 mice transgenically expressing only H2D^d^, in the genetic absence of H2K^b^ and H2D^b^ (D8-KODO), comparable to B10.D2 mice (**Fig 1C**), indicating resistance was specifically due to H2D^d^. Resistance was clearly dependent upon NK cells but not CD8^+^ T cells, as shown by antibody depletion of NK or CD8^+^ T cells, respectively (**Fig 1B, C**). Thus, MHC-restricted resistance to Δm157-MCMV is due to H2D^d^ and is NK cell-dependent.

Perforin-deficient (*Prf1^-^*^/-^) D8-KODO mice were as susceptible as wild type (WT) B6 mice whereas both perforin WT (*Prf1^+/+^*) and perforin heterozygous (*Prf1^+/^*^-^) D8-KODO mice controlled Δm157-MCMV (**Fig 1D**). Likewise, granzyme-deficient D8-KODO mice could not control Δm157-MCMV (**Fig 1E**). Although NKG2D enhances NK cell responses to MCMV in B6 mice [22, 23], we found no change in viral control by NKG2D-deficient D8-KODO mice as compared to D8-KODO mice (**Fig 1F**). Thus, NK cell-dependent control of Δm157-MCMV requires cytotoxicity, implying direct target contact, but NKG2D was not required.

### NK cell resistance requires Ly49 receptor expression

Since Ly49s recognize MHC-I, we assessed their candidacy for being responsible for the MHC-I-restricted, NK-dependent resistance to Δm157-MCMV by using CRISPR-Cas9 to target their deletion directly in B6 zygotes. When we used a guide RNA (gRNA) intentionally chosen for its promiscuity for several *Ly49*s, we generated ΔLy49-1 mice with two distinct deletions: 1) 149kb deletion between *Ly49a* and *Ly49g* with an out-of-frame fusion; and 2) 66Kb deletion between two pseudogenes, *Ly49n* (*Klra14-ps*) and *Ly49k* (*Klra11-ps*), such that *Ly49h* was deleted (**Fig 2A**). Flow cytometry confirmed the loss of Ly49A, Ly49C, Ly49G, and Ly49H expression. Ly49D expression was markedly decreased but its coding sequence was intact, suggesting an as yet unidentified locus control region within one of the deleted segments. We also generated single and compound *Ly49* deleted mice, detailed below (**Fig S1** and **Fig S2**). Predicted potential off-target sites were absent by PCR amplification and sequencing (**Table S1)**. To further eliminate any off-target effects and genetic mosaicism, we backcrossed all CRISPR-Cas9 founder mice to WT B6 for two generations followed by additional crosses to KODO mice, then D8 (H2D^d^) transgenic mice, to generate the indicated homozygous Ly49 knockout mice on the D8-KODO background.

**Fig 2.**
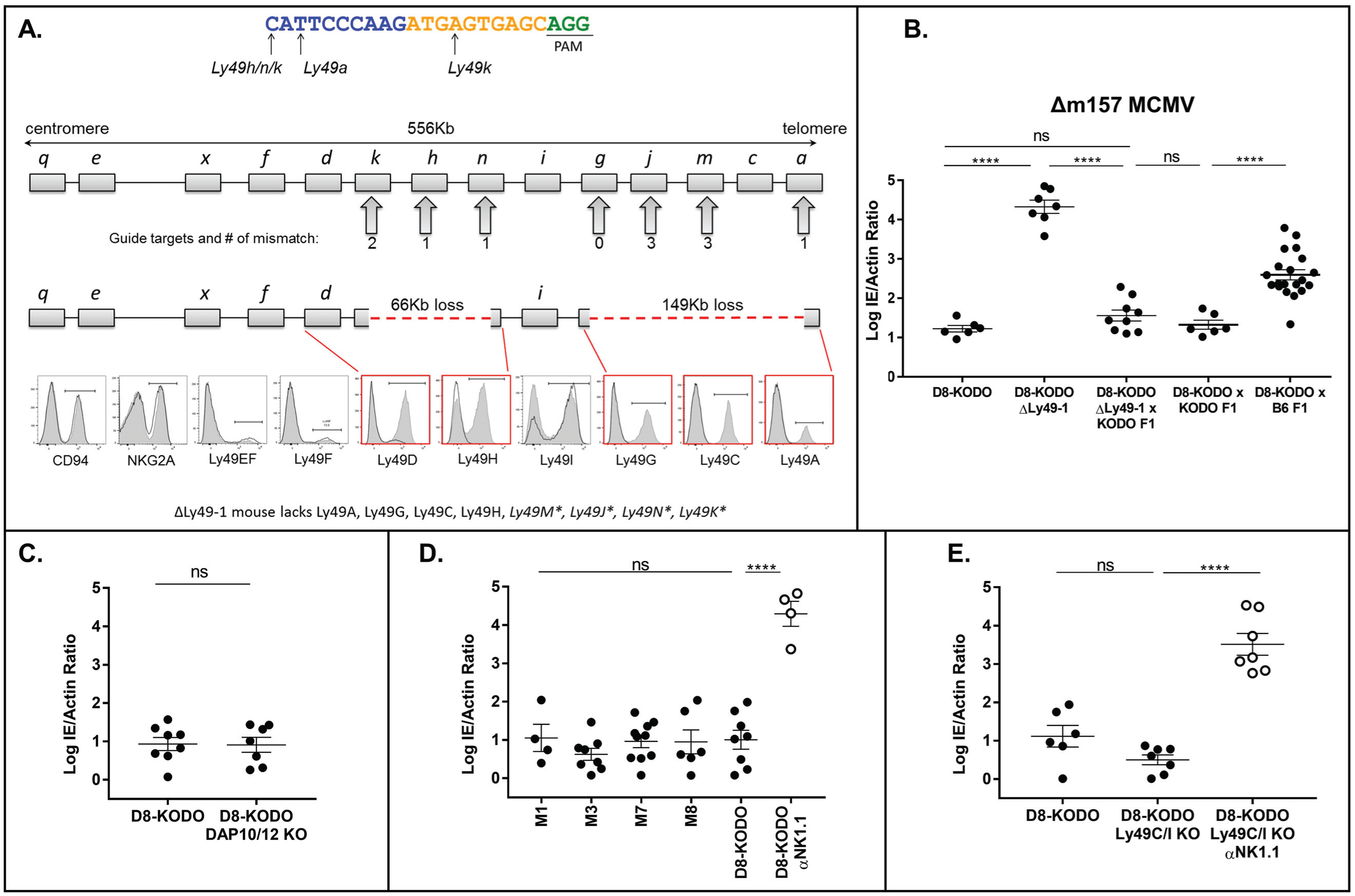
NK cell resistance to Δm157-MCMV requires specific Ly49 receptors. (A) The CRISPR sgRNA used for targeting is shown; PAM site underlined, 10nt core in yellow. Mismatches between the sgRNA and targeted *Ly49s* are represented below the guide. B6 *Ly49* cluster with the number of predicted mismatches shown below. Loss of genetic regions is indicated as a red dashed line; flow cytometry histograms indicate wild-type D8-KODO (shaded) and ΔLy49-1 D8-KODO (solid line) NK cell expression. (B-E) Δm157-MCMV splenic titers; data represent a composite of at least two independent experiments with 3-7 mice per group: (B) D8-KODO, ΔLy49-1 D8-KODO and the indicated F1 mice; (C-E) D8-KODO mice (C) lacking DAP10 and DAP12, (D) with *Ly49m* deletions and (E) lacking Ly49C and Ly49I.

MHC-restricted, NK cell-dependent resistance to Δm157-MCMV in D8-KODO mice was absent in ΔLy49-1 D8-KODO mice (**Fig 2B**). Additionally, KODO mice with intact *Ly49*s complemented ΔLy49-1 D8-KODO mice as their F1 hybrids showed fully restored Δm157-MCMV resistance (**Fig 2B**), indicating that heterozygous expression of Ly49A, Ly49C, Ly49G, Ly49H (**Fig S2**), deleted in ΔLy49-1, was sufficient for resistance. Similarly, {(D8-KODO x KODO) F1 hybrids} were also resistant (**Fig 2B**), indicating that H2D^d^ heterozygosity is sufficient for antiviral protection when *Ly49* genes were intact.

### Inhibitory Ly49A and Ly49G receptors are required for H2D^d^-dependent resistance

To decipher which Ly49 receptor(s) are involved in Δm157-MCMV resistance in D8-KODO mice, we first considered Ly49 activation receptors. However, a cross between resistant D8-KODO mice with susceptible B6 (H2^b^) mice generating heterozygosity for H2D^d^ and H2^b^ resulted in an intermediate infection phenotype (**Fig 2B**), highlighting an apparent role of MHC-I context for antiviral protection and contrasting MCMV resistance explained by Ly49 activation receptors when complete reversal was found in crossing susceptible and resistant mice [16,24,25]. Nonetheless, since ΔLy49-1 D8-KODO mice had low levels of Ly49D and lacked *Ly49h* and both receptors require DAP12 (and DAP10 to a lesser extent) for expression, including Ly49H-mediated resistance to MCMV, we evaluated the role of Ly49D and Ly49H by studying D8-KODO mice deficient in DAP10 and DAP12. For reasons not immediately clear, Ly49A, Ly49F and Ly49G expression was markedly decreased **(Fig S2)**. Regardless, these mice were still resistant **(Fig 2C)**, indicating that activation signals through DAP10/DAP12 are not required, consistent with the lack of involvement of NKG2D (**Fig 1E**) which also requires DAP10 or DAP12 for surface expression, depending on its isoform (**Fig S2)** [26]. Thus, Ly49 activation receptors are surprisingly not required for resistance to Δm157-MCMV in D8-KODO mice.

Having ruled out Ly49 activation receptors, we assessed Ly49 pseudogenes which could theoretically contribute to resistance by splicing events, which has been considered for other *Ly49* genes [27, 28]. To test *Ly49m* (*Klra13-ps*), harboring a premature stop codon (third exon), we generated four independent CRISPR-Cas9 KO strains. When crossed to D8-KODO background, all four lines showed resistance to Δm157-MCMV, indicating no apparent role for *Ly49m* (**Fig 2D**). Of the other remaining regions of the *Ly49* cluster disrupted in the ΔLy49-1 strain, we did not pursue *Ly49j*, *Ly49k* and *Ly49n* because they are predicted to encode severely truncated proteins, not expressed as receptors on NK cells [27].

We then turned our attention to Ly49 inhibitory receptors. Since Ly49C and Ly49I reportedly have specificity for H2^d^ [9, 10], they could be relevant to ΔLy49-1 D8-KODO mice which express Ly49I but not Ly49C (**Fig 2A**). However, lack of these two Ly49s in a new D8-KODO KO strain had no effect on H2D^d^ resistance to Δm157-MCMV which remained entirely dependent upon NK cells (**Fig 2E**).

Finally, we evaluated the potential contribution of Ly49A and Ly49G, since both are deleted in ΔLy49-1 mice and there are substantial data indicating that they both recognize H2D^d^ [29, 30]. In D8-KODO mice, Ly49A depletion had no significant change in viral titers while Ly49G depletion moderately increased Δm157-MCMV levels (**Fig 3A**). Interestingly, depletion of both Ly49A and Ly49G led to a major loss of viral control, similar to complete NK cell depletion (**Fig 3A**), but Ly49G depletion results in approximately 50% decrease of all NK cells (**Fig S2**). Nonetheless, Ly49D depletion, also affecting 50-60% of NK cells, did not alter viral resistance, suggesting a potential redundant contribution of Ly49A and Ly49G, independent of quantitative NK cell loss.

**Fig 3.**
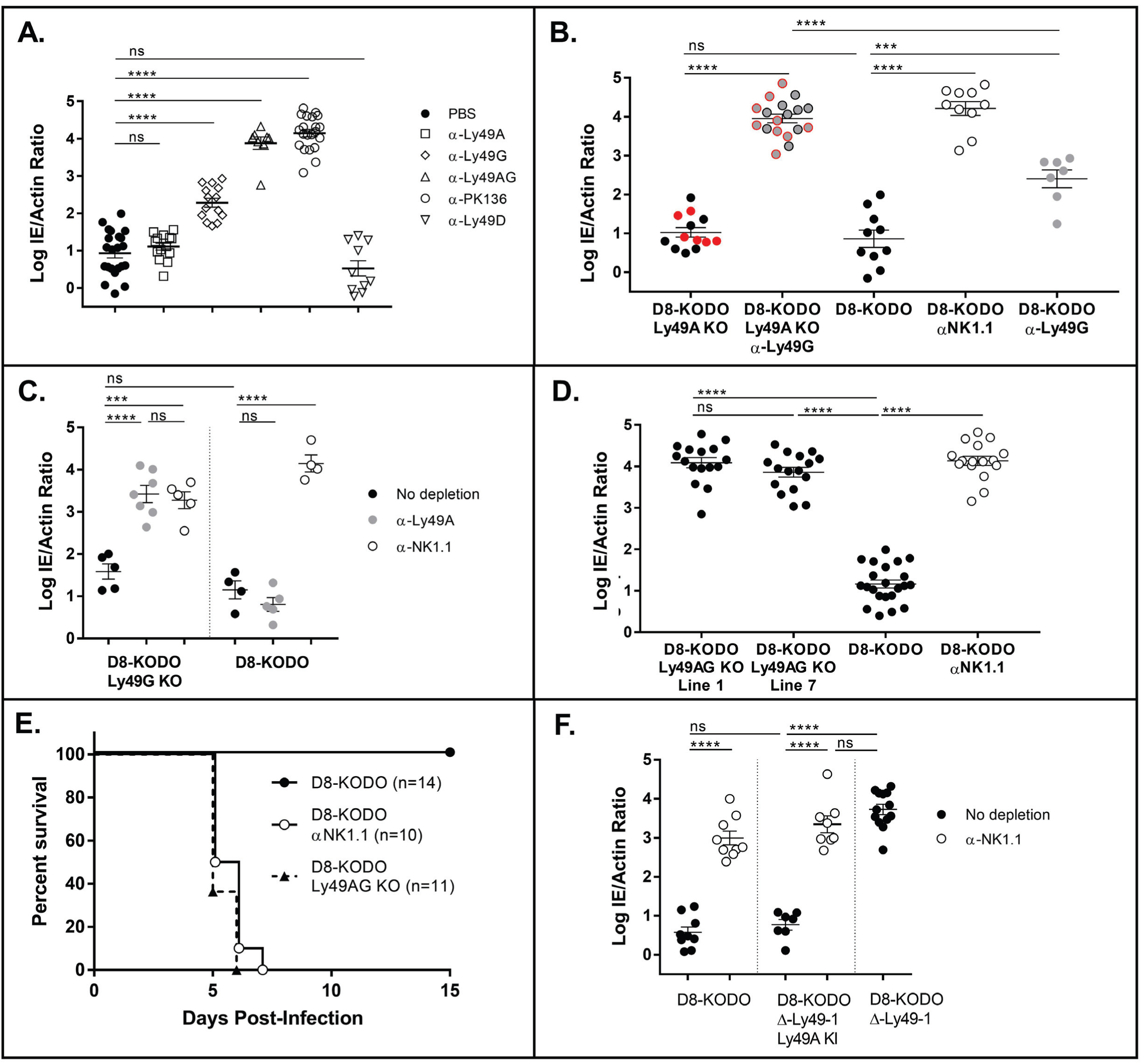
Ly49G and Ly49A are required for H2Dd-dependent Δm157-MCMV resistance. (A-D, F) Δm157-MCMV splenic titers; data represent a composite of multiple independent experiments, as noted, with 3-7 mice per group with individual points representing a single mouse. Mice used were (A) D8-KODO, five experiments; and (B) two lines lacking Ly49A (black vs. red), three experiments; (C) Ly49G knockout mice, representative of two experiments; (D) two lines lacking Ly49A and Ly49G, five experiments; (F) and ΔLy49-1 D8-KODO with or without the Ly49A knockin, two experiments. (E) Composite survival analysis of D8-KODO and Ly49A/G knockout mice; two experiments.

To definitively determine the role of Ly49A and Ly49G in resistance, we derived new Ly49 KO strains on the D8-KODO background (**Fig S1** and **Fig S2**). While deletion of Ly49A alone was insufficient to reverse MCMV resistance, Ly49G depletion in these mice allowed MCMV titers higher than levels with Ly49G depletion in Ly49A-sufficient mice and similar to that with NK cell depletion (**Fig 3B**). Reciprocally, Ly49G KO mice were resistant and Ly49A depletion reversed resistance, again comparable to NK cell depletion (**Fig 3C**). Finally, we studied two mouse strains with knockouts of both Ly49A and Ly49G (Ly49AG KO) (**Fig S1**) which displayed relatively unchanged NK cell numbers, repertoire of other Ly49s, and development (**Fig S2**). Infection of either Ly49AG KO strain resulted in viral titers similar to NK cell depletion of D8-KODO mice (**Fig 3D**) and lethality (**Fig 3E**). To provide additional evidence, we generated Ly49A knockin (KI) mice (into *Ncr1* on chromosome 7) which expressed Ly49A at near normal levels on all NKp46^+^ NK cells (**Fig S3**). ΔLy49-1 Ly49A KI D8-KODO mice demonstrated NK-cell dependent resistance to Δm157-MCMV, unlike susceptible parental ΔLy49-1 D8-KODO mice, showing complementation by Ly49A (**Fig 3F**). Therefore, Ly49A and Ly49G act redundantly in H2D^d^ mice to promote NK cell-dependent resistance to Δm157-MCMV, unequivocally establishing the protective role of inhibitory Ly49 receptors in NK cell-dependent viral control.

### NK cell licensing and missing-self rejection both require Ly49A and Ly49G receptors in D8-KODO mice

To further delineate the potential mechanism for MHC-restricted viral resistance, we investigated whether loss of Ly49A and Ly49G would have an impact on NK cell licensing by stimulating NK cells from D8-KODO, Ly49AG KO D8-KODO, and TKO (KODO β2m^-/-^) mice with plate-bound anti-NK1.1 for interferon gamma production [11]. Remarkably, the total NK cell pool in Ly49AG KO D8-KODO mice exhibited significantly reduced IFN-gamma production compared to D8-KODO NK cells and similar to unlicensed NK cells from TKO mice (Fig 4A, B). These results strongly suggest that Ly49A and Ly49G are required for NK cell licensing through H2D^d^.

**Fig 4.**
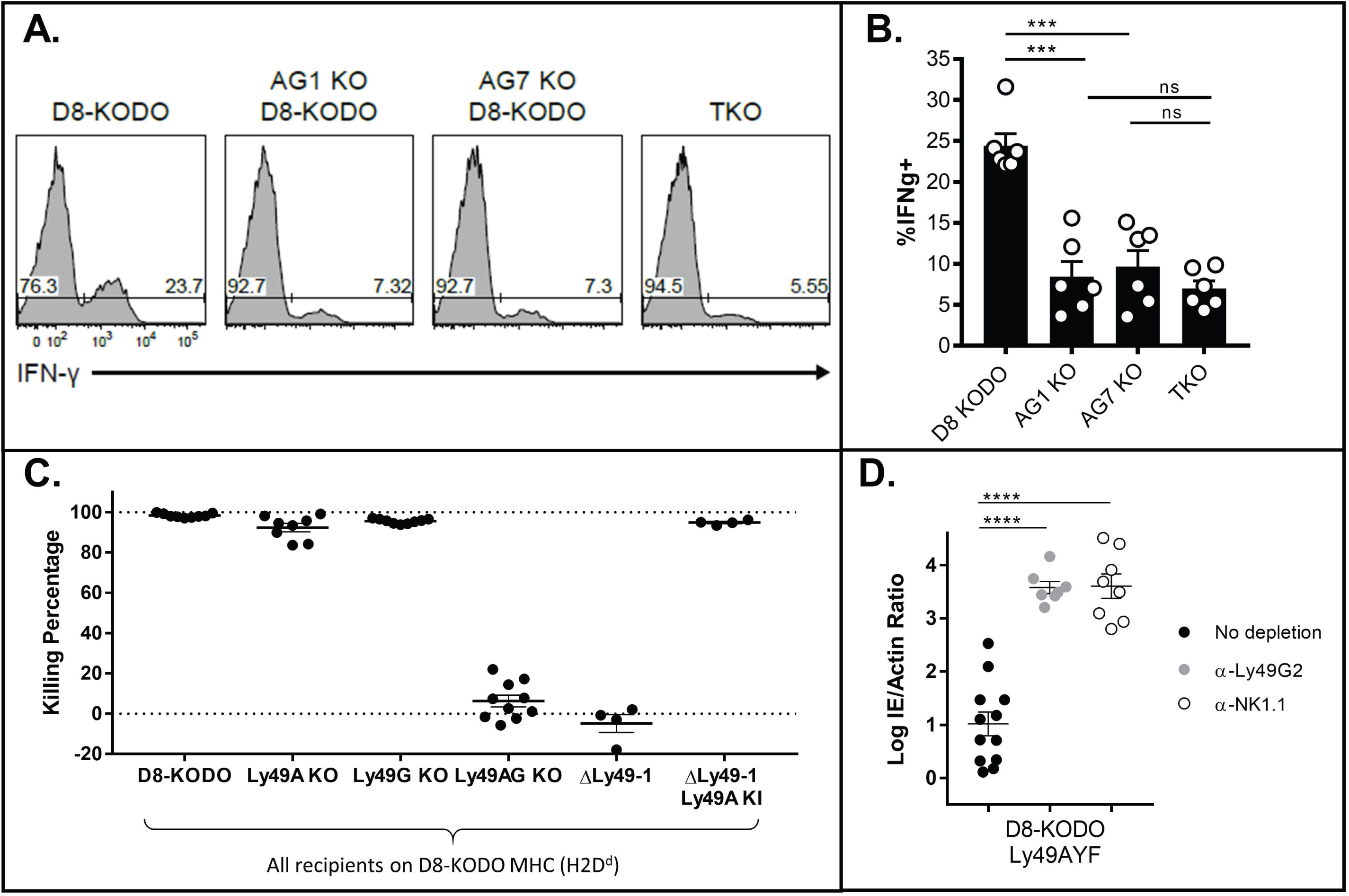
NK cell licensing and missing-self rejection in D8-KODO mice both require Ly49A and Ly49G. (A) Representative histogram plots depicting the frequency of interferon gamma positive (IFNg+) NK cells following NK1.1 stimulation in D8-KODO, Ly49A/G KO mice, and TKO (KODO β2m KO) mice. (B) Composite plot of IFNg+ frequencies from (A); two independent experiments with 3 mice per group. (C) *In vivo* cytotoxicity of KODO or D8-KODO splenocytes. Data are cumulative over two independent experiments with 4-5 recipient mice per group. (D) Δm157-MCMV infection as in **Fig 2B**; three independent experiments.

We next tested whether loss of inhibitory Ly49s would affect missing-self recognition *in vivo* as suggested by *in vitro* studies [5, 31] and anti-Ly49 antibody experiments *in vivo* [32]. Upon injection, labelled KODO (MHC-I-deficient) splenocytes were effectively cleared in D8-KODO mice that were either WT, or lacked only Ly49A or only Ly49G (**Fig 4C, S4**), in an NK cell-dependent manner as shown by NK cell depletion (**Fig S4A, B**). Importantly, ΔLy49-1 D8-KODO mice and Ly49AG KO D8-KODO mice were unable to reject KODO targets while complementation of ΔLy49-1 D8-KODO mice with Ly49A KI restored this capacity. Thus, these results strongly suggest Ly49A and Ly49G are redundant missing-self receptors in D8-KODO mice as they enable NK cells to reject missing-self targets through NK cell licensing.

### Inhibitory Ly49s mediate protection from Δm157-MCMV is ITIM-dependent

We recently generated KI mice carrying Ly49A with a non-functional ITIM (termed Ly49AYF), demonstrating that effector inhibition and licensing are both mediated by the Ly49A ITIM [31]. Ly49AYF D8-KODO mice were resistant to Δm157-MCMV (**Fig 4D**) similar to D8-KODO mice but, in contrast to D8-KODO mice (**Fig 3A, B**), Ly49G depletion led to significantly elevated viral titers, similar to anti-NK1.1 depletion (**Fig 4D**). These data also mirror Ly49G depletion of Ly49A-KO D8-KODO mice (**Fig 3B**), suggesting that Ly49A mediates resistance to Δm157-MCMV in an ITIM-dependent manner. We also observed that Ly49G-depleted Ly49AYF D8-KODO mice were unable to reject missing-self targets, indicating that the ITIM is required for Ly49A to mediate missing-self recognition *in vivo* (**Fig S4**). These findings strongly suggest that Ly49A mediates resistance to Δm157-MCMV in an ITIM-dependent manner.

### MCMV infection generates targets for missing-self rejection

We next investigated the mechanism by which Δm157-MCMV might contribute to the Ly49 effects. We initially focused on two MCMV immunoevasins known to downmodulate MHC-I, *m06* and *m152*, [33] although they have variable effects dependent upon MHC-I alleles [34]. To establish their relevance here, we first infected SV40-immortalized D8-KODO MEFs with a GFP-expressing Δm157-MCMV, indicating that H2D^d^ was down-regulated in GFP-positive infected cells (**Fig 5A, B**). Next, we produced mutant Δm157-GFP viruses deficient for either *m06*, *m152*, or both. Remarkably, H2D^d^ downmodulation was completely abrogated only when both m06 and m152 were targeted (**Fig 5A, B**). In contrast, WT m152 (Δm06Δm157 MCMV) or WT m06 (Δm152Δm157 MCMV) decreased surface expression of H2D^d^ in infected cells, consistent with virus-free overexpression systems demonstrating that both m06 [35] and m152 [36] redundantly promote MHC down-regulation.

**Fig 5.**
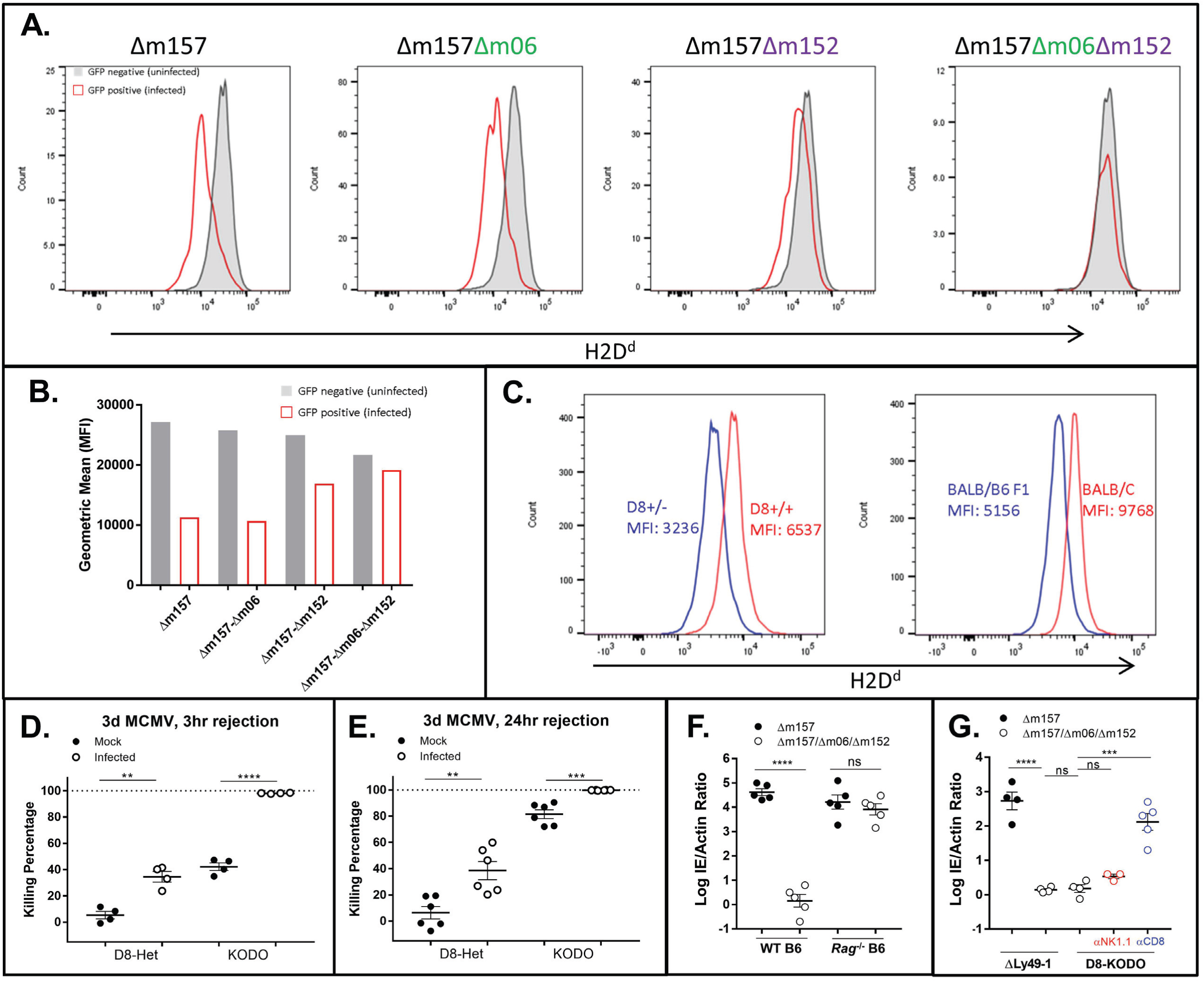
Down-regulation of H2D^d^ through *m06* and *m152* affects host MCMV control. (A,B) 24hr *in vitro* infection and flow cytometric evaluation of D8-KODO MEFs with GFP-expressing MCMV lacking the indicated ORFs; representative of two independent infections. (C) MFI of H2D^d^ in mice homozygous and heterozygous for the D8 transgene (*left*), compared to BALB/c (H2^d/d^) and BALB/c x B6 F1 (H2^b/d^) mice (*right*); representative of two independent experiments. (D) 3 hr or (E) 24 hr *in vivo* cytotoxicity with KODO or D8-KODO heterozygous (D8-Het; H2D^d+/-^) donor cells. Composite of (E) one or (D) two independent experiments. (F) Δm157 or Δm157/m06/m152 MCMV infection of B6 or *Rag1*^-/-^ and (G) D8-KODO or ΔLy49-1 D8-KODO mice.

D8-KODO mice heterozygous for H2D^d^ (**Fig 2B**) displayed H2D^d^ expression at an MFI of nearly 50% of homozygous mice (**Fig 5C**), comparable to the loss seen during *in vitro* MCMV infection (**Fig 5A, B**). There was a similar level of H2D^d^ expression in (BALB/c x B6)F1 hybrid mice, confirming that physiologically relevant levels of H2D^d^ were being assessed, both *in vitro* and *in vivo*. Indeed, heterozygous D8 targets in uninfected homozygous D8-KODO mice were protected from clearance following a 3hr (**Fig 5D**) or 24hr (**Fig 5E**) *in vivo* cytotoxicity assay. In contrast, KODO targets were completely eliminated at both 24hrs (**Fig 5E**) and 48hrs (**Fig 4C**), and partially eliminated at 3hrs after injection (**Fig 5D**). During MCMV infection, there was enhanced clearance of both MHC-null and MHC-heterozygous targets at all times post-infection (**Fig 5D**), consistent with our previous study, where we identified a role for cytokines modulating missing-self rejection during MCMV infection [37]. While heterozygous target cell rejection never reached levels seen with MHC-null targets, NK cell-dependent clearance of missing-self targets can discriminate between normal levels and partial down-regulation of MHC molecules in the context of MCMV infection.

### Deletion of MCMV genes *m06* and *m152* prevents NK-dependent clearance while enhancing CD8^+^ T cell-dependent protection

To determine if *m06* and *m152* perturb NK-dependent viral resistance, we generated an independent set of Δm157-MCMV stocks in a virulent MCMV strain with mutations in both *m06* and *m152*, termed Δm157/Δm06/Δm152-MCMV. Δm157-MCMV and Δm157/Δm06/Δm152-MCMV showed similar rates of *in vitro* growth and viral titers, though Δm157-MCMV had a slight advantage (**Fig S6**). Regardless, infection of *Rag1*^−/−^ mice on the B6 background indicated that both viruses had comparable replication *in vivo* (**Fig 5F**). By contrast, Δm157/Δm06/Δm152-MCMV infection of B6 and ΔLy49-1 D8-KODO mice, otherwise susceptible to Δm157-MCMV (**Fig 1B, 2B**), resulted in a resistant phenotype (**Fig 5F, 5G**). D8-KODO mice were also resistant to Δm157/Δm06/Δm152-MCMV but resistance was not dependent on NK cells (**Fig 5G**), contrasting their NK cell-dependent resistance to Δm157-MCMV (**Fig 1C**). Instead, CD8 depletion significantly reversed resistance to Δm157/Δm06/Δm152-MCMV (**Fig 5G**), unlike the absence of an effect on resistance to Δm157-MCMV (**Fig 1C**).

## Discussion

Here, we clearly demonstrate that specific NK cell Ly49 inhibitory receptors have a critical MHC-restricted role in controlling viral infection *in vivo* (**Fig S7**). Their role is dependent on their specificity for self-MHC-I that influences their effects on NK cell licensing. Viral modulation of MHC-I was required because when MHC-I was no longer down-regulated, early resistance was due to CTLs instead of NK cells. Both the host (*Ly49a* and *Ly49g*) and the virus (*m06* and *m152*) encode multiple molecules involved in this resistance, a redundancy highlighting the ongoing arms race between the host and pathogen, and providing definitive support for the missing-self hypothesis.

In other strains of mice, there are likely differences in the NK cell response to MCMV due to Ly49 polymorphisms, particularly their specificities for MHC-I, and receptor repertoire, including subset distribution. Similarly, hosts may have different MHC-I alleles with varying capacities to license NK cells and susceptibilities to downregulation by viral MHC-I inhibitors. These factors likely account for differences in the MHC-restricted phenotypes described here. Moreover, there appears to be activation receptors, akin to Ly49H in B6 mice, which may dominate NK cell function if their ligands are expressed on infected cells, in which case licensed NK cells may play a secondary role. Finally, MCMV itself has evolved alleles of its ORFs that modulate these processes.

Our studies suggest that MCMV should encode molecules to specifically inhibit licensed NK cells. Indeed, MCMV-encoded m157 can engage Ly49 inhibitory receptors in mouse strains that do not have Ly49H activation receptor-equivalents; *e.g.* Ly49I in 129 mice [15] and Ly49C in BALB/c [20]. We predict that m157 inhibition of NK cell function will depend on whether these inhibitory receptors are in hosts with appropriate MHC-I alleles for licensing. Indeed, m157 effects on inhibiting NK cell control appear to be MHC-dependent [20], suggesting such potential interactions. Moreover, MCMV has 11 ORFs with predicted MHC-I folds [38], some of which have been verified by crystallographic studies, and thus may be similarly involved in modulating NK cells. Prior findings have suggested that upon downmodulation of MHC-I via m06 and m152, a third immunoevasin (m04) acts to stabilize certain MHC-I alleles on the surface of infected cells, leading to NK cell-dependent resistance, depending on additional activation receptors [19]. However, our studies do not show an apparent role for m04, indicating the MHC-I-restricted, inhibitory NK cell receptor-dependent anti-viral effects described here are fundamentally and mechanistically different from prior reports.

Beyond viral control, our studies also establish that Ly49 inhibitory receptors play a critical role in missing-self rejection, as previously predicted, based on *in vitro* observations and mice with global defects in MHC-I expression that have unlicensed NK cells. Here we clearly show that absence of specific inhibitory Ly49 receptors in a mouse expressing MHC-I results in unlicensed NK cells and an inability to perform missing-self rejection. It should be noted that prior studies of Ly49 specificities were often dependent on overexpression systems that may not reflect physiological interactions. Understanding of these ambiguities will be aided by the Ly49 KO mice described here as well as complementation by restoring expression of a single Ly49 receptor. Taken together, these data also provide definitive evidence that the inhibitory receptors are required for missing-self rejection *in vivo*.

As in viral control, NK cell effects on tumor control greatly rely on both NK cell activation and inhibitory receptor signaling [39]. MHC-I downmodulation during tumor growth to evade CTLs provides an attractive target for oncogenic control, yet the *in vivo* requirements for this potential critical function are poorly understood. Indeed, we recently showed that missing-self rejection in a mouse with inducible β2m deletion is markedly enhanced by inflammatory stimuli, such as MCMV infection, otherwise licensed NK cells can lose the licensed phenotype when profound MHC-I deletion occurs [37], consistent with results reported here. Therefore, our studies on NK cell control of viral infection, demonstrating of the complex *in vivo* interaction between inhibitory receptors and self-MHC-I, can serve as an analogous framework for considering how to modulate NK cells for controlling cancer.

## Methods

### Mice

C57BL/6Ncr (B6) and BALB/cAnNCr (BALB/c) mice were purchased from Charles River Laboratories. C57BL/10SnJ (B10; 000666), B10.D2/nSnJ (B10.D2; 000666), Prf1^-/-^ (002407) and RAG1^-/-^ (002216) strains were purchased from Jackson Laboratory. DAP10^-/-^DAP12^-/-^ mice were provided by T. Takai [40]. NKG2D^-/-^ mice were obtained from Bolan Polic (University of Rijeka, Croatia) [23]. Granzyme B KO mice were obtained from T. Ley (Washington University, St. Louis) [41]. H2K^b−/−^ H2D^b−/−^ (KODO) mice were purchased from Taconic Farms. Triple knockout (TKO) mice which are H2K^b-/-^ H2D^b-/-^ and lack β2m were obtained from Dr. Ted Hansen (Washington University, St. Louis). D8 transgenic mice expressing an H2D^d^ genomic construct [42] were provided by D. Marguiles (National Institute of Allergy and Infectious Diseases, Bethesda, MD). D8-KODO mice were generated by crossing D8-transgenic mice to KODO mice. Generation and characterization of the ITIM-mutant AYF mice on the H2^d^ background have been previously described [31]. KLRA7^em1(IMPC)J^ (Ly49G KO) mice on the C57BL/6NJ background were purchased from the Jackson Laboratory (027444); this allele was generated at the Jackson Laboratory by CRISPR-Cas9 injection of Cas9 RNA and 3 gRNAs: TCTTGTACTTGTGCATAACC, CAGTCCTCACTAGTTTCTGC and GACATGGACTGACCAAATT resulting in a 241 bp deletion beginning in 5-prime upstream sequence and ending within exon 1. Additional strains of mice generated through CRISPR-Cas9 are described below. All mice, with the exception of the RAG^-/-^ strain, used in these studies were initially obtained or generated on the B6 genetic background and later backcrossed to the KODO and then D8-KODO background (H2D^d^ MHC). B6 mice used in experiments were obtained directly from Charles River Laboratories. All other experimental and control mice were bred in-house at Washington University. Mice were 8–14-wk old at the start of experiments unless otherwise stated. Male and/or female mice were used in individual experiments without blinding or randomization. This study was carried out in strict accordance with the recommendations in the Guide for the Care and Use of Laboratory Animals of the National Institutes of Health. The studies were approved by the Animal Studies Committee at Washington University School of Medicine under animal protocol 20180293.

### Development of CRISPR-Cas9 modified knock-out mice

While NCBI BLAST (http://blast.ncbi.nlm.nih.gov/) was initially used to assess sequence similarity of potential sgRNAs, GT-Scan (https://gt-scan.csiro.au/gt-scan) [43] and CCTop (https://crispr.cos.uni-heidelberg.de/) [44] were primarily used to confirm the correct and specific targeting our sgRNA designs. The sequences of synthetic guide RNAs (sgRNA) and the strains of mice generated are shown in **Table S1**. Cas9 mRNA and sgRNA synthesis, RNA micro-injection into zygotes is identical to what we described earlier [45]. Specifically, we used 20ng of each guide when multiple guides were indicated, and 100ng of Cas9 mRNA. Pups were screened for *Ly49* gene disruption at birth by PCR. If multiple Ly49 alleles were present, they were separated by one backcross to B6 and re-screened by PCR. All mice with in-frame insertions or deletions (indels) were excluded from further analysis. Multiple lines with out-of-frame indels were chosen for downstream characterization. These genetic lesions were verified by absence of PCR amplicons for each of the indicated *Ly49* genes in homozygous mice, and specific PCR amplicons for the deleted genomic regions, that were sequenced to determine the exact breakpoints (**Fig S1** and **Fig S2**). Finally, Ly49 protein expression was examined, where antibodies were available, at 8 weeks in homozygous mice following at least one additional backcross to parental B6. Since all sgRNAs targeted the second exon of *Ly49* genes, we focused on this exon to specifically interrogate on-target and off-target analysis. As shown in **Fig S1**, all exons and the sgRNA with on-target and off-target potential are depicted. Sanger sequence traces of homozygous mice are shown below the schematic. Flow analysis across genes expressed within the NKC in D8-KODO mice is presented as **Fig S2**. Specifically, we generated Ly49C/I-double-deficient mice (two lines; one used in experiments), *Ly49m*-deficient mice (four lines used in experiments), Ly49A-deficient mice (two lines used in experiments), Ly49A/G-double-deficient mice (two lines used in experiments), and the ΔLy49-1 D8-KODO multi-deficient mice (one line used in experiments). The Ly49G-deficient mice were purchased from Jackson Laboratory, however, full on-target, off-target (**Fig S1**) and flow cytometric (**Fig S2**) analyses was performed by our laboratory. Importantly, no major impact on maturation was observed in any of these mice.

### Development of CRISPR-Cas9 modified Ly49A knockin mice

The NKp46 (*Ncr1*) locus was chosen for insertion of the *Ly49A* cDNA (**Fig S3**). The donor construct was designed to replace the stop codon of NKp46 (while maintaining the 3-prime untranslated region) with a P2A peptide-cleavage site upstream of the *Ly49A* cDNA obtained from reverse transcription of Ly49A-expressing B6 NK cells. The result is that Ly49A expression would be restricted to all NK cells. Given that the *Ly49A* knockin is located on chromosome 7, we were able to cross this mouse with the ΔLy49-1 mouse with Ly49 deletions on chromosome 6 without linkage restrictions. GT-Scan [43] and CCTop [44] were primarily used to confirm the correct and specific targeting our sgRNA to the NKp46 locus. The sequences of guide RNAs are shown in **Table S1**. Cas9 mRNA and sgRNA synthesis, RNA and donor DNA micro-injection is identical to what we described earlier [45]. Specifically, we used 10ng of donor DNA, 20ng of each guide (2 total), and 100ng of Cas9 mRNA. Pups were screened for the donor DNA by PCR soon after birth and maintained as heterozygotes for flow-based confirmation at 8 weeks. After two rounds of B6 backcrossing, the mice were bred to ΔLy49-1 D8-KODO. Mice homozygous for the NKC deletion but heterozygous for the Ly49A knockin were screened for expression of Ly49A (**Fig S3**). Subsequently, various NKC surface molecules were analyzed by flow cytometry (**Fig S2**). Again, no major impact on maturation was observed.

### Development of CRISPR-Cas9 modified MCMV

Δm157-MCMV and GFP-expressing MCMV were modified to knock-out various viral ORFs using CRISPR-Cas9 editing to obviate the need for bacterial artificial chromosome modification. The GFP-expressing MCMV was a generous gift from S.C. Henry and J. Hamilton (Duke University, Durham, NC, USA) [46] and previously described by our lab to harbor a non-functional *m157* gene [47]. The GFP-expressing MCMV and CRISPR-derived progeny were used in *in vitro* studies while the Δm157-MCMV derived from WT1 [48] and its CRISPR-modified progeny were used in the in vivo analyses. CCTop [44] was used to identify and confirm specific targeting our sgRNAs to the MCMV ORFs. The stand-alone version of CCTop was loaded with the MCMV genome (GenBank Accession GU305914.1) to effectively eliminate off-target cleavage potential. The table of all possible guide site for MCMV is provided (**Table S2**). The pX330 vector [49, 50] was obtained from Addgene and modified by replacing the *Sbf*I-*Psi*I site with the Neo marker (*Sbf*I-*Hinc*II fragment) from pCDNA3. The oligonucleotide duplexes containing single sgRNAs specific for MCMV *m06* or *m152* (**Table S3**) were cloned into this vector that was subsequently transfected into SV40-immortalized B6 MEFs with G418 selection. Parental virus was used to infect these sgRNA and Cas9-expressing MEFs at a low MOI (0.1). Once confluent lysis was observed, viral supernatants were used to reinfect B6 MEFs, individual plaques were picked, and sequence variants were confirmed (**Fig S5**). For double-deficient MCMV ORF knockouts, a pure stock of virus (e.g. *m06* knockout) was used to infect MEFs expressing Cas9 and the alternate sgRNA of interest (e.g. *m152*). For *in vivo* analysis, plaque-purified virus was used to generate salivary gland passaged stocks in BALB/c mice as described above. Salivary gland-derived MCMV lacking m06, m152, and m157 was compared to MCMV lacking m157 in a multi-step growth curve on NIH 3T12 (ATCC CCL-164) fibroblasts (MOI = 0.1). Viral genome copies quantified from cell lysates and culture supernatants, as previously described [51], demonstrated that replication of the triple knock-out virus was unimpaired yet delayed slightly in overall growth (**Fig S6**).

### Viral infection and quantification

The salivary gland propagated MCMV stocks were generated from purified and sequenced clones. Virus was inoculated via the intraperitoneal (IP) route in a total volume of 200μl PBS at a dose of 40,000 plaque forming units (pfu) per mouse for WT-MCMV and 20,000 pfu per mouse for m157-deficient MCMV [51] (Δm157-MCMV) and Δm157/m06/m152-MCMV for determination of splenic titers at day 4. Viral titers from infected spleens were quantified as a ratio of MCMV IE DNA to host beta-actin DNA using real-time PCR and Taqman probes as previously described [51]. When shown, individual data points represent a single mouse. Survival analysis endpoints were determined as either death or >20% weight loss from starting weights, using 300,000 pfu of Δm157-MCMV per mouse. To examine H2D^d^ MHC-I downmodulation, SV40-immortalized D8-KODO MEFs were infected with GFP-expressing MCMV at an MOI of 1 for 24 hours and subsequently released from plates with Versene (Thermo Fisher, Waltham, MA) prior to antibody staining and flow cytometric analysis.

### Antibody depletions

Purified 3D10 (α-Ly49H), PK136 (α-NK1.1), JR9 (α-Ly49A), 4E5 (α-Ly49D), LGL-1 (α-Ly49G) and YTS-169 (α-CD8α) were obtained from hybridomas purified by the Rheumatic Diseases Core Center Protein Purification and Production Facility (Washington University). Antibodies were injected IP at a dose of 200μg per mouse 48 hours prior to infection. >98% depletion was confirmed via flow cytometry in a subset of treated mice. Injection of PBS alone was used as a control where indicated.

### Antibodies and Flow Cytometry

The following antibodies and reagents were purchased from eBioscience: anti-CD3e (145-2C11), anti-CD19 (eBio1D3), anti-NK1.1 (PK136), anti-NKp46 (29A1.4), anti-CD27 (LG.7F9), anti-CD11b (M1/70), anti-Ly49D (eBio4E5), anti-Ly49E/F (CM4), anti-Ly49G2 (eBio4D11), anti-Ly49H (3D10), anti-Ly49I (YLI-90), anti-CD94 (18d3), anti-NKG2A^B6^ (16a11), anti–IFN-γ (XMG1.2), anti-CD122 (5H4), anti-CD127 (SB/199), anti-CD69 (H1.2F3), anti-2B4 (2B4), anti-NKG2D (CX5) and Fixable Viability Dye eFluor 506. The following antibodies and reagents were purchased from BD Biosciences: anti-Ly49F (HBF-719), anti-Ly49G2 (4D11), and streptavidin (SA)-phycoerythrin. The following antibodies and reagents were purchased from BioLegend: anti-NK1.1 (PK136), anti-H2D^d^ (34-2-12) and SA-allophycocyanin. Anti-Ly49I (YLI-90) was purchased from Abcam. Anti-Ly49A (JR9) was purified in our laboratory from hybridoma supernatants and subsequently conjugated to biotin or FITC. The JR9 hybridoma was generously provided by Jacques Roland (Pasteur Institute, Paris, France) [52]. Anti-Ly49C (4LO33) [53] was purified in our laboratory from hybridoma supernatants and subsequently conjugated to biotin. The 4LO hybridoma was generously provided by Suzanne Lemieux (Institut National de la Recherche Scientifique-Institut Armand-Frappier, Laval, Quebec, Canada). Anti-NK1.1 (PK136) was purified in our laboratory from hybridoma supernatants. The PK136 hybridoma was purchased from American Type Culture Collection. Fc receptor blocking was performed with 2.4G2 (anti-FcγRII/III) hybridoma (American Type Culture Collection) culture supernatants. Surface staining was performed on ice in staining buffer (1% BSA and 0.01% NaN3 in PBS). Samples were collected using a FACSCanto (BD Biosciences), and data were analyzed using FlowJo (TreeStar).

### *In vivo* cytotoxicity

Mice used for donor splenocytes in *in vivo* cytotoxicity assays were 8–12 weeks old at the time of transfer. WT-MCMV, where indicated, was inoculated 3 days prior at a dose of 10,000 pfu per mouse [51]. Donor splenocytes were harvested and labeled *in vitro* with 2.5 µM CFSE (Life Technologies) and 5 or 0.2 µM CellTrace violet (CT violet; Thermo Fisher). Recipient mice were injected intravenously with 2 × 10^6^ of each donor. Spleens from recipient mice were harvested 3 (**Fig 5B**), 24 (**Fig 5C**) or 48 (**Fig 4C**, **Fig S4**) hours after transfer of donor cells. NK cell-specific rejection was calculated by gating on transferred CFSE-positive cells and excluding dead cells by forward scatter and side scatter. Rejection was quantified as Killing Percentage = [1 − (Target/Control)/(Target/Control)Average(NK depleted)] × 100, where the target was the MHC-deficient (H2D^d-/-^)or heterozygous (H2D^d+/-^) donor cell and the control was a H2D^d+/+^ donor cell. The ratio of target to control cells was normalized to the average ratio recovered from NK cell-depleted mice to calculate rejection by NK cells.

### NK cell in vitro stimulation and intracellular cytokine staining

Splenocytes were stimulated with anti-NK1.1 (PK136) as previously described [11]. Briefly, 24-well culture plates were coated with 500 μL of purified PK136 (1 μg/mL). Plates were washed with PBS, and 5 × 10^6^ splenocytes were then added to each well in 500 μL of R10 (RPMI 1640 supplemented with 10% fetal bovine serum) media. Splenocytes were stimulated in parallel with 0.5 μg/mL PMA (Sigma-Aldrich) and 4 μg/mL ionomycin (Sigma-Aldrich) and incubated at 37 °C and in 5% CO2 for a total of 7 h. Brefeldin A (GolgiPlug; BD Biosciences) was added to the cells after 1 h. After staining surface antigens, cells were fixed and permeabilized (Cytofix/Cytoperm; BD Biosciences) followed by staining for IFN-γ. NK cells were gated as viable CD3^−^ CD19^−^ NKp46^+^ lymphocytes.

### Sequence analysis

PCR amplicons were amplified with the Phusion high-fidelity DNA polymerase (NEB, Ipswich, MA) using manufacturer recommended cycling conditions, column purified (Macherey-Nagel, Bethlehem, PA) and were sequenced on an ABI 3730 at Genewiz, Inc (South Plainfield, NJ, USA). The resulting chromatograms were aligned using SnapGene software (GSL Biotech, Chicago, IL) and the relevant reference sequences for MCMV (GenBank Accession GU305914.1) or the C57BL/6 (GRCm38/mm10). All oligonucleotides were synthesized by IDT (Coralville, IA).

### Statistical analysis

Statistical analysis was performed using Prism (GraphPad software). Unpaired, two-tailed Student’s t-tests were used to determine statistically significant differences between experimental groups. For all t-tests, the number of degrees of freedom (df) equals the total sample size minus 2. Error bars in all figures represent the standard error of the mean (SEM). \*\*\*\**p* < .0001, \*\*\**p* < .001, \*\**p* < .01, \**p* < .05, ns = not significant.

### Data and biological material availability

The data that support the findings of this study are available from the corresponding authors on reasonable request. Novel cell lines, viral constructs, and mouse strains are available from the author’s laboratory (W.M.Y.) upon request.

## Acknowledgments

We thank J. Michael White (Transgenic, Knockout, and Micro-Injection Core at Washington University) for CRISPR/Cas9 injections and Andrew Cao, Trenton J (TJ) Dawson, and Darryl Higuchi for technical assistance. Experimental support was provided by the Protein Production and Purification Core Facility. This work was supported by National Institutes of Health grants R01AI129545 and R01AI131680 (to W.M.Y) and K08-AI104991 (to B.A.P.)

## Author contributions

B.A.P and W.M.Y designed the research. B.A.P., M.D.B., S.J.P., L.Y., D.L.B., and J.P.-L. performed the experiments. B.A.P and W.M.Y analyzed the data and wrote the paper.

The authors declare no competing financial interests.

## Supplementary Information

### Supplementary Figure Legends

**Fig S1. Characterization of on-target and off-target CRISPR deletions targeting specific *Ly49* genes.** Genomic DNA was isolated from back-crossed H2D^d^ CRISPR-Cas9 modified mice used in these studies. The specific CRISPR-targeted exons (those with homology to the ITIM) were Sanger sequenced as representative of the most likely position of off-target effects and to confirm and characterize the frameshift in on-target variants. The primers used for PCR and subsequent sequence analysis are shown in **Table S4**. (A-P) Each panel depicts the reference sequence (above), the region of the exon targeted (in blue), the on-target and off-target guide sites (in purple), and the Sanger alignments (below) for the mice indicated. Mice are designated as to which strains (in parentheses) were analyzed. B6 is the wild-type genome for comparison. Red boxes indicate indels (inserted nucleotides are shown, while deleted bases are represented as a dash). Sequence analysis was confirmed in both the forward and reverse directions, however, for clarity only one direction is shown. Sequences assessed in each panel are as follows: (A) *Klra1/Ly49a*, (B) *Klra2/Ly49b*, (C) *Klra3/Ly49c*, (D) *Klra4/Ly49d*, (E) *Klra5/Ly49e*, (F) *Klra6/Ly49f*, (G) *Klra7/Ly49g*, (H) *Klra8/Ly49h*, (I) *Klra9/Ly49i*, (J) *Klra10/Ly49j*, (K) *Klra11-ps/Ly49k*, (L) *Klra13-ps/Ly49m*, (M) *Klra14/Ly49n*, (N) *Klra17/Ly49q*, (O) *Gm6548*, and (P) *Gm15854/Ly49x*. (Q) Nucleotide level analysis of the ΔLy49-1 deletions for the haplotype corresponding to Fig 2A at the *Ly49g-Ly49a* and *Ly49n-Ly49k* junctions.

**Fig S2. Flow cytometric characterization of CRISPR-Cas9 modified mice.** Offset flow histograms are shown for mice on the D8-KODO (H2D^d^) MHC background with the indicated CRISPR-Cas9 modifications. Single-cell suspensions of mouse splenocytes were gated for expression of specific proteins with loci near or in the NKC. (A-O) Shown is a representative plot of NK1.1^+^/NKp46^+^/CD19^-^/CD3^-^ lymphocytes. Protein levels assessed in each panel are as follows: (A) Ly49A, (B) Ly49C, (C) Ly49D, (D) Ly49EF, (E) Ly49F, (F) Ly49G, (G) Ly49H, (H) Ly49I, (I) CDD94, (J) NKG2A, (K) NKG2D, (L) CD69, (M) CD122, (N) CD127 and (O) 2B4. The flow cytometric analyses were repeated twice with 3 mice per group. Below the plots, quantification of the flow data as frequencies (A-J) or MFI (K-O) from one of two representative experiments is provided. Statistical analysis was performed with a Student’s *t*-Test with D8-KODO expression frequency used as a comparator. (P) Representative maturation plots are shown for all experimental mice with the summation of a representative experiment provided at the bottom. No significant deviations were noted. Analyses were repeated twice with 3 mice per group. (Q) Quantification of NK cell frequencies gated as CD3^-^CD19^-^NK1.1^+^ lymphocytes. Statistical analysis was performed with a Student’s *t*-Test with D8-KODO expression frequency used as a comparator as above. (R) Flow cytometric analysis of mice heterozygous for the ΔLy49-1 D8-KODO KO (F1 hybrids of KODO mice with intact *Ly49*s and ΔLy49-1 D8-KODO mice from **Fig2B**) compared with KO and WT mice. Experiments were performed twice with 5-7 mice per group, with the exception that WT mice were assessed only once in this panel with values consistent with panels above. (S) Flow cytometry gating strategy.

**Fig S3. Strategy and characterization of the NKp46-Ly49A knockin.**

(A) CRISPR knockin strategy for Ly49A expression on NKp46-expressing cells. 2 CRISPR guides, T1 and T3 (**Table S1**) were injected into B6 zygotes along with Cas9 mRNA and a donor DNA construct engineered to express the *Ly49A* cDNA following a P2A cleavage site such that the Ly49A expression would be proportional to NKp46 (*Ncr1*) transcript levels. (B) The final NKp46 TGA stop codon and several downstream bases (yellow highlighted region) in the coding region of exon 7 (blue text) were removed and replaced with a P2A site (purple text), Ly49A cDNA (red text) and finally the remainder of the NKp46 3’ UTR sequence (green text). This donor vector with NKp46 homologous arms was co-injected with sgRNA for *Ncr1* and Cas9 into B6 zygotes and positive knockin mice were selected at birth by PCR analysis and confirmed at 6 weeks of age by flow cytometry. (C) One Ly49A KI line was crossed to the ΔLy49-1 D8-KODO mice (Fig 2A) heterozygous for one knockin allele and confirmed by flow cytometry. Lymphocytes for splenic single cell suspensions were assessed for both NK1.1 and Ly49A expression. Approximately 18% of NK cells from WT (D8-KODO) mice expressed Ly49A, while the ΔLy49-1 D8-KODO mice lacked all Ly49A expression unless expressing the Ly49A from the knockin construct (nearly 100% expression). As shown, Ly49A expression was restricted to NK1.1^+^ splenocytes.

**Fig S4. In vivo cytotoxicity in Ly49AG KO and Ly49AYF mice.**

*In vivo* cytotoxicity of MHC-I deficient (KODO) or sufficient (D8-KODO) splenocytes following differential labelling with Celltrace Violet and flow cytometric analysis recovered from spleens at 2 days post-injection. (A) Flow cytometric analysis of input cells is shown (prior to injection). The experimental design is also depicted with the timing of antibody depletion relative to injection and harvest. (B) Representative histogram plots of the mice indicated in each row and treatments by column. The peaks are identified by the input plot shown in (A). (C) Quantification was performed as previously described (Fig 4C). Data are representative of two independent experiments with three recipient mice per group that received the same mix of donor cells in each experiment. Standard error of the mean is shown; statistical analysis performed with a Student’s *t*-Test. \**p* < .05, \*\*\*\**p* < .0001.

**Fig S5. Sanger sequence analysis of m06 and m152 in CRISPR-modified MCMV strains.**

Viral DNA was isolated from *in vitro* cell cultures infected with CRISPR-Cas9 modified MCMV used in these studies. The specific CRISPR-targeted ORFs, (A) *m06* and (B) *m152*, were Sanger sequenced to confirm and characterize the frameshift in on-target variants. The primers used for PCR and subsequent sequence analysis are shown in **Table S5**. Each panel depicts the reference sequence above with the region of the ORF expanded for analysis. The Sanger alignments below for the viruses indicated are designated as to which strains (*left* of trace) were analyzed. Δm157-MCMV is the wild-type genome at these ORFs, for comparison. Red boxes indicate where inserted nucleotides were identified. Sequence analysis was confirmed in both the forward and reverse direction, however, only one direction is shown for clarity.

**Fig S6. Multi-step growth curve**

Multi-step *in vitro* growth kinetics of two strains of MCMV, as indicated. Cells and supernatants were harvested at the indicated days and quantified by real-time PCR. Data is a representative of two independent experiments with each data point performed in triplicate. All timepoints were shown to significantly different in terms of viral load (*p* < .0001); statistical analysis performed with a Student’s *t*-Test.

**Fig S7. Model for MHC-restricted and Ly49 receptor-dependent MCMV resistance.**

(A) Activation receptor Ly49H-dependent induces killing of m157-expressing MCMV-infected cells. (B) Deficiency of m157 prevents rejection of MCMV in B6 mice expressing H2K^b^ (H2D^b^ is present but not shown for clarity). (C) Granzyme and perforin expression on licensed NK cells are required for control of m157-deficient virus in H2D^d^-expressing mice. (D) MHC-I downmodulation by MCMV prevents recognition of missing-self rejection in mice lacking Ly49A and Ly49G with unlicensed NK cells. (E) MCMV ORFs, *m06* and *m152*, are essential for the NK-specific protective response and the loss of these molecules results in the emergence of T cell-dependent resistance and concomitant loss of NK cell-dependent control by licensed NK cells.

