## Supplemental Figures for "Major histocompatibility complex class I-restricted protection against murine cytomegalovirus requires missing-self recognition by the natural killer cell inhibitory Ly49 receptors"

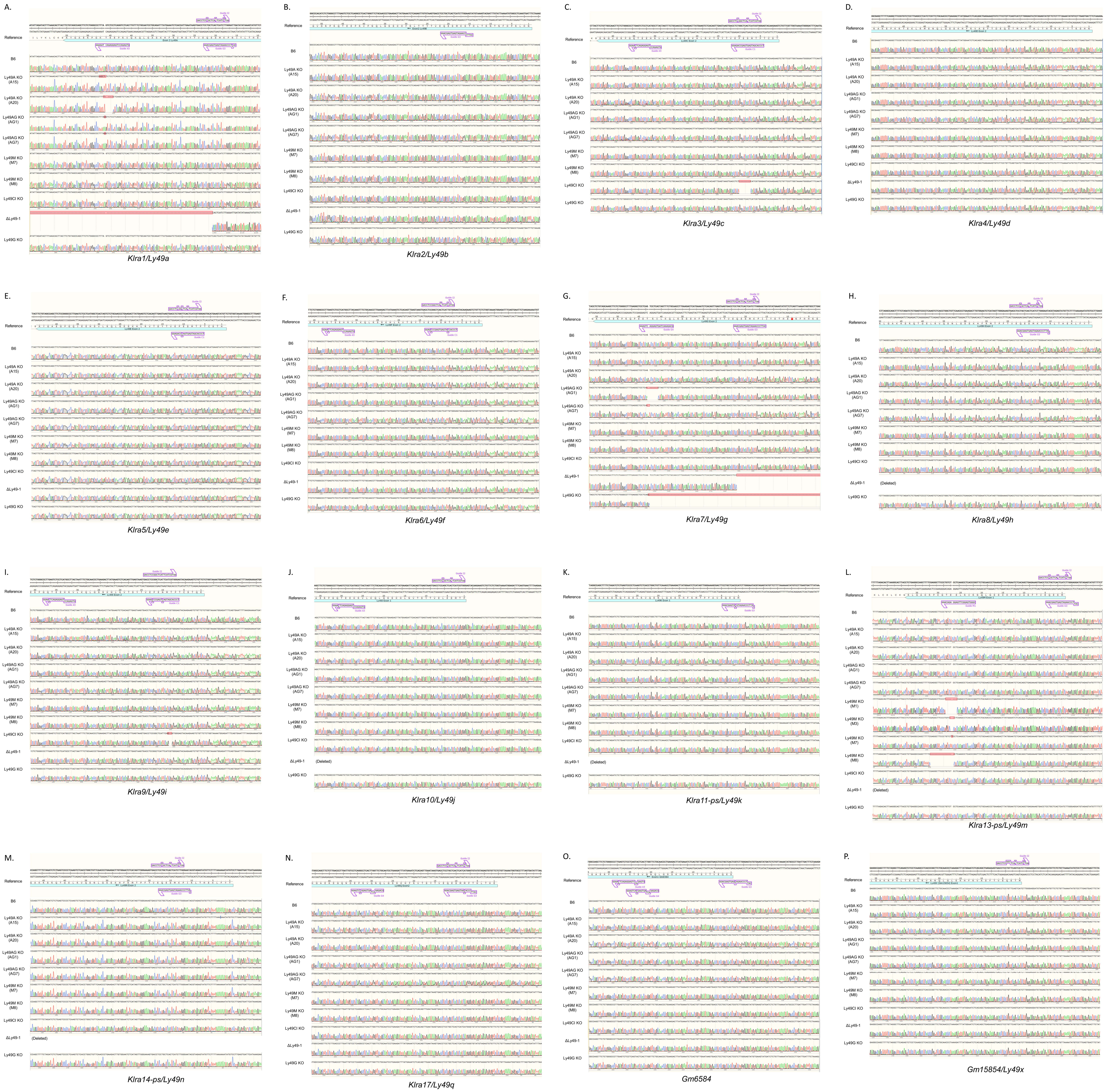


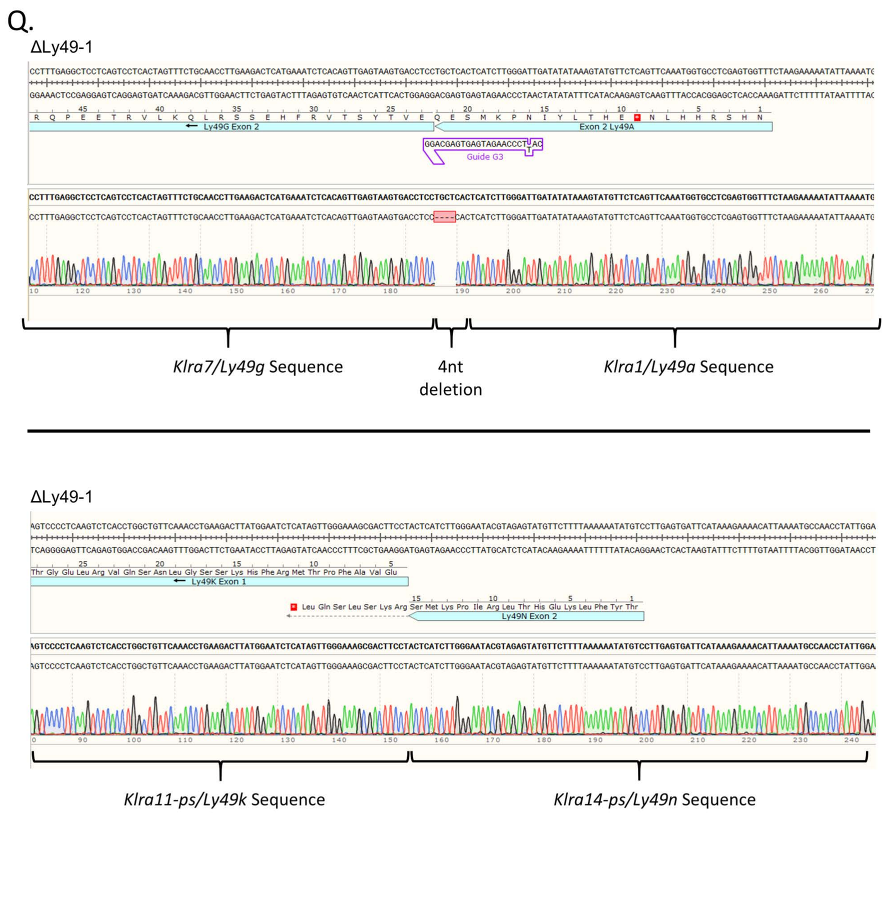


**Fig S1. Characterization of on-target and off-target CRISPR deletions targeting specific *Ly49* genes.**
Genomic DNA was isolated from back-crossed H2D$^d$ CRISPR-Cas9 modified mice used in these studies. The specific CRISPR-targeted exons (those with homology to the ITIM) were Sanger sequenced as representative of the most likely position of off-target effects and to confirm and characterize the frameshift in on-target variants. The primers used for PCR and subsequent sequence analysis are shown in **Table S4**. (A-P) Each panel depicts the reference sequence (above), the region of the exon targeted (in blue), the on-target and off-target guide sites (in purple), and the Sanger alignments (below) for the mice indicated. Mice are designated as to which strains (in parentheses) were analyzed. B6 is the wild-type genome for comparison. Red boxes indicate indels (inserted nucleotides are shown, while deleted bases are represented as a dash). Sequence analysis was confirmed in both the forward and reverse directions, however, for clarity only one direction is shown. Sequences assessed in each panel are as follows: (A) *Klra1/Ly49a*, (B) *Klra2/Ly49b*, (C) *Klra3/Ly49c*, (D) *Klra4/Ly49d*, (E) *Klra5/Ly49e*, (F) *Klra6/Ly49f*, (G) *Klra7/Ly49g*, (H) *Klra8/Ly49h*, (I) *Klra9/Ly49i*, (J) *Klra10/Ly49j*, (K) *Klra11-ps/Ly49k*, (L) *Klra13-ps/Ly49m*, (M) *Klra14/Ly49n*, (N) *Klra17/Ly49q*, (O) *Gm6548*, and (P) *Gm15854/Ly49x*. (Q) Nucleotide level analysis of the ΔLy49-1 deletions for the haplotype corresponding to **Fig 2A** at the *Ly49g-Ly49a* and *Ly49n-Ly49k* junctions.

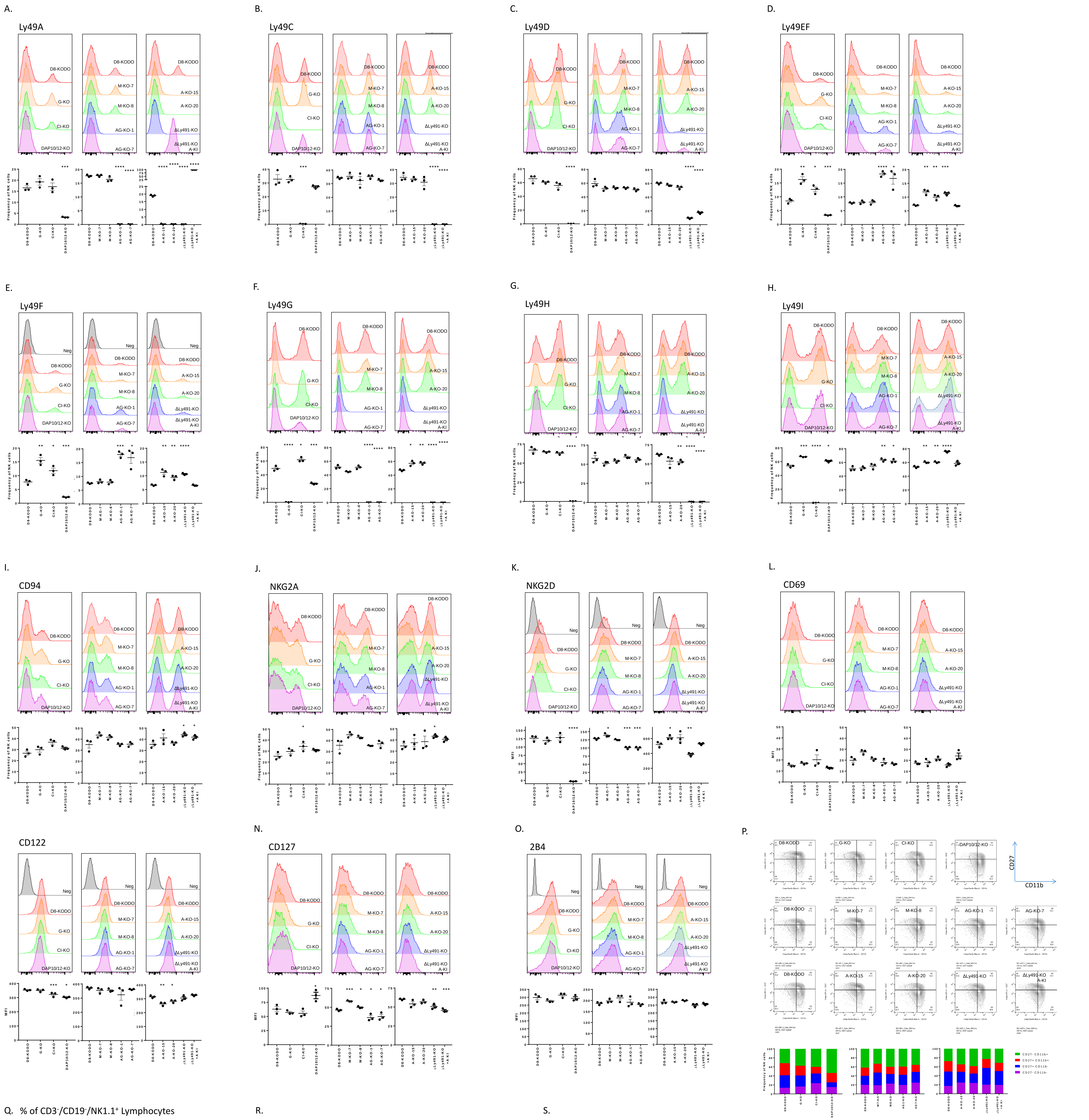


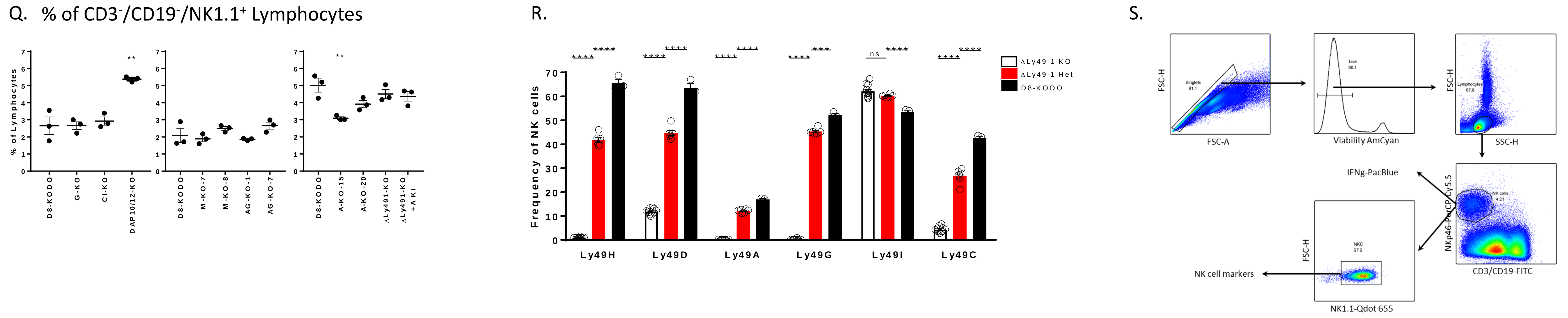


**Fig S2. Flow cytometric characterization of CRISPR-Cas9 modified mice.**
Offset flow histograms are shown for mice on the D8-KODO (H2D$^d$) MHC background with the indicated CRISPR-Cas9 modifications. Single-cell suspensions of mouse splenocytes were gated for expression of specific proteins with loci near or in the NKC. (A-O) Shown is a representative plot of NK1.1$^+$/NKp46$^+$/CD19$^-$/CD3$^-$ lymphocytes. Protein levels assessed in each panel are as follows: (A) Ly49A, (B) Ly49C, (C) Ly49D, (D) Ly49EF, (E) Ly49F, (F) Ly49G, (G) Ly49H, (H) Ly49I, (I) CDD94, (J) NKG2A, (K) NKG2D, (L) CD69, (M) CD122, (N) CD127 and (O) 2B4. The flow cytometric analyses were repeated twice with 3 mice per group. Below the plots, quantification of the flow data as frequencies (A-J) or MFI (K-O) from one of two representative experiments is provided. Statistical analysis was performed with a Student's *t*-Test with D8-KODO expression frequency used as a comparator. (P) Representative maturation plots are shown for all experimental mice with the summation of a representative experiment provided at the bottom. No significant deviations were noted. Analyses were repeated twice with 3 mice per group. (Q) Quantification of NK cell frequencies gated as CD3$^-$CD19$^-$NK1.1$^+$ lymphocytes. Statistical analysis was performed with a Student's *t*-Test with D8-KODO expression frequency used as a comparator as above. (R) Flow cytometric analysis of mice heterozygous for the ΔLy49-1 D8-KODO KO ($F_1$ hybrids of KODO mice with intact *Ly49*s and ΔLy49-1 D8-KODO mice from **Fig2B**) compared with KO and WT mice. Experiments were performed twice with 5-7 mice per group, with the exception that WT mice were assessed only once in this panel with values consistent with panels above. (S) Flow cytometry gating strategy.

**A.**

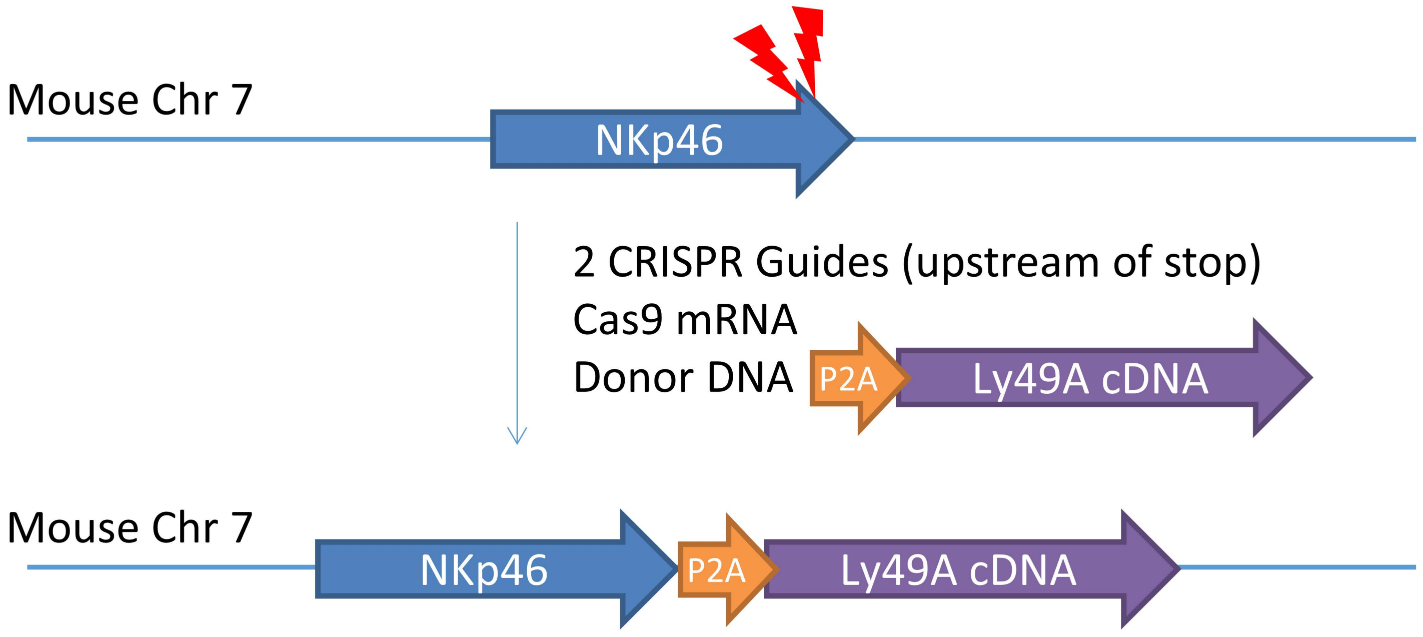


**B.**

Segment of chr7:4344460-4345184 (GRCm38/mm10)

Exon 7 coding region of Ncr1 / 3' UTR of Ncr1

Replaced with P2A-Ly49A cDNA

actccgccttctgggatcacacaacccagaatctcattcgaattggtctggcatgcataatcctgataactctagtatggcttttgactgaagactg
gctcagcaagaggaaagatcatgaagaggccaacagattaacaaattggaatgcaggagaagatggagaatgcagcattactttgaaga
ggaacaaagaaatgcaatatctatgatggaactgaaggcaactcctggggcattgtgagctctatgctgtaccctggataaagaaagagatct
tttcagaaatgaaggggtaggagtgttaccaagtttggcaaagcattctaaaggacactgtgagaggaagcctctaacagacctctctgtctct
gctaagctttaagtactttaacgcttatttactgcaagttatacttttgatcactctggtctttcctagaaataacttcgttccaatgtgatggccatggga
atatttccttcctactttaggtcttatcgtcatttctttttctaattttatttattttatgttgccgggaattgaactcaagttgtctggaggagcaacaagtgct
cttaactacaaaccatatctccagctccatcctcaatattttatattgtgtgtgcatgtctgtaggatgcatgtgtgtgcacacatatgtacatatattct
atgccattaaaatgaaggtcagagaacacctctgtgaagtca

ggatctggagcaacaaacttctcactactcaaacaagcaggtgacgtggaggagaatcccgggcctatgagtgagcaggaggtcacttattc
aatggtgagatttcataaatctgcaggattgcagaaacaagtgagacctgaggagactaaagggcccagagaagctggctacagaaggtg
ttcattccactggaagttcattgtgatagctcttggcatcttctgtttccttcttctggtagctgtttcagtgttggcaataaaaatttttcagtatgatcaac
aaaaaaaactgcaggaatttctaaaccaccacaataactgcagcaacatgcaaagtgacatcaacttgaaggatgaaatgctgaaaaata
agtctatagagtgtgatcttctggaatccctcaacagggatcagaacagattgtataataaaaccaagactgttttagattccttacagcacaca
ggcagaggtgataaagtatactggttctgctatggtatgaaatgttattatttcgtcatggacagaaaaacatggagtggatgtaaacagacctg
ccagagttccagtttatcccttctgaagatagatgatgaggatgaactgaagttccttcagctcgtggttccttcagacagttgctgggttggattgtc
atatgataataagaaaaaagattgggcatggattgacaatcgcccatctaaacttgccttgaacacaaggaaatacaatataagagatgggg
gatgtatgttgttatctaaaacaagactagacaatggtaactgtgatcaagtattcatctgtatttgtgggaagagactggataaattccctcattga

**C.**

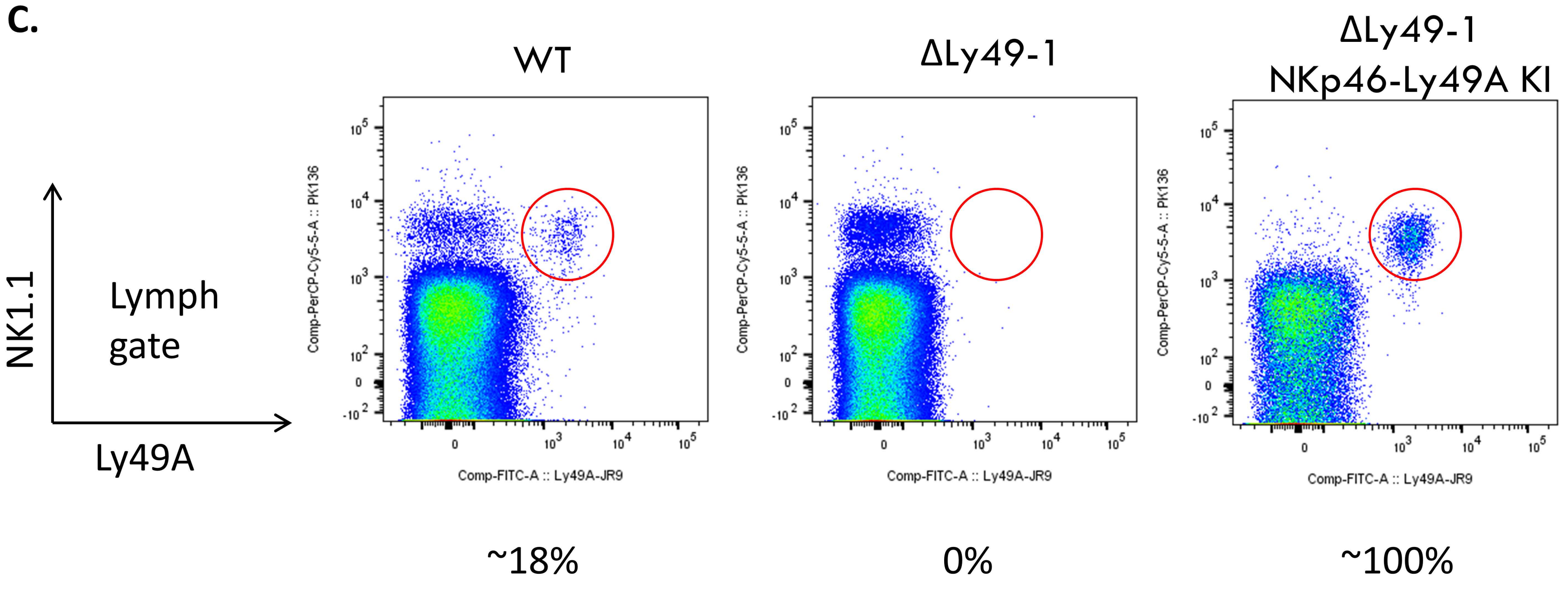


**Fig S3. Strategy and characterization of the NKp46-Ly49A knockin.**
(A) CRISPR knockin strategy for Ly49A expression on NKp46-expressing cells. 2 CRISPR guides, T1 and T3 (**Table S1**) were injected into B6 zygotes along with Cas9 mRNA and a donor DNA construct engineered to express the *Ly49A* cDNA following a P2A cleavage site such that the Ly49A expression would be proportional to NKp46 (*Ncr1*) transcript levels. (B) The final NKp46 TGA stop codon and several downstream bases (yellow highlighted region) in the coding region of exon 7 (blue text) were removed and replaced with a P2A site (purple text), Ly49A cDNA (red text) and finally the remainder of the NKp46 3' UTR sequence (green text). This donor vector with NKp46 homologous arms was co-injected with sgRNA for *Ncr1* and Cas9 into B6 zygotes and positive knockin mice were selected at birth by PCR analysis and confirmed at 6 weeks of age by flow cytometry. (C) One Ly49A KI line was crossed to the ΔLy49-1 D8-KODO mice (**Fig 2A**) heterozygous for one knockin allele and confirmed by flow cytometry. Lymphocytes for splenic single cell suspensions were assessed for both NK1.1 and Ly49A expression. Approximately 18% of NK cells from WT (D8-KODO) mice expressed Ly49A, while the ΔLy49-1 D8-KODO mice lacked all Ly49A expression unless expressing the Ly49A from the knockin construct (nearly 100% expression). As shown, Ly49A expression was restricted to $NK1.1^+$ splenocytes.

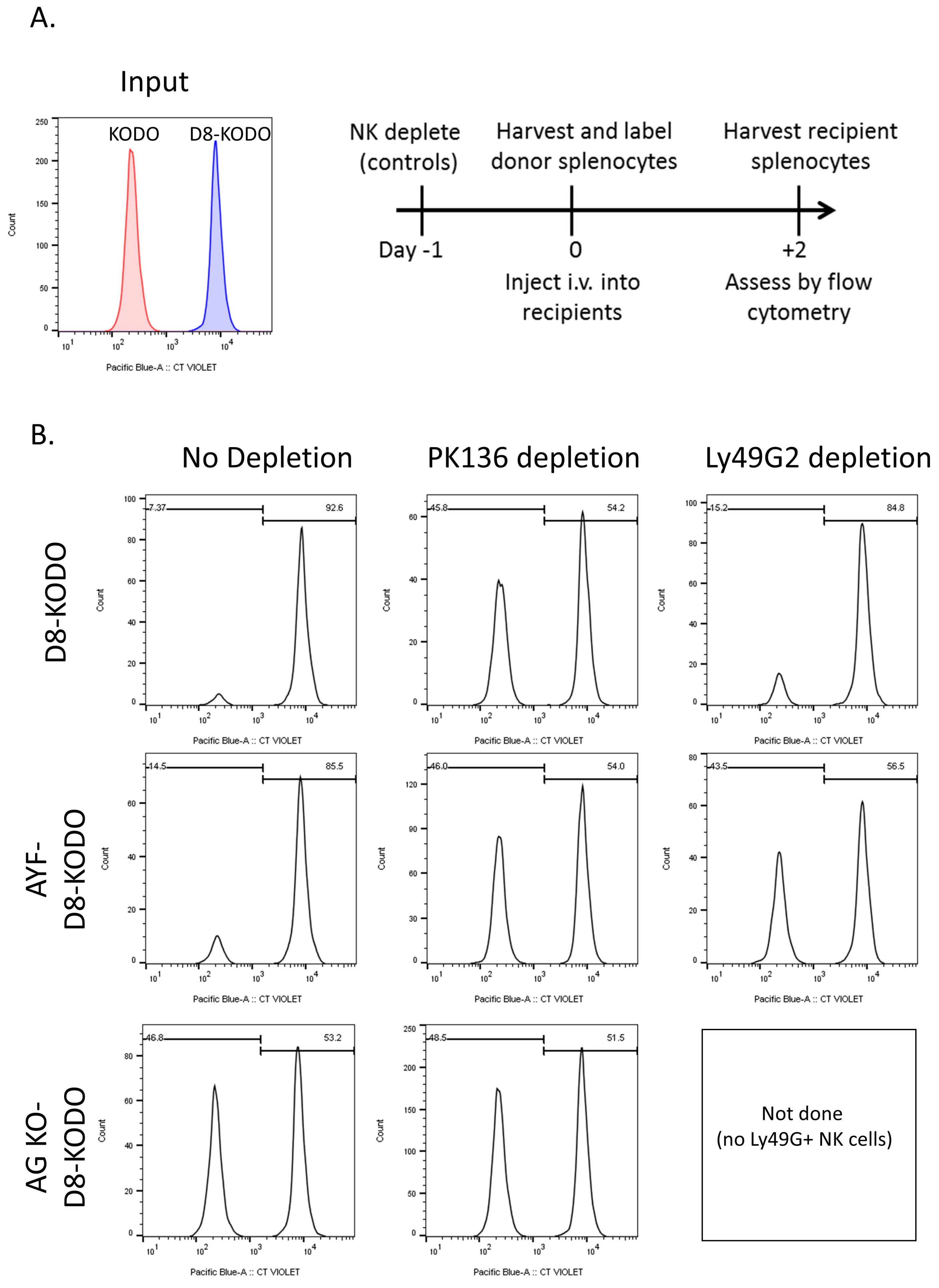


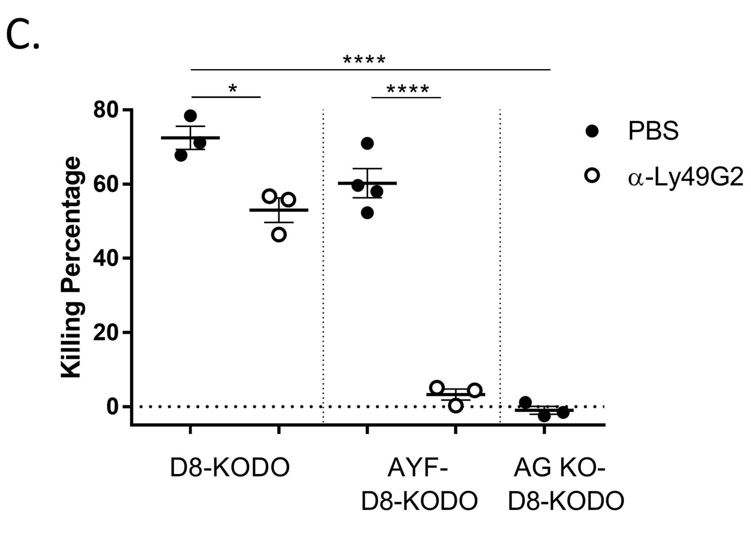


**Fig S4.** ***In vivo*** **cytotoxicity in Ly49AG KO and Ly49AYF mice.**

*In vivo* cytotoxicity of MHC-I deficient (KODO) or sufficient (D8-KODO) splenocytes following differential labelling with Celltrace Violet and flow cytometric analysis recovered from spleens at 2 days post-injection. (A) Flow cytometric analysis of input cells is shown (prior to injection). The experimental design is also depicted with the timing of antibody depletion relative to injection and harvest. (B) Representative histogram plots of the mice indicated in each row and treatments by column. The peaks are identified by the input plot shown in (A). (C) Quantification was performed as previously described (**Fig 4C**). Data are representative of two independent experiments with three recipient mice per group that received the same mix of donor cells in each experiment. Standard error of the mean is shown; statistical analysis performed with a Student's *t*-Test. $*p < .05$, $****p < .0001$.

A.

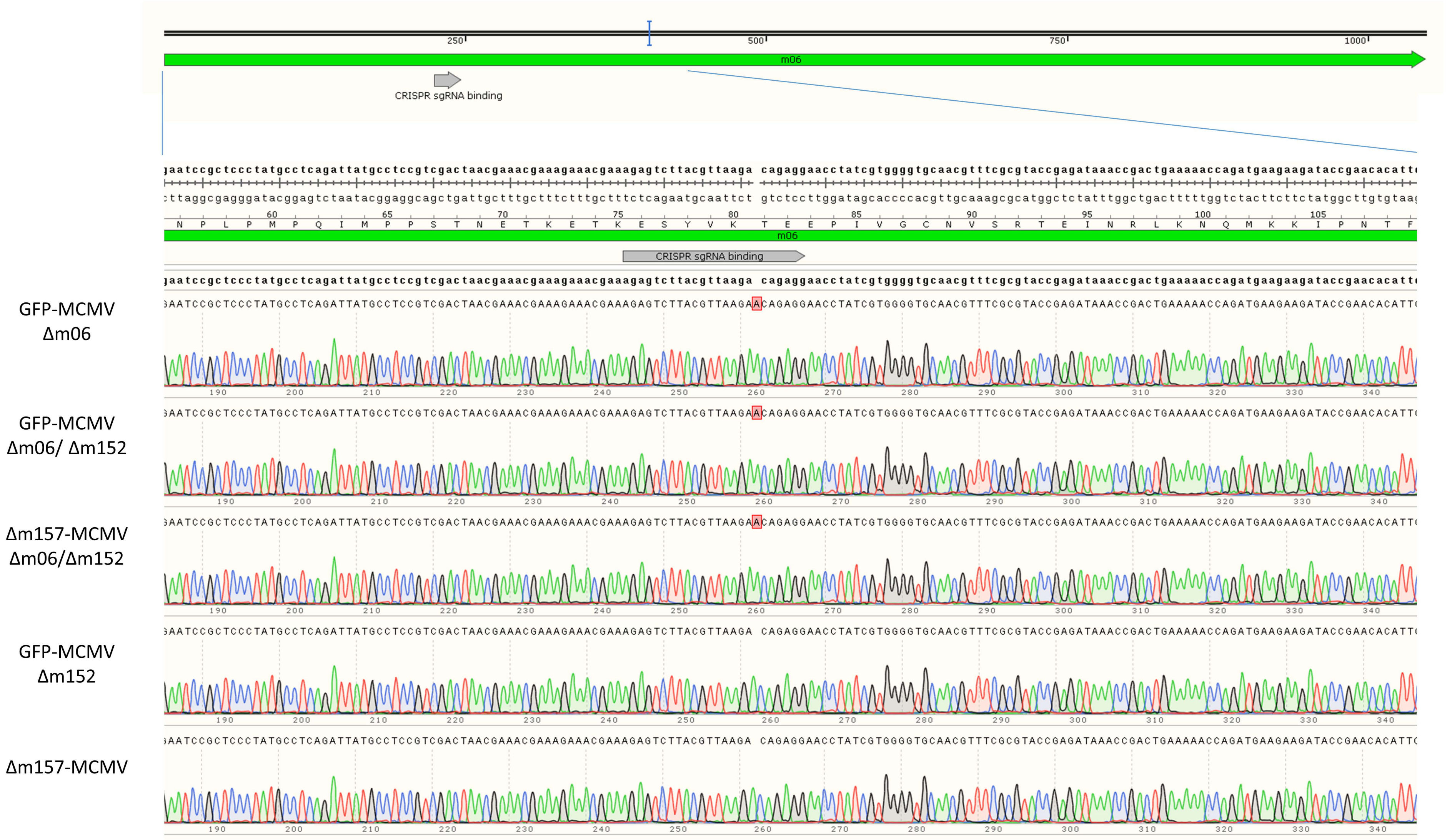


B.

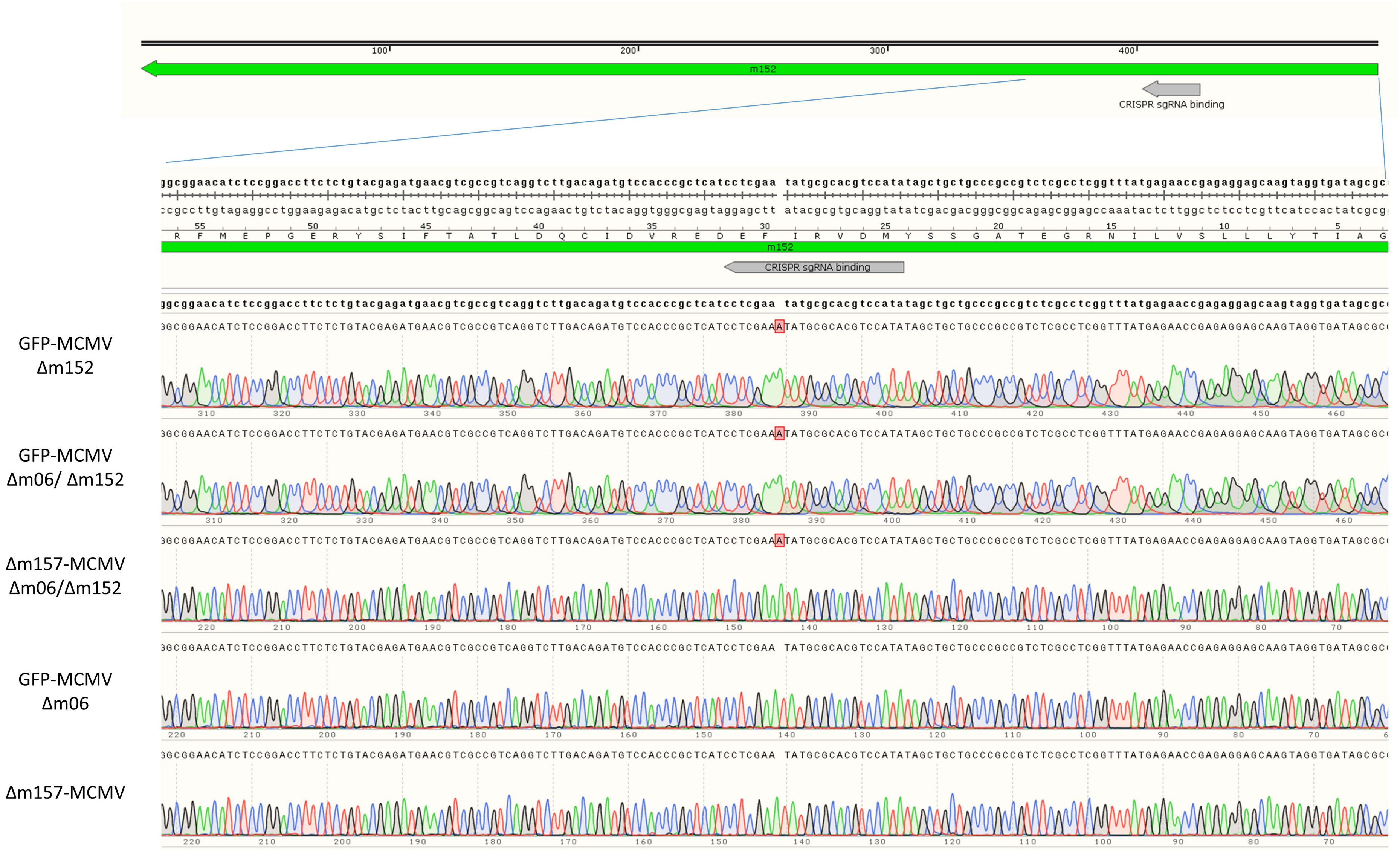


**Fig S5. Sanger sequence analysis of m06 and m152 in CRISPR-modified MCMV strains.**
Viral DNA was isolated from *in vitro* cell cultures infected with CRISPR-Cas9 modified MCMV used in these studies. The specific CRISPR-targeted ORFs, (A) *m06* and (B) *m152*, were Sanger sequenced to confirm and characterize the frameshift in on-target variants. The primers used for PCR and subsequent sequence analysis are shown in **Table S5**. Each panel depicts the reference sequence above with the region of the ORF expanded for analysis. The Sanger alignments below for the viruses indicated are designated as to which strains (*left* of trace) were analyzed. Δm157-MCMV is the wild-type genome at these ORFs, for comparison. Red boxes indicate where inserted nucleotides were identified. Sequence analysis was confirmed in both the forward and reverse direction, however, only one direction is shown for clarity.

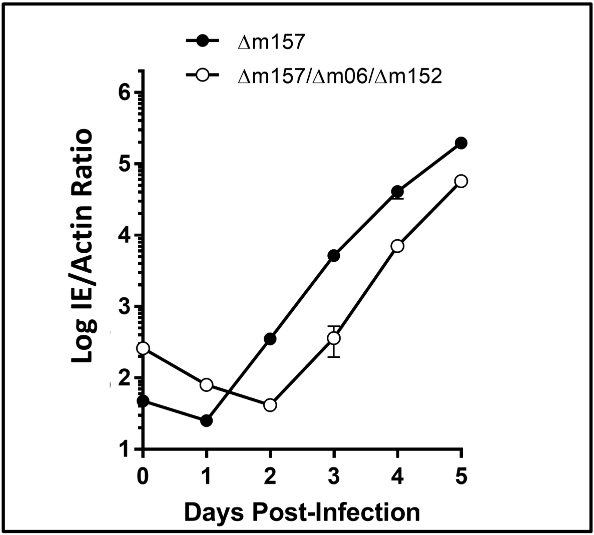


**Fig S6. Multi-step growth curve**

Multi-step in vitro growth kinetics of two strains of MCMV, as indicated. Cells and supernatants were harvested at the indicated days and quantified by real-time PCR. Data is a representative of two independent experiments with each data point performed in triplicate. All timepoints were shown to significantly different in terms of viral load ($p$ < .0001); statistical analysis performed with a Student's *t*-Test.

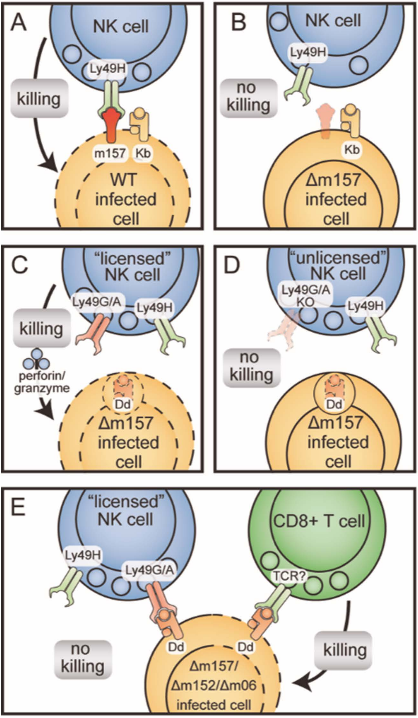


**Fig S7. Model for MHC-restricted and Ly49 receptor-dependent MCMV resistance.**

(A) Activation receptor Ly49H-dependent induces killing of m157-expressing MCMV-infected cells. (B) Deficiency of m157 prevents rejection of MCMV in B6 mice expressing $H2K^b$ ($H2D^b$ is present but not shown for clarity). (C) Granzyme and perforin expression on licensed NK cells are required for control of m157-deficient virus in $H2D^d$-expressing mice. (D) MHC-I downmodulation by MCMV prevents recognition of missing-self rejection in mice lacking Ly49A and Ly49G with unlicensed NK cells. (E) MCMV ORFs, *m06* and *m152*, are essential for the NK-specific protective response and the loss of these molecules results in the emergence of T cell-dependent resistance and concomitant loss of NK cell-dependent control by licensed NK cells.
