## Supplemental Table S1 for "Major histocompatibility complex class I-restricted protection against murine cytomegalovirus requires missing-self recognition by the natural killer cell inhibitory Ly49 receptors"

| Guide | Target | Sequence + PAM | Mouse Generated | Chr | Off-Target sites Worksheet |
|---|---|---|---|---|---|
| G3 | *Ly49g/a/h* | CATTCCCAAGATGAGTGAGCAGG | ΔLy49-1 | 6 | G3 |
| I2 | *Ly49i* | GACCTCCGGCTCATTCATCGTGG | C14B | 6 | I2 |
| C1 | *Ly49c* | TCCCACGATGAGTGAGCCAGAGG | C14B | 6 | C1 |
| M1 | *Ly49m* | CGGGTGAGGCTTGAGGAGACAGG | M1, M4, M7, M8 | 6 | M1 |
| G4 | *Ly49g* | GCAGAAACTAGTGAGGACTGAGG | AG1, AG7 | 6 | G4 |
| A5 | *Ly49a* | GTGAGACCTGAGGAGACTAAAGG | AG1, AG7, A15, A20 | 6 | A5 |
| NKP46-T3 | *Ncr1* | GGTACAGCATAGAGCTCACAAGG | Ly49A-KI | 7 | T3 |
| NKP46-T1 | *Ncr1* | GTGAGCTCTATGCTGTACCCTGG | Ly49A-KI | 7 | T1 |

The names of the sgRNA guides (Guide ID) are provided along with the intended genomic targets, the sgRNA sequence including the "NGG" protospacer adjacent motif (PAM) site underlined, the mice generated with the sgRNA and the chromosomal location of the targeted gene. A complete list containing all relevant on-target and off-target sites in B6 mice is provided as worksheets linked within this file and references the specific guides using the Guide ID. In this file, individual on-target and potential off-target sites are indicated as identified by CCTop (see Methods). Specifically, the input sgRNA sequence along with the number of mismatches and the location of mismatches within the 12nt core (less likely to bind) or outside the core are shown. Chromosome positions of targets along with gene names, if available are indicated. Potential off-target sites within the target chromosome are highlighted and were carefully examined with sequence (**Fig S1**) and flow cytometric methods (**Fig S2**).

Species: Mouse (Mus musculus GRCm38/mm10)
Input: CCTGCTCACTCATCTTGGGAATG
PAM: NGG
Target site length: 20
Core length: 12
Core MM: 2
Total MM: 4

T1 CATTCCCAAGATGAGTGAGCAGG 1000 CRISPRater score 0.597489705

<table>
<tr><th>Chromosome</th><th>start</th><th>end</th><th>strand</th><th>MM</th><th>target_seq</th><th>PAM</th><th>alignment</th><th>distance</th><th>position</th><th>gene name</th><th>gene id</th></tr>
<tr><td>chr6</td><td>130231605</td><td>130231627</td><td>-</td><td>0</td><td>CATTCCCAAGATGAGTGAGC</td><td>AGG</td><td>||||||||[||||||||||||]PAM</td><td>0</td><td>Exonic</td><td>Klra7</td><td>ENSMUSG00000067599</td></tr>
<tr><td>chr6</td><td>130160736</td><td>130160758</td><td>-</td><td>1</td><td>TATTCCCAAGATGAGTGAGC</td><td>AGG</td><td>-|||||||[||||||||||||]PAM</td><td>0</td><td>Exonic</td><td>Klra14-ps</td><td>ENSMUSG00000072721</td></tr>
<tr><td>chr6</td><td>130128148</td><td>130128170</td><td>-</td><td>1</td><td>TATTCCCAAGATGAGTGAGC</td><td>AGG</td><td>-|||||||[||||||||||||]PAM</td><td>0</td><td>Exonic</td><td>Klra8</td><td>ENSMUSG00000089727</td></tr>
<tr><td>chr6</td><td>130380650</td><td>130380672</td><td>-</td><td>1</td><td>CAATCCCAAGATGAGTGAGC</td><td>AGG</td><td>||-|||||[||||||||||||]PAM</td><td>0</td><td>Exonic</td><td>Klra1</td><td>ENSMUSG00000079853</td></tr>
<tr><td>chr6</td><td>130306415</td><td>130306437</td><td>-</td><td>3</td><td>TCCTCCCAAGATGAGTGAGC</td><td>AGG</td><td>---|||||[||||||||||||]PAM</td><td>0</td><td>Exonic</td><td>Klra13-ps</td><td>ENSMUSG00000030178</td></tr>
<tr><td>chr6</td><td>131245314</td><td>131245336</td><td>-</td><td>3</td><td>TACTTCCAAGATGAGTGAGC</td><td>AGG</td><td>-|-|-|||[||||||||||||]PAM</td><td>0</td><td>Exonic</td><td>Klra2</td><td>ENSMUSG00000030187</td></tr>
<tr><td>chr13</td><td>55077477</td><td>55077499</td><td>-</td><td>4</td><td>GGCTCTCAAGATGAGTGAGC</td><td>AGG</td><td>---||-||[||||||||||||]PAM</td><td>0</td><td>Exonic</td><td>Uimc1</td><td>ENSMUSG00000025878</td></tr>
<tr><td>chr6</td><td>130281910</td><td>130281932</td><td>-</td><td>3</td><td>TACTCCCACGATGAGTGAGC</td><td>TGG</td><td>-|-|||||[-|||||||||||]PAM</td><td>0</td><td>Exonic</td><td>Klra10</td><td>ENSMUSG00000072718</td></tr>
<tr><td>chr15</td><td>73072809</td><td>73072831</td><td>-</td><td>2</td><td>GATTCCCAAGAAGAGTGAGC</td><td>AGG</td><td>-|||||||[|||-||||||||]PAM</td><td>11605</td><td>Intergenic</td><td>Trappc9</td><td>ENSMUSG00000047921</td></tr>
<tr><td>chr6</td><td>129874895</td><td>129874917</td><td>-</td><td>4</td><td>TACACCCATGATGAGTGAGC</td><td>AGG</td><td>-|--||||[-|||||||||||]PAM</td><td>0</td><td>Exonic</td><td>Klra17</td><td>ENSMUSG00000014543</td></tr>
<tr><td>chr7</td><td>19522381</td><td>19522403</td><td>-</td><td>4</td><td>GAGGCCCATGATGAGTGAGC</td><td>GGG</td><td>-|--||||[-|||||||||||]PAM</td><td>316</td><td>Intronic</td><td>Nkpd1</td><td>ENSMUSG00000060621</td></tr>
<tr><td>chr10</td><td>98815111</td><td>98815133</td><td>+</td><td>4</td><td>CAGGCACGAGATGAGTGAGC</td><td>TGG</td><td>||--|-|-[||||||||||||]PAM</td><td>NA</td><td>Intergenic</td><td>NA</td><td>NA</td></tr>
<tr><td>chr6</td><td>130407926</td><td>130407948</td><td>-</td><td>3</td><td>TACTCCCAAGACGAGTGAGC</td><td>AGG</td><td>-|-|||||[|||-||||||||]PAM</td><td>0</td><td>Exonic</td><td>Gm6584</td><td>ENSMUSG00000084204</td></tr>
<tr><td>chr4</td><td>102882973</td><td>102882995</td><td>+</td><td>4</td><td>TAATTCCAAGTTGAGTGAGC</td><td>AGG</td><td>-|-|-|||[||-|||||||||]PAM</td><td>0</td><td>Exonic</td><td>Sgip1</td><td>ENSMUSG00000028524</td></tr>
<tr><td>chr6</td><td>130094664</td><td>130094686</td><td>-</td><td>2</td><td>TATTCCCAAGATGGGTGAGC</td><td>AGG</td><td>-|||||||[|||||-||||||]PAM</td><td>0</td><td>Exonic</td><td>Klra11-ps</td><td>ENSMUSG00000072717</td></tr>
<tr><td>chr6</td><td>130025207</td><td>130025229</td><td>-</td><td>4</td><td>TACTCCCACAATGAGTGAGC</td><td>CGG</td><td>-|-|||||[--||||||||||]PAM</td><td>0</td><td>Exonic</td><td>Klra6</td><td>ENSMUSG00000061769</td></tr>
<tr><td>chr11</td><td>101325482</td><td>101325504</td><td>+</td><td>4</td><td>CAAGCCCAACGTGAGTGAGC</td><td>TGG</td><td>||--||||[|--|||||||||]PAM</td><td>0</td><td>Exonic</td><td>Aoc2</td><td>ENSMUSG00000078651</td></tr>
<tr><td>chr11</td><td>101331341</td><td>101331363</td><td>+</td><td>4</td><td>CAAGCCCAACGTGAGTGAGC</td><td>TGG</td><td>||--||||[|--|||||||||]PAM</td><td>0</td><td>Exonic</td><td>Aoc3</td><td>ENSMUSG00000019326</td></tr>
<tr><td>chr14</td><td>101503678</td><td>101503700</td><td>-</td><td>4</td><td>CATGGCCAATGTGAGTGAGC</td><td>CGG</td><td>|||--|||[|--|||||||||]PAM</td><td>1007</td><td>Intronic</td><td>Tbc1d4</td><td>ENSMUSG00000033083</td></tr>
<tr><td>chr7</td><td>30110644</td><td>30110666</td><td>-</td><td>4</td><td>CTATCCCAACAGGAGTGAGC</td><td>TGG</td><td>|--|||||[|-|-||||||||]PAM</td><td>3022</td><td>Intergenic</td><td>Zfp260</td><td>ENSMUSG00000049421</td></tr>
<tr><td>chr7</td><td>96636324</td><td>96636346</td><td>-</td><td>4</td><td>AAGTCCCAAGCTCAGTGAGC</td><td>AGG</td><td>-|-|||||[||-|-|||||||]PAM</td><td>7255</td><td>Intronic</td><td>Gm15414</td><td>ENSMUSG00000085792</td></tr>
<tr><td>chr9</td><td>120082119</td><td>120082141</td><td>+</td><td>4</td><td>CATAACCAGGATGGGTGAGC</td><td>AGG</td><td>|||--|||[-||||-||||||]PAM</td><td>9992</td><td>Intergenic</td><td>Ccr8</td><td>ENSMUSG00000042262</td></tr>
<tr><td>chr2</td><td>90802873</td><td>90802895</td><td>+</td><td>4</td><td>AGTTCCCAAGCTGTGTGAGC</td><td>AGG</td><td>--||||||[||-||-||||||]PAM</td><td>147</td><td>Intronic</td><td>Agbl2</td><td>ENSMUSG00000040812</td></tr>
<tr><td>chr10</td><td>108909798</td><td>108909820</td><td>-</td><td>4</td><td>AGTTCCCAAGCTGCGTGAGC</td><td>AGG</td><td>--||||||[||-||-||||||]PAM</td><td>7483</td><td>Intronic</td><td>Syt1</td><td>ENSMUSG00000035864</td></tr>
<tr><td>chr6</td><td>130191289</td><td>130191311</td><td>-</td><td>4</td><td>TACTCCCACGATGAATGAGC</td><td>CGG</td><td>-|-|||||[-|||||-|||||]PAM</td><td>0</td><td>Exonic</td><td>Klra9</td><td>ENSMUSG00000033024</td></tr>
<tr><td>chr18</td><td>50314596</td><td>50314618</td><td>+</td><td>4</td><td>CTTTTCCTAGATGACTGAGC</td><td>TGG</td><td>|-||-||-[||||||-|||||]PAM</td><td>31577</td><td>Intergenic</td><td>Fam170a</td><td>ENSMUSG00000035420</td></tr>
<tr><td>chr12</td><td>112277749</td><td>112277771</td><td>+</td><td>4</td><td>CCTTCCCCAGGTCAGTGAGC</td><td>AGG</td><td>|-|||||-[||-|-|||||||]PAM</td><td>69518</td><td>Intergenic</td><td>Gm38123</td><td>ENSMUSG00000103080</td></tr>
<tr><td>chr9</td><td>59176101</td><td>59176123</td><td>-</td><td>4</td><td>CTTCCCTAAGATGAGAGAGC</td><td>AGG</td><td>|-|-||-|[|||||||-||||]PAM</td><td>29891</td><td>Intergenic</td><td>Gm7589</td><td>ENSMUSG00000066592</td></tr>
<tr><td>chr15</td><td>41502054</td><td>41502076</td><td>-</td><td>4</td><td>CATCCCTAAGAGGTGTGAGC</td><td>AGG</td><td>|||-||-|[|||-|-||||||]PAM</td><td>33494</td><td>Intronic</td><td>Oxr1</td><td>ENSMUSG00000022307</td></tr>
<tr><td>chr3</td><td>68153939</td><td>68153961</td><td>+</td><td>4</td><td>CCTTACCAAGATAGGTGAGC</td><td>AGG</td><td>|-||-|||[||||--||||||]PAM</td><td>88932</td><td>Intronic</td><td>Schip1</td><td>ENSMUSG00000027777</td></tr>
<tr><td>chrX</td><td>160605795</td><td>160605817</td><td>+</td><td>4</td><td>TGTTCCCAAGCTGAGGGAGC</td><td>TGG</td><td>--||||||[||-||||-||||]PAM</td><td>6917</td><td>Intergenic</td><td>Phka2</td><td>ENSMUSG00000031295</td></tr>
<tr><td>chr1</td><td>92236338</td><td>92236360</td><td>+</td><td>4</td><td>CAATCACAAGAGGAATGAGC</td><td>AGG</td><td>||-||-||[|||-||-|||||]PAM</td><td>55997</td><td>Intergenic</td><td>Hdac4</td><td>ENSMUSG00000026313</td></tr>
<tr><td>chr14</td><td>68278685</td><td>68278707</td><td>+</td><td>4</td><td>CTTGCCCAAGATCATTGAGC</td><td>AGG</td><td>|-|-||||[||||-|-|||||]PAM</td><td>NA</td><td>Intergenic</td><td>NA</td><td>NA</td></tr>
<tr><td>chr10</td><td>18910845</td><td>18910867</td><td>-</td><td>4</td><td>CTTTCCCCAGAGGACTGAGC</td><td>AGG</td><td>|-|||||-[|||-||-|||||]PAM</td><td>10521</td><td>Intergenic</td><td>Gm38175</td><td>ENSMUSG00000103820</td></tr>
<tr><td>chr14</td><td>79076122</td><td>79076144</td><td>+</td><td>4</td><td>GATCTCCAAGATGAGTGTGC</td><td>TGG</td><td>-||--|||[|||||||||-||]PAM</td><td>4969</td><td>Intronic</td><td>Vwa8</td><td>ENSMUSG00000058997</td></tr>
<tr><td>chr7</td><td>145195430</td><td>145195452</td><td>+</td><td>4</td><td>CAATACCAAGATGGATGAGC</td><td>TGG</td><td>||-|-|||[|||||--|||||]PAM</td><td>58471</td><td>Intergenic</td><td>Tpcn2</td><td>ENSMUSG00000048677</td></tr>
<tr><td>chr2</td><td>167581093</td><td>167581115</td><td>-</td><td>4</td><td>CATTGCACAGATGAGTAAGC</td><td>TGG</td><td>||||-|--[||||||||-|||]PAM</td><td>10280</td><td>Intronic</td><td>Gm11475</td><td>ENSMUSG00000081511</td></tr>
<tr><td>chr15</td><td>24386242</td><td>24386264</td><td>-</td><td>4</td><td>CATTACACAGATGAGTTAGC</td><td>GGG</td><td>||||-|--[||||||||-|||]PAM</td><td>NA</td><td>Intergenic</td><td>NA</td><td>NA</td></tr>
<tr><td>chr1</td><td>41038254</td><td>41038276</td><td>-</td><td>4</td><td>CACACCCAAGGTGAGTCAGC</td><td>TGG</td><td>||--||||[||-|||||-|||]PAM</td><td>NA</td><td>Intergenic</td><td>NA</td><td>NA</td></tr>
<tr><td>chr9</td><td>50180476</td><td>50180498</td><td>-</td><td>4</td><td>CATTGCTAAGATGGCTGAGC</td><td>TGG</td><td>||||-|-|[|||||--|||||]PAM</td><td>5545</td><td>Intergenic</td><td>Gm8907</td><td>ENSMUSG00000098012</td></tr>
<tr><td>chr5</td><td>3877184</td><td>3877206</td><td>+</td><td>4</td><td>CACACCCAAGAGGAGTCAGC</td><td>AGG</td><td>||--||||[|||-||||-|||]PAM</td><td>1030</td><td>Intergenic</td><td>Rps4x-ps</td><td>ENSMUSG00000104699</td></tr>
<tr><td>chr19</td><td>29217804</td><td>29217826</td><td>-</td><td>4</td><td>CATTAATAAGATGAGTGTGC</td><td>AGG</td><td>||||---|[|||||||||-||]PAM</td><td>34002</td><td>Intergenic</td><td>Jak2</td><td>ENSMUSG00000024789</td></tr>
<tr><td>chr1</td><td>173455229</td><td>173455251</td><td>-</td><td>3</td><td>CATACCCAAGATGACAGAGC</td><td>AGG</td><td>|||-||||[||||||--||||]PAM</td><td>83</td><td>Intronic</td><td>Aim2</td><td>ENSMUSG00000037860</td></tr>
<tr><td>chr14</td><td>10779509</td><td>10779531</td><td>+</td><td>4</td><td>CATCCACAAGAGGAGTAAGC</td><td>TGG</td><td>|||-|-||[|||-||||-|||]PAM</td><td>508</td><td>Intronic</td><td>Fhitos</td><td>ENSMUSG00000089833</td></tr>
<tr><td>chr11</td><td>44019220</td><td>44019242</td><td>+</td><td>4</td><td>AAGTCCCAAGATGACAGAGC</td><td>AGG</td><td>-|-|||||[||||||--||||]PAM</td><td>7275</td><td>Intronic</td><td>Gm12153</td><td>ENSMUSG00000087172</td></tr>
<tr><td>chr3</td><td>149798955</td><td>149798977</td><td>-</td><td>4</td><td>TAGTCCCAAGATGATAGAGC</td><td>TGG</td><td>-|-|||||[||||||--||||]PAM</td><td>NA</td><td>Intergenic</td><td>NA</td><td>NA</td></tr>
<tr><td>chr18</td><td>9533263</td><td>9533285</td><td>-</td><td>4</td><td>CAGTCCCTGGATGAGTGGGC</td><td>AGG</td><td>||-||||-[-||||||||-||]PAM</td><td>81029</td><td>Intergenic</td><td>Gm22765</td><td>ENSMUSG00000088996</td></tr>
<tr><td>chr17</td><td>33074253</td><td>33074275</td><td>-</td><td>4</td><td>CATTGACAAGATCAGTCAGC</td><td>AGG</td><td>||||--||[||||-|||-|||]PAM</td><td>6195</td><td>Intergenic</td><td>4921501E0</td><td>ENSMUSG00000023350</td></tr>
<tr><td>chr9</td><td>74043734</td><td>74043756</td><td>+</td><td>4</td><td>GATTGCCAAGATGGGTCAGC</td><td>GGG</td><td>-|||-|||[|||||-||-|||]PAM</td><td>15583</td><td>Intergenic</td><td>n-R5s86</td><td>ENSMUSG00000095631</td></tr>
<tr><td>chr13</td><td>51685161</td><td>51685183</td><td>+</td><td>4</td><td>CATTAGCCAGATGAGTGACC</td><td>AGG</td><td>||||--|-[||||||||||-|]PAM</td><td>346</td><td>Intergenic</td><td>Gm15440</td><td>ENSMUSG00000051107</td></tr>
<tr><td>chr17</td><td>56022053</td><td>56022075</td><td>-</td><td>4</td><td>CAGACCCAACATGAGTGACC</td><td>AGG</td><td>||--||||[|-||||||||-|]PAM</td><td>1862</td><td>Intronic</td><td>Sh3gl1</td><td>ENSMUSG00000003200</td></tr>
<tr><td>chr15</td><td>37848268</td><td>37848290</td><td>+</td><td>4</td><td>ACTTCTCAAGATGAGTGAGT</td><td>AGG</td><td>--|||-||[|||||||||||-]PAM</td><td>19796</td><td>Intergenic</td><td>Rps19-ps5</td><td>ENSMUSG00000097679</td></tr>
<tr><td>chr2</td><td>80408988</td><td>80409010</td><td>-</td><td>4</td><td>CTTTTGCAAGATGAGTGAGG</td><td>TGG</td><td>|-||--||[|||||||||||-]PAM</td><td>2960</td><td>Intergenic</td><td>Frzb</td><td>ENSMUSG00000027004</td></tr>
<tr><td>chr3</td><td>66883057</td><td>66883079</td><td>+</td><td>4</td><td>CACACCAAAGATGAGTGAGT</td><td>GGG</td><td>||--||-|[|||||||||||-]PAM</td><td>0</td><td>Exonic</td><td>Gm6555</td><td>ENSMUSG00000103700</td></tr>
<tr><td>chr8</td><td>82495716</td><td>82495738</td><td>+</td><td>4</td><td>CATTGGCAAGATGAGGTAGC</td><td>AGG</td><td>||||--||[|||||||--|||]PAM</td><td>82029</td><td>Intergenic</td><td>Gm8167</td><td>ENSMUSG00000097984</td></tr>
<tr><td>chr11</td><td>95507363</td><td>95507385</td><td>-</td><td>4</td><td>CTTTCCCTAGATGAGCAAGC</td><td>GGG</td><td>|-|||||-[|||||||--|||]PAM</td><td>2460</td><td>Intergenic</td><td>Nxph3</td><td>ENSMUSG00000046719</td></tr>
<tr><td>chr5</td><td>22094531</td><td>22094553</td><td>-</td><td>4</td><td>CATTTCGAAGATGAGGAAGC</td><td>AGG</td><td>||||-|-|[|||||||--|||]PAM</td><td>8671</td><td>Intronic</td><td>Reln</td><td>ENSMUSG00000042453</td></tr>
<tr><td>chr7</td><td>63664289</td><td>63664311</td><td>-</td><td>4</td><td>CATTCTTAAGATGAGAAAGC</td><td>AGG</td><td>|||||--|[|||||||--|||]PAM</td><td>13324</td><td>Intronic</td><td>Otud7a</td><td>ENSMUSG00000033510</td></tr>
<tr><td>chr13</td><td>97924305</td><td>97924327</td><td>-</td><td>4</td><td>GAGTCCCAAGATGAGGGTGC</td><td>TGG</td><td>-|-|||||[|||||||-|-||]PAM</td><td>5084</td><td>Intronic</td><td>Arhgef28</td><td>ENSMUSG00000021662</td></tr>
<tr><td>chr6</td><td>130065290</td><td>130065312</td><td>-</td><td>4</td><td>TACTCCCAAGATGACTGAAC</td><td>AGG</td><td>-|-|||||[||||||-|||-|]PAM</td><td>0</td><td>Exonic</td><td>Klra4</td><td>ENSMUSG00000079852</td></tr>
<tr><td>chr7</td><td>70787617</td><td>70787639</td><td>-</td><td>4</td><td>TATTCCAAAGGTGAGTGAGA</td><td>GGG</td><td>-|||||-|[||-||||||||-]PAM</td><td>35117</td><td>Intergenic</td><td>Gm24880</td><td>ENSMUSG00000089401</td></tr>
<tr><td>chr18</td><td>24473609</td><td>24473631</td><td>-</td><td>4</td><td>CATACACAAGGTGAGTGAGG</td><td>AGG</td><td>|||-|-||[||-||||||||-]PAM</td><td>460</td><td>Intronic</td><td>2700062C0</td><td>ENSMUSG00000024273</td></tr>
<tr><td>chr14</td><td>32443836</td><td>32443858</td><td>-</td><td>4</td><td>CAGACCCAAGATGACTGAAC</td><td>GGG</td><td>||--||||[||||||-|||-|]PAM</td><td>2929</td><td>Intronic</td><td>Chat</td><td>ENSMUSG00000021919</td></tr>
<tr><td>chr3</td><td>43048978</td><td>43049000</td><td>-</td><td>4</td><td>CAGTTCCAAGAAGAGTGAGT</td><td>GGG</td><td>||-|-|||[|||-|||||||-]PAM</td><td>NA</td><td>Intergenic</td><td>NA</td><td>NA</td></tr>
<tr><td>chr10</td><td>80082999</td><td>80083021</td><td>+</td><td>4</td><td>CATGGCCAAGAAGAGTGAGG</td><td>AGG</td><td>|||--|||[|||-|||||||-]PAM</td><td>273</td><td>Intronic</td><td>Sbno2</td><td>ENSMUSG00000035673</td></tr>
<tr><td>chr3</td><td>82553453</td><td>82553475</td><td>-</td><td>4</td><td>CATGCACAAGATGACTGACC</td><td>TGG</td><td>|||-|-||[||||||-|||-|]PAM</td><td>5369</td><td>Intergenic</td><td>Npy2r</td><td>ENSMUSG00000028004</td></tr>
<tr><td>chr11</td><td>19633014</td><td>19633036</td><td>+</td><td>4</td><td>AATTTCCAAGATGAGTCTGC</td><td>TGG</td><td>-|||-|||[||||||||--||]PAM</td><td>4953</td><td>Intergenic</td><td>Gm12027</td><td>ENSMUSG00000086911</td></tr>
<tr><td>chr11</td><td>86566488</td><td>86566510</td><td>+</td><td>4</td><td>AATTCCCCAGATGACTGAAC</td><td>AGG</td><td>-||||||-[||||||-|||-|]PAM</td><td>527</td><td>Intronic</td><td>Tubd1</td><td>ENSMUSG00000020513</td></tr>
<tr><td>chrX</td><td>118896177</td><td>118896199</td><td>-</td><td>4</td><td>CAGTGCCAAGATGTGTGAGT</td><td>AGG</td><td>||-|-|||[|||||-|||||-]PAM</td><td>NA</td><td>Intergenic</td><td>NA</td><td>NA</td></tr>
<tr><td>chr10</td><td>105681894</td><td>105681916</td><td>+</td><td>4</td><td>CATCCCCCAGATGAGAGATC</td><td>TGG</td><td>|||-|||-[|||||||-||-|]PAM</td><td>81273</td><td>Intergenic</td><td>Mettl25</td><td>ENSMUSG00000036009</td></tr>
<tr><td>chr11</td><td>108553978</td><td>108554000</td><td>-</td><td>4</td><td>CATTTGCAAGATGAGTCAAC</td><td>TGG</td><td>||||--||[||||||||-|-|]PAM</td><td>16288</td><td>Intronic</td><td>Cep112</td><td>ENSMUSG00000020728</td></tr>
<tr><td>chr4</td><td>152206815</td><td>152206837</td><td>+</td><td>4</td><td>CATTCTGAAGATGATTGAGG</td><td>AGG</td><td>|||||--|[||||||-||||-]PAM</td><td>0</td><td>Exonic</td><td>Acot7</td><td>ENSMUSG00000028937</td></tr>
<tr><td>chr1</td><td>151369669</td><td>151369691</td><td>-</td><td>3</td><td>CATTACCAAGATGAGTGCTC</td><td>AGG</td><td>||||-|||[|||||||||--|]PAM</td><td>693</td><td>Intronic</td><td>Swt1</td><td>ENSMUSG00000052748</td></tr>
<tr><td>chr19</td><td>35375161</td><td>35375183</td><td>-</td><td>4</td><td>CATGGCCAAGATGAGAGAGG</td><td>GGG</td><td>|||--|||[|||||||-|||-]PAM</td><td>98639</td><td>Intergenic</td><td>Gm23075</td><td>ENSMUSG00000093373</td></tr>
<tr><td>chr12</td><td>106173901</td><td>106173923</td><td>+</td><td>4</td><td>AATTCTCAAGATGAGTGTCC</td><td>AGG</td><td>-||||-||[|||||||||--|]PAM</td><td>47152</td><td>Intergenic</td><td>Gm24986</td><td>ENSMUSG00000077640</td></tr>
<tr><td>chr8</td><td>106523883</td><td>106523905</td><td>+</td><td>4</td><td>CCTTCACAAGATGAGTGCTC</td><td>AGG</td><td>|-|||-||[|||||||||--|]PAM</td><td>11503</td><td>Intronic</td><td>Cdh3</td><td>ENSMUSG00000061048</td></tr>
<tr><td>chr4</td><td>140903499</td><td>140903521</td><td>+</td><td>4</td><td>CTTTCCCCAGATGAGTGGGG</td><td>TGG</td><td>|-|||||-[|||||||||-|-]PAM</td><td>2823</td><td>Intergenic</td><td>Padi2</td><td>ENSMUSG00000028927</td></tr>
</table>

Species: Mouse (Mus musculus GRCm38/mm10)
Input: GACCTCCGGCTCATTCATCGTGG
PAM: NGG
Target site length: 20
Core length: 12
Core MM: 2
Total MM: 4

<table>
<tr><td>T1</td><td>GACCTCCGGCTCATTCATCGTGG</td><td>1000</td><td>CRISPRater score</td><td>0.819410535</td><td></td><td></td><td></td><td></td><td></td><td></td><td></td></tr>
<tr><td>Chromosome</td><td>start</td><td>end</td><td>strand</td><td>MM</td><td>target_seq</td><td>PAM</td><td>alignment</td><td>distance</td><td>position</td><td>gene name</td><td>gene id</td></tr>
<tr><td>chr6</td><td>130191284</td><td>130191306</td><td>+</td><td>0</td><td>GACCTCCGGCTCATTCATCG</td><td>TGG</td><td>||||||||[||||||||||||]PAM</td><td>0</td><td>Exonic</td><td>Klra9</td><td>ENSMUSG00000033024</td></tr>
<tr><td>chr6</td><td>74205167</td><td>74205189</td><td>-</td><td>4</td><td>GTCTTCCACCTCATTCATCG</td><td>AGG</td><td>|-|-|||-[-|||||||||||]PAM</td><td>NA</td><td>Intergenic</td><td>NA</td><td>NA</td></tr>
<tr><td>chr6</td><td>130335782</td><td>130335804</td><td>+</td><td>2</td><td>GACCTCTGGCTCACTCATCG</td><td>TGG</td><td>||||||-|[|||||-||||||]PAM</td><td>0</td><td>Exonic</td><td>Klra3</td><td>ENSMUSG00000067591</td></tr>
<tr><td>chr6</td><td>130281905</td><td>130281927</td><td>+</td><td>2</td><td>GACCTCCAGCTCACTCATCG</td><td>TGG</td><td>|||||||-[|||||-||||||]PAM</td><td>0</td><td>Exonic</td><td>Klra10</td><td>ENSMUSG00000072718</td></tr>
<tr><td>chr19</td><td>6277250</td><td>6277272</td><td>+</td><td>4</td><td>CACCACCGACTCCTTCATCG</td><td>CGG</td><td>-|||-|||[-|||-|||||||]PAM</td><td>0</td><td>Exonic</td><td>Ehd1</td><td>ENSMUSG00000024772</td></tr>
<tr><td>chr6</td><td>129911444</td><td>129911466</td><td>+</td><td>3</td><td>GACCTCTGGTTCACTCATCG</td><td>TGG</td><td>||||||-|[|-|||-||||||]PAM</td><td>0</td><td>Exonic</td><td>Klra5</td><td>ENSMUSG00000030173</td></tr>
<tr><td>chr6</td><td>118394078</td><td>118394100</td><td>+</td><td>4</td><td>GACCTGAGTCTCATGCATCG</td><td>AGG</td><td>|||||--|[-|||||-|||||]PAM</td><td>1087</td><td>Intronic</td><td>Bms1</td><td>ENSMUSG00000030138</td></tr>
<tr><td>chr18</td><td>4657158</td><td>4657180</td><td>+</td><td>4</td><td>GAGCCCCGGCACATTCCTCG</td><td>TGG</td><td>||-|-|||[||-|||||-|||]PAM</td><td>7753</td><td>Intronic</td><td>9430020K0</td><td>ENSMUSG00000033960</td></tr>
<tr><td>chr10</td><td>11149812</td><td>11149834</td><td>+</td><td>4</td><td>TCCCTCCGGCTCCTTCTTCG</td><td>AGG</td><td>--||||||[||||-|||-|||]PAM</td><td>226</td><td>Intronic</td><td>Shprh</td><td>ENSMUSG00000090112</td></tr>
<tr><td>chr15</td><td>11995916</td><td>11995938</td><td>-</td><td>4</td><td>GACCCCGGGCCCATTCCTCG</td><td>GGG</td><td>||||-|-|[||-|||||-|||]PAM</td><td>13</td><td>Intronic</td><td>Sub1</td><td>ENSMUSG00000022205</td></tr>
<tr><td>chr1</td><td>18277941</td><td>18277963</td><td>+</td><td>4</td><td>TATCTCTGGCTCATTCATTG</td><td>GGG</td><td>-|-|||-|[||||||||||-|]PAM</td><td>12803</td><td>Intergenic</td><td>Defb41</td><td>ENSMUSG00000067773</td></tr>
<tr><td>chr12</td><td>117519158</td><td>117519180</td><td>+</td><td>4</td><td>CACCTTCTGCTCATTCATAG</td><td>GGG</td><td>-||||-|-[||||||||||-|]PAM</td><td>1022</td><td>Intronic</td><td>Rapgef5</td><td>ENSMUSG00000041992</td></tr>
<tr><td>chr6</td><td>130025202</td><td>130025224</td><td>+</td><td>2</td><td>GACCTCCGGCTCACTCATTG</td><td>TGG</td><td>||||||||[|||||-||||-|]PAM</td><td>0</td><td>Exonic</td><td>Klra6</td><td>ENSMUSG00000061769</td></tr>
<tr><td>chr9</td><td>123294520</td><td>123294542</td><td>+</td><td>4</td><td>AGCCTCCTGCTCATTCATCT</td><td>GGG</td><td>--|||||-[|||||||||||-]PAM</td><td>12966</td><td>Intergenic</td><td>Scp2-ps2</td><td>ENSMUSG00000058492</td></tr>
<tr><td>chr19</td><td>27732746</td><td>27732768</td><td>+</td><td>4</td><td>GATCTCTGGCTCCTTCATTG</td><td>GGG</td><td>||-|||-|[||||-|||||-|]PAM</td><td>28739</td><td>Intergenic</td><td>Gm24782</td><td>ENSMUSG00000088266</td></tr>
<tr><td>chr14</td><td>32426168</td><td>32426190</td><td>-</td><td>4</td><td>GCCCTCCAGCTCCTTCATGG</td><td>TGG</td><td>|-|||||-[||||-|||||-|]PAM</td><td>0</td><td>Exonic</td><td>Chat</td><td>ENSMUSG00000021919</td></tr>
<tr><td>chr6</td><td>130160731</td><td>130160753</td><td>+</td><td>3</td><td>GACCTCCTGCTCACTCATCT</td><td>TGG</td><td>|||||||-[|||||-|||||-]PAM</td><td>0</td><td>Exonic</td><td>Klra14-ps</td><td>ENSMUSG00000072721</td></tr>
<tr><td>chr6</td><td>130128143</td><td>130128165</td><td>+</td><td>3</td><td>GACCTCCTGCTCACTCATCT</td><td>TGG</td><td>|||||||-[|||||-|||||-]PAM</td><td>0</td><td>Exonic</td><td>Klra8</td><td>ENSMUSG00000089727</td></tr>
<tr><td>chr6</td><td>130231600</td><td>130231622</td><td>+</td><td>3</td><td>GACCTCCTGCTCACTCATCT</td><td>TGG</td><td>|||||||-[|||||-|||||-]PAM</td><td>0</td><td>Exonic</td><td>Klra7</td><td>ENSMUSG00000067599</td></tr>
<tr><td>chr6</td><td>130306410</td><td>130306432</td><td>+</td><td>3</td><td>GACCTCCTGCTCACTCATCT</td><td>TGG</td><td>|||||||-[|||||-|||||-]PAM</td><td>0</td><td>Exonic</td><td>Klra13-ps</td><td>ENSMUSG00000030178</td></tr>
<tr><td>chr6</td><td>130380645</td><td>130380667</td><td>+</td><td>3</td><td>GACCTCCTGCTCACTCATCT</td><td>TGG</td><td>|||||||-[|||||-|||||-]PAM</td><td>0</td><td>Exonic</td><td>Klra1</td><td>ENSMUSG00000079853</td></tr>
<tr><td>chr6</td><td>131245309</td><td>131245331</td><td>+</td><td>3</td><td>GACCTCCTGCTCACTCATCT</td><td>TGG</td><td>|||||||-[|||||-|||||-]PAM</td><td>0</td><td>Exonic</td><td>Klra2</td><td>ENSMUSG00000030187</td></tr>
<tr><td>chr6</td><td>129874890</td><td>129874912</td><td>+</td><td>3</td><td>GACCTCCTGCTCACTCATCA</td><td>TGG</td><td>|||||||-[|||||-|||||-]PAM</td><td>0</td><td>Exonic</td><td>Klra17</td><td>ENSMUSG00000014543</td></tr>
<tr><td>chr3</td><td>108364634</td><td>108364656</td><td>-</td><td>3</td><td>GACCTCCAGCTCATTCAACA</td><td>GGG</td><td>|||||||-[|||||||||-|-]PAM</td><td>255</td><td>Intergenic</td><td>Mybphl</td><td>ENSMUSG00000068745</td></tr>
<tr><td>chr1</td><td>135325183</td><td>135325205</td><td>+</td><td>4</td><td>GCCCTCGGGCTCATTCAACC</td><td>GGG</td><td>|-||||-|[|||||||||-|-]PAM</td><td>0</td><td>Exonic</td><td>Lmod1</td><td>ENSMUSG00000048096</td></tr>
<tr><td>chr5</td><td>70880265</td><td>70880287</td><td>+</td><td>4</td><td>CACCTCCAGCTCATTCATTC</td><td>AGG</td><td>-||||||-[||||||||||--]PAM</td><td>37648</td><td>Intergenic</td><td>Gabrg1</td><td>ENSMUSG00000001260</td></tr>
</table>

Species: Mouse (Mus musculus GRCm38/mm10)
Input: CCTCTGGCTCACTCATCGTGGGA
PAM: NGG
Target site length: 20
Core length: 12
Core MM: 2
Total MM: 4

T1 TCCCACGATGAGTGAGCCAGAGG 1000 CRISPRater score 0.601786315

<table>
<tr><th>Chromosome</th><th>start</th><th>end</th><th>strand</th><th>MM</th><th>target_seq</th><th>PAM</th><th>alignment</th><th>distance</th><th>position</th><th>gene name</th><th>gene id</th></tr>
<tr><td>chr6</td><td>130335784</td><td>130335806</td><td>-</td><td>0</td><td>TCCCACGATGAGTGAGCCAG</td><td>AGG</td><td>||||||||[||||||||||||]PAM</td><td>0</td><td>Exonic</td><td>Klra3</td><td>ENSMUSG00000067591</td></tr>
<tr><td>chr5</td><td>30405733</td><td>30405755</td><td>-</td><td>4</td><td>ACAAACGGTGAGTGAGCCAG</td><td>AGG</td><td>-|--|||-[||||||||||||]PAM</td><td>0</td><td>Exonic</td><td>Otof</td><td>ENSMUSG00000062372</td></tr>
<tr><td>chr2</td><td>153196734</td><td>153196756</td><td>-</td><td>3</td><td>TCACAGGACGAGTGAGCCAG</td><td>AGG</td><td>||-||-||[-|||||||||||]PAM</td><td>1169</td><td>Intronic</td><td>Tm9sf4</td><td>ENSMUSG00000068040</td></tr>
<tr><td>chr4</td><td>21065300</td><td>21065322</td><td>-</td><td>4</td><td>GACCAAGTTGAGTGAGCCAG</td><td>AGG</td><td>--|||-|-[||||||||||||]PAM</td><td>54360</td><td>Intergenic</td><td>Gm11871</td><td>ENSMUSG00000083315</td></tr>
<tr><td>chr18</td><td>46613640</td><td>46613662</td><td>-</td><td>4</td><td>TAACACCAAGAGTGAGCCAG</td><td>TGG</td><td>|--|||-|[-|||||||||||]PAM</td><td>3415</td><td>Intergenic</td><td>Eif1a</td><td>ENSMUSG00000057561</td></tr>
<tr><td>chr1</td><td>60522779</td><td>60522801</td><td>-</td><td>4</td><td>TCACGGGACGAGTGAGCCAG</td><td>AGG</td><td>||-|--||[-|||||||||||]PAM</td><td>38</td><td>Intronic</td><td>Gm11575</td><td>ENSMUSG00000081562</td></tr>
<tr><td>chr2</td><td>30448154</td><td>30448176</td><td>-</td><td>4</td><td>TGCCATTAGGAGTGAGCCAG</td><td>GGG</td><td>|-|||--|[-|||||||||||]PAM</td><td>348</td><td>Intergenic</td><td>Ppp2r4</td><td>ENSMUSG00000039515</td></tr>
<tr><td>chr4</td><td>40372173</td><td>40372195</td><td>+</td><td>4</td><td>TCACTTGATGAATGAGCCAG</td><td>CGG</td><td>||-|--||[|||-||||||||]PAM</td><td>54454</td><td>Intergenic</td><td>4930509K1</td><td>ENSMUSG00000087137</td></tr>
<tr><td>chr16</td><td>31433609</td><td>31433631</td><td>+</td><td>4</td><td>TTCCACGGGCAGTGAGCCAG</td><td>GGG</td><td>|-|||||-[--||||||||||]PAM</td><td>2460</td><td>Intronic</td><td>Bdh1</td><td>ENSMUSG00000046598</td></tr>
<tr><td>chr16</td><td>38469454</td><td>38469476</td><td>-</td><td>3</td><td>TTCCACGAAGAGAGAGCCAG</td><td>TGG</td><td>|-||||||[-|||-|||||||]PAM</td><td>4390</td><td>Intronic</td><td>Cd80</td><td>ENSMUSG00000075122</td></tr>
<tr><td>chr11</td><td>70073397</td><td>70073419</td><td>-</td><td>4</td><td>TCCCAGCACTAGTGAGCCAG</td><td>AGG</td><td>|||||--|[--||||||||||]PAM</td><td>15503</td><td>Intergenic</td><td>Asgr1</td><td>ENSMUSG00000020884</td></tr>
<tr><td>chr6</td><td>129911446</td><td>129911468</td><td>-</td><td>1</td><td>TCCCACGATGAGTGAACCAG</td><td>AGG</td><td>||||||||[|||||||-||||]PAM</td><td>0</td><td>Exonic</td><td>Klra5</td><td>ENSMUSG00000030173</td></tr>
<tr><td>chr10</td><td>57477333</td><td>57477355</td><td>+</td><td>4</td><td>TACAAAGATGAGTCAGCCAG</td><td>AGG</td><td>|-|-|-||[|||||-||||||]PAM</td><td>1026</td><td>Intergenic</td><td>4930467K1</td><td>ENSMUSG00000097370</td></tr>
<tr><td>chr7</td><td>109987562</td><td>109987584</td><td>-</td><td>4</td><td>TCCCAGAATTTGTGAGCCAG</td><td>AGG</td><td>|||||--|[|--|||||||||]PAM</td><td>494</td><td>Intergenic</td><td>Gm24842</td><td>ENSMUSG00000089306</td></tr>
<tr><td>chr13</td><td>47059919</td><td>47059941</td><td>+</td><td>4</td><td>TCCAGCGAAGAGAGAGCCAG</td><td>AGG</td><td>|||--|||[-|||-|||||||]PAM</td><td>812</td><td>Intronic</td><td>Kdm1b</td><td>ENSMUSG00000038080</td></tr>
<tr><td>chr4</td><td>12406133</td><td>12406155</td><td>-</td><td>4</td><td>TCCATCTATGAGTTAGCCAG</td><td>TGG</td><td>|||--|-|[|||||-||||||]PAM</td><td>91550</td><td>Intergenic</td><td>Gm11846</td><td>ENSMUSG00000085909</td></tr>
<tr><td>chr10</td><td>22496948</td><td>22496970</td><td>-</td><td>3</td><td>TCCCACTATGTGAGAGCCAG</td><td>AGG</td><td>||||||-|[||-|-|||||||]PAM</td><td>43428</td><td>Intergenic</td><td>Gm26585</td><td>ENSMUSG00000097473</td></tr>
<tr><td>chr10</td><td>22564519</td><td>22564541</td><td>+</td><td>3</td><td>TCCCACCATGTGAGAGCCAG</td><td>AGG</td><td>||||||-|[||-|-|||||||]PAM</td><td>80470</td><td>Intergenic</td><td>Slc2a12</td><td>ENSMUSG00000037490</td></tr>
<tr><td>chrX</td><td>18162016</td><td>18162038</td><td>-</td><td>4</td><td>CTCCACGAGGAGTGCGCCAG</td><td>AGG</td><td>--||||||[-|||||-|||||]PAM</td><td>537</td><td>Intergenic</td><td>Kdm6a</td><td>ENSMUSG00000037369</td></tr>
<tr><td>chr2</td><td>129087670</td><td>129087692</td><td>+</td><td>4</td><td>TCCCAGTATTAGGGAGCCAG</td><td>AGG</td><td>|||||--|[|-||-|||||||]PAM</td><td>1302</td><td>Intronic</td><td>Ttl</td><td>ENSMUSG00000027394</td></tr>
<tr><td>chr5</td><td>53512094</td><td>53512116</td><td>+</td><td>4</td><td>TGGAACGATGAGTGAACCAG</td><td>TGG</td><td>|---||||[|||||||-||||]PAM</td><td>7144</td><td>Intronic</td><td>Gm10441</td><td>ENSMUSG00000106807</td></tr>
<tr><td>chr19</td><td>40225125</td><td>40225147</td><td>-</td><td>4</td><td>TCCCAGCATGTGGGAGCCAG</td><td>AGG</td><td>|||||--|[||-|-|||||||]PAM</td><td>1769</td><td>Intronic</td><td>Pdlim1</td><td>ENSMUSG00000055044</td></tr>
<tr><td>chr14</td><td>8535671</td><td>8535693</td><td>+</td><td>3</td><td>TCCCAGGCTGAGTGACCCAG</td><td>TGG</td><td>|||||-|-[|||||||-||||]PAM</td><td>825</td><td>Intronic</td><td>4930452B0</td><td>ENSMUSG00000021747</td></tr>
<tr><td>chr10</td><td>33355638</td><td>33355660</td><td>+</td><td>4</td><td>TACCATGATGAATTAGCCAG</td><td>TGG</td><td>|-|||-||[|||-|-||||||]PAM</td><td>8318</td><td>Intronic</td><td>Trdn</td><td>ENSMUSG00000019787</td></tr>
<tr><td>chr7</td><td>40978490</td><td>40978512</td><td>-</td><td>4</td><td>TCCCAACATGAAGGAGCCAG</td><td>AGG</td><td>|||||--|[|||--|||||||]PAM</td><td>3952</td><td>Intergenic</td><td>Gm29293</td><td>ENSMUSG00000099768</td></tr>
<tr><td>chr4</td><td>105870422</td><td>105870444</td><td>+</td><td>4</td><td>TCCTAAGATCAGTGGGCCAG</td><td>GGG</td><td>|||-|-||[|-||||-|||||]PAM</td><td>75711</td><td>Intergenic</td><td>Gm12728</td><td>ENSMUSG00000083780</td></tr>
<tr><td>chr10</td><td>130129335</td><td>130129357</td><td>-</td><td>4</td><td>TCCTAAGATGACTCAGCCAG</td><td>TGG</td><td>|||-|-||[|||-|-||||||]PAM</td><td>2280</td><td>Intergenic</td><td>Olfr824</td><td>ENSMUSG00000095804</td></tr>
<tr><td>chr5</td><td>5031018</td><td>5031040</td><td>+</td><td>4</td><td>TCCCACTGTCAGTAAGCCAG</td><td>TGG</td><td>||||||--[|-|||-||||||]PAM</td><td>5386</td><td>Intronic</td><td>Cdk14</td><td>ENSMUSG00000028926</td></tr>
<tr><td>chr7</td><td>36809307</td><td>36809329</td><td>-</td><td>4</td><td>ACCCACTCTGAGTGACCCAG</td><td>GGG</td><td>-|||||--[|||||||-||||]PAM</td><td>35754</td><td>Intergenic</td><td>Tshz3</td><td>ENSMUSG00000021217</td></tr>
<tr><td>chr4</td><td>135743982</td><td>135744004</td><td>+</td><td>4</td><td>TGCCAAGATGAATGTGCCAG</td><td>TGG</td><td>|-|||-||[|||-||-|||||]PAM</td><td>682</td><td>Intronic</td><td>Il22ra1</td><td>ENSMUSG00000037157</td></tr>
<tr><td>chr16</td><td>37048695</td><td>37048717</td><td>-</td><td>4</td><td>TGCCACGGTGAGAAAGCCAG</td><td>AGG</td><td>|-|||||-[||||--||||||]PAM</td><td>0</td><td>Exonic</td><td>Polq</td><td>ENSMUSG00000034206</td></tr>
<tr><td>chrX</td><td>78583753</td><td>78583775</td><td>-</td><td>4</td><td>GCTCACGGTGAGTGAGGCAG</td><td>CGG</td><td>-|-||||-[||||||||-|||]PAM</td><td>0</td><td>Exonic</td><td>Prrg1</td><td>ENSMUSG00000047996</td></tr>
<tr><td>chr7</td><td>98118595</td><td>98118617</td><td>-</td><td>4</td><td>TACCAGCATGAGTGAGGCAG</td><td>GGG</td><td>|-|||--|[||||||||-|||]PAM</td><td>659</td><td>Intronic</td><td>Myo7a</td><td>ENSMUSG00000030761</td></tr>
<tr><td>chr8</td><td>110743814</td><td>110743836</td><td>+</td><td>3</td><td>TCCCAGGAGGAGTGAGACAG</td><td>AGG</td><td>|||||-||[-|||||||-|||]PAM</td><td>1035</td><td>Intronic</td><td>Il34</td><td>ENSMUSG00000031750</td></tr>
<tr><td>chr5</td><td>38220071</td><td>38220093</td><td>+</td><td>4</td><td>TCACGCGGTGAGTGAGGCAG</td><td>GGG</td><td>||-|-||-[||||||||-|||]PAM</td><td>230</td><td>Intronic</td><td>Zbtb49</td><td>ENSMUSG00000029127</td></tr>
<tr><td>chr15</td><td>37333366</td><td>37333388</td><td>-</td><td>4</td><td>TCCCAGGCTGAGCAAGCCAG</td><td>GGG</td><td>|||||-|-[||||--||||||]PAM</td><td>0</td><td>Exonic</td><td>Grhl2</td><td>ENSMUSG00000022286</td></tr>
<tr><td>chr7</td><td>105150497</td><td>105150519</td><td>-</td><td>4</td><td>CCCCACCATGAGGGGGCCAG</td><td>TGG</td><td>-|||||-|[||||-|-|||||]PAM</td><td>6188</td><td>Intergenic</td><td>Olfr683</td><td>ENSMUSG00000044120</td></tr>
<tr><td>chr2</td><td>13319917</td><td>13319939</td><td>+</td><td>4</td><td>TCCTATGATGGGTGATCCAG</td><td>GGG</td><td>|||-|-||[||-||||-||||]PAM</td><td>0</td><td>Exonic</td><td>Cubn</td><td>ENSMUSG00000026726</td></tr>
<tr><td>chr19</td><td>39101340</td><td>39101362</td><td>+</td><td>4</td><td>CCCCATGAAGAGTGAGTCAG</td><td>AGG</td><td>-||||-||[-|||||||-|||]PAM</td><td>7396</td><td>Intergenic</td><td>Cyp2c65</td><td>ENSMUSG00000067231</td></tr>
<tr><td>chr17</td><td>13711547</td><td>13711569</td><td>-</td><td>3</td><td>TCCAAGGATGAGTGAGCAAG</td><td>TGG</td><td>|||-|-||[|||||||||-||]PAM</td><td>875</td><td>Intronic</td><td>Tcte2</td><td>ENSMUSG00000038347</td></tr>
<tr><td>chr4</td><td>29110840</td><td>29110862</td><td>+</td><td>3</td><td>TCCAATGATGAGTGAGCAAG</td><td>AGG</td><td>|||-|-||[|||||||||-||]PAM</td><td>15084</td><td>Intergenic</td><td>Gm11918</td><td>ENSMUSG00000080911</td></tr>
<tr><td>chr6</td><td>55461608</td><td>55461630</td><td>+</td><td>4</td><td>TCCCCAGGTGAGTGAGGCAG</td><td>GGG</td><td>||||--|-[||||||||-|||]PAM</td><td>0</td><td>Exonic</td><td>Adcyap1r1</td><td>ENSMUSG00000029778</td></tr>
<tr><td>chr7</td><td>9208519</td><td>9208541</td><td>+</td><td>4</td><td>TCCCCTGTTGAGTGAGTCAG</td><td>AGG</td><td>||||--|-[||||||||-|||]PAM</td><td>1427</td><td>Intronic</td><td>Vmn2r37</td><td>ENSMUSG00000066828</td></tr>
<tr><td>chr7</td><td>8473883</td><td>8473905</td><td>+</td><td>4</td><td>TCCCCTGTTGAGTGAGTCAG</td><td>AGG</td><td>||||--|-[||||||||-|||]PAM</td><td>1428</td><td>Intronic</td><td>Vmn2r45</td><td>ENSMUSG00000090662</td></tr>
<tr><td>chr7</td><td>7674240</td><td>7674262</td><td>+</td><td>4</td><td>TCCCCTGTTGAGTGAGTCAG</td><td>AGG</td><td>||||--|-[||||||||-|||]PAM</td><td>1428</td><td>Intronic</td><td>Vmn2r34</td><td>ENSMUSG00000070841</td></tr>
<tr><td>chr7</td><td>9550112</td><td>9550134</td><td>+</td><td>4</td><td>TCCCCTGTTGAGTGAGTCAG</td><td>AGG</td><td>||||--|-[||||||||-|||]PAM</td><td>1497</td><td>Intronic</td><td>Gm10302</td><td>ENSMUSG00000096294</td></tr>
<tr><td>chr7</td><td>9819183</td><td>9819205</td><td>+</td><td>4</td><td>TCCCCTGTTGAGTGAGTCAG</td><td>AGG</td><td>||||--|-[||||||||-|||]PAM</td><td>1497</td><td>Intronic</td><td>Vmn2r47</td><td>ENSMUSG00000094107</td></tr>
<tr><td>chr7</td><td>8012741</td><td>8012763</td><td>+</td><td>4</td><td>TCCCCTGTTGAGTGAGTCAG</td><td>AGG</td><td>||||--|-[||||||||-|||]PAM</td><td>1499</td><td>Intronic</td><td>Gm4073</td><td>ENSMUSG00000094736</td></tr>
<tr><td>chr7</td><td>8803539</td><td>8803561</td><td>+</td><td>4</td><td>TCCCCTGTTGAGTGAGTCAG</td><td>AGG</td><td>||||--|-[||||||||-|||]PAM</td><td>1647</td><td>Intronic</td><td>Vmn2r-ps4</td><td>ENSMUSG00000096707</td></tr>
<tr><td>chr8</td><td>54638172</td><td>54638194</td><td>+</td><td>4</td><td>GCTCACGCTGAGTGAGCAAG</td><td>TGG</td><td>-|-||||-[|||||||||-||]PAM</td><td>93</td><td>Intronic</td><td>Wdr17</td><td>ENSMUSG00000039375</td></tr>
<tr><td>chr2</td><td>155715329</td><td>155715351</td><td>+</td><td>4</td><td>ACCCACGGTCAGTGAGTCAG</td><td>GGG</td><td>-||||||-[|-||||||-|||]PAM</td><td>659</td><td>Intronic</td><td>Edem2</td><td>ENSMUSG00000038312</td></tr>
<tr><td>chr14</td><td>28259265</td><td>28259287</td><td>+</td><td>4</td><td>TCCCAGCACGAGTGAGGCAG</td><td>AGG</td><td>|||||--|[-|||||||-|||]PAM</td><td>12291</td><td>Intronic</td><td>Erc2</td><td>ENSMUSG00000040640</td></tr>
<tr><td>chr12</td><td>68549987</td><td>68550009</td><td>-</td><td>4</td><td>TCCCCCGCCGAGTGAGTCAG</td><td>AGG</td><td>||||-||-[-|||||||-|||]PAM</td><td>50961</td><td>Intergenic</td><td>n-R5s61</td><td>ENSMUSG00000084660</td></tr>
<tr><td>chr9</td><td>99423693</td><td>99423715</td><td>+</td><td>4</td><td>TCCAATGATGAGGGACCCAG</td><td>TGG</td><td>|||-|-||[||||-||-||||]PAM</td><td>2062</td><td>Intronic</td><td>Mras</td><td>ENSMUSG00000032470</td></tr>
<tr><td>chr18</td><td>67847794</td><td>67847816</td><td>-</td><td>4</td><td>TCCCAGTATTAGTGAGGCAG</td><td>AGG</td><td>|||||--|[|-||||||-|||]PAM</td><td>212</td><td>Intronic</td><td>Cep192</td><td>ENSMUSG00000024542</td></tr>
<tr><td>chr6</td><td>125039264</td><td>125039286</td><td>+</td><td>4</td><td>TCCCAGCATTAGTGAGGCAG</td><td>AGG</td><td>|||||--|[|-||||||-|||]PAM</td><td>474</td><td>Intronic</td><td>Ing4</td><td>ENSMUSG00000030330</td></tr>
<tr><td>chr7</td><td>97783585</td><td>97783607</td><td>+</td><td>4</td><td>TCCCAGCATTAGTGAGGCAG</td><td>AGG</td><td>|||||--|[|-||||||-|||]PAM</td><td>4934</td><td>Intergenic</td><td>Pak1</td><td>ENSMUSG00000030774</td></tr>
<tr><td>chr6</td><td>130025204</td><td>130025226</td><td>-</td><td>2</td><td>TCCCACAATGAGTGAGCCGG</td><td>AGG</td><td>||||||-|[||||||||||-|]PAM</td><td>0</td><td>Exonic</td><td>Klra6</td><td>ENSMUSG00000061769</td></tr>
<tr><td>chr13</td><td>67831551</td><td>67831573</td><td>-</td><td>3</td><td>TCCCAAGATGAGGGAGGCAG</td><td>AGG</td><td>|||||-||[||||-|||-|||]PAM</td><td>566</td><td>Intergenic</td><td>4930525G2</td><td>ENSMUSG00000097276</td></tr>
<tr><td>chr13</td><td>113506429</td><td>113506451</td><td>+</td><td>4</td><td>TCCCAGCATGTGTGAGGCAG</td><td>AGG</td><td>|||||--|[||-|||||-|||]PAM</td><td>85729</td><td>Intergenic</td><td>Snx18</td><td>ENSMUSG00000042364</td></tr>
<tr><td>chr17</td><td>30740177</td><td>30740199</td><td>-</td><td>4</td><td>TCCAACAATGACTGAGACAG</td><td>TGG</td><td>|||-||-|[|||-||||-|||]PAM</td><td>422</td><td>Intronic</td><td>Dnah8</td><td>ENSMUSG00000033826</td></tr>
<tr><td>chr2</td><td>51325399</td><td>51325421</td><td>+</td><td>4</td><td>GCCCACAATAAGTGAGCAAG</td><td>AGG</td><td>-|||||-|[|-|||||||-||]PAM</td><td>29454</td><td>Intergenic</td><td>Gm13497</td><td>ENSMUSG00000086349</td></tr>
<tr><td>chr16</td><td>29628448</td><td>29628470</td><td>-</td><td>4</td><td>TCCCAGCATGACTGAGGCAG</td><td>AGG</td><td>|||||--|[|||-||||-|||]PAM</td><td>227</td><td>Intronic</td><td>Opa1</td><td>ENSMUSG00000038084</td></tr>
<tr><td>chr10</td><td>80339602</td><td>80339624</td><td>-</td><td>4</td><td>TCCCAGCATGAGGGAGGCAG</td><td>AGG</td><td>|||||--|[||||-|||-|||]PAM</td><td>25</td><td>Intronic</td><td>Adamtsl5</td><td>ENSMUSG00000043822</td></tr>
<tr><td>chr2</td><td>131180761</td><td>131180783</td><td>+</td><td>4</td><td>TCCCAGCATGAGGGAGACAG</td><td>AGG</td><td>|||||--|[||||-|||-|||]PAM</td><td>694</td><td>Intronic</td><td>Cenpb</td><td>ENSMUSG00000068267</td></tr>
<tr><td>chr9</td><td>78490625</td><td>78490647</td><td>+</td><td>4</td><td>TCCCAACATGAGGGAGACAG</td><td>AGG</td><td>|||||--|[||||-|||-|||]PAM</td><td>1474</td><td>Intergenic</td><td>Eef1a1</td><td>ENSMUSG00000037742</td></tr>
<tr><td>chrX</td><td>108827072</td><td>108827094</td><td>-</td><td>4</td><td>TCCCAGAATGAGGGAGGCAG</td><td>AGG</td><td>|||||--|[||||-|||-|||]PAM</td><td>1577</td><td>Intronic</td><td>Brwd3</td><td>ENSMUSG00000063663</td></tr>
<tr><td>chr1</td><td>74253053</td><td>74253075</td><td>+</td><td>4</td><td>TCCCAGCATGAGGGAGGCAG</td><td>AGG</td><td>|||||--|[||||-|||-|||]PAM</td><td>1753</td><td>Intronic</td><td>Arpc2</td><td>ENSMUSG00000006304</td></tr>
<tr><td>chr14</td><td>12168701</td><td>12168723</td><td>-</td><td>4</td><td>TCCCAGCATGAGGGAGGCAG</td><td>AGG</td><td>|||||--|[||||-|||-|||]PAM</td><td>1824</td><td>Intronic</td><td>Ptprg</td><td>ENSMUSG00000021745</td></tr>
<tr><td>chr11</td><td>40291942</td><td>40291964</td><td>-</td><td>4</td><td>TCCCAGTATGAGGGAGGCAG</td><td>AGG</td><td>|||||--|[||||-|||-|||]PAM</td><td>2100</td><td>Intronic</td><td>Gm12132</td><td>ENSMUSG00000085301</td></tr>
<tr><td>chr2</td><td>132103896</td><td>132103918</td><td>+</td><td>4</td><td>TCCCAGCATGAGGGAGGCAG</td><td>AGG</td><td>|||||--|[||||-|||-|||]PAM</td><td>2170</td><td>Intronic</td><td>Slc23a2</td><td>ENSMUSG00000027340</td></tr>
<tr><td>chr9</td><td>98982506</td><td>98982528</td><td>+</td><td>4</td><td>TCCCAGCATGAGGGAGGCAG</td><td>AGG</td><td>|||||--|[||||-|||-|||]PAM</td><td>3845</td><td>Intergenic</td><td>Faim</td><td>ENSMUSG00000032463</td></tr>
<tr><td>chr18</td><td>20737865</td><td>20737887</td><td>+</td><td>4</td><td>TCCCAGCATGAGGGAGGCAG</td><td>AGG</td><td>|||||--|[||||-|||-|||]PAM</td><td>7466</td><td>Intronic</td><td>B4galt6</td><td>ENSMUSG00000056124</td></tr>
<tr><td>chr10</td><td>12777550</td><td>12777572</td><td>+</td><td>4</td><td>TCCCAGCATGAGAGAGACAG</td><td>AGG</td><td>|||||--|[||||-|||-|||]PAM</td><td>12218</td><td>Intronic</td><td>Utrn</td><td>ENSMUSG00000019820</td></tr>
<tr><td>chr15</td><td>25604029</td><td>25604051</td><td>-</td><td>4</td><td>TCCCAGCATGAGGGAGGCAG</td><td>AGG</td><td>|||||--|[||||-|||-|||]PAM</td><td>18474</td><td>Intergenic</td><td>Myo10</td><td>ENSMUSG00000022272</td></tr>
<tr><td>chr9</td><td>7606218</td><td>7606240</td><td>+</td><td>4</td><td>TCCCAGCATGAGGGAGGCAG</td><td>GGG</td><td>|||||--|[||||-|||-|||]PAM</td><td>21991</td><td>Intergenic</td><td>Mmp20</td><td>ENSMUSG00000018620</td></tr>
<tr><td>chr19</td><td>22174128</td><td>22174150</td><td>-</td><td>4</td><td>TCCTACAATGAGTAAGTCAG</td><td>AGG</td><td>|||-||-|[|||||-||-|||]PAM</td><td>34827</td><td>Intronic</td><td>Trpm3</td><td>ENSMUSG00000052387</td></tr>
<tr><td>chr6</td><td>130191286</td><td>130191308</td><td>-</td><td>2</td><td>TCCCACGATGAATGAGCCGG</td><td>AGG</td><td>||||||||[|||-||||||-|]PAM</td><td>0</td><td>Exonic</td><td>Klra9</td><td>ENSMUSG00000033024</td></tr>
<tr><td>chr1</td><td>27231457</td><td>27231479</td><td>+</td><td>3</td><td>TCCCACGTTGAGTGGGTCAG</td><td>AGG</td><td>|||||||-[||||||-|-|||]PAM</td><td>NA</td><td>Intergenic</td><td>NA</td><td>NA</td></tr>
<tr><td>chr7</td><td>48958467</td><td>48958489</td><td>+</td><td>4</td><td>TCCCCAGATGGGTGAGCCTG</td><td>GGG</td><td>||||--||[||-|||||||-|]PAM</td><td>584</td><td>Intronic</td><td>Nav2</td><td>ENSMUSG00000052512</td></tr>
<tr><td>chr7</td><td>117940935</td><td>117940957</td><td>-</td><td>4</td><td>TCTGATGATGAGTGAGCCAA</td><td>GGG</td><td>||--|-||[|||||||||||-]PAM</td><td>NA</td><td>Intergenic</td><td>NA</td><td>NA</td></tr>
<tr><td>chr1</td><td>168199676</td><td>168199698</td><td>-</td><td>4</td><td>TCCAAATATGAGTGAGCCAT</td><td>GGG</td><td>|||-|--|[|||||||||||-]PAM</td><td>3649</td><td>Intronic</td><td>Pbx1</td><td>ENSMUSG00000052534</td></tr>
<tr><td>chr11</td><td>97819247</td><td>97819269</td><td>-</td><td>4</td><td>CCCCAAGATGAGTGAGGGAG</td><td>AGG</td><td>-||||-||[||||||||--||]PAM</td><td>3465</td><td>Intronic</td><td>Lasp1</td><td>ENSMUSG00000038366</td></tr>
<tr><td>chr6</td><td>130281907</td><td>130281929</td><td>-</td><td>2</td><td>TCCCACGATGAGTGAGCTGG</td><td>AGG</td><td>||||||||[|||||||||--|]PAM</td><td>0</td><td>Exonic</td><td>Klra10</td><td>ENSMUSG00000072718</td></tr>
<tr><td>chr6</td><td>130231602</td><td>130231624</td><td>-</td><td>3</td><td>TCCCAAGATGAGTGAGCAGG</td><td>AGG</td><td>|||||-||[|||||||||--|]PAM</td><td>0</td><td>Exonic</td><td>Klra7</td><td>ENSMUSG00000067599</td></tr>
<tr><td>chr6</td><td>130306412</td><td>130306434</td><td>-</td><td>3</td><td>TCCCAAGATGAGTGAGCAGG</td><td>AGG</td><td>|||||-||[|||||||||--|]PAM</td><td>0</td><td>Exonic</td><td>Klra13-ps</td><td>ENSMUSG00000030178</td></tr>
<tr><td>chr6</td><td>130160733</td><td>130160755</td><td>-</td><td>3</td><td>TCCCAAGATGAGTGAGCAGG</td><td>AGG</td><td>|||||-||[|||||||||--|]PAM</td><td>0</td><td>Exonic</td><td>Klra14-ps</td><td>ENSMUSG00000072721</td></tr>
<tr><td>chr6</td><td>130128145</td><td>130128167</td><td>-</td><td>3</td><td>TCCCAAGATGAGTGAGCAGG</td><td>AGG</td><td>|||||-||[|||||||||--|]PAM</td><td>0</td><td>Exonic</td><td>Klra8</td><td>ENSMUSG00000089727</td></tr>
<tr><td>chr6</td><td>130380647</td><td>130380669</td><td>-</td><td>3</td><td>TCCCAAGATGAGTGAGCAGG</td><td>AGG</td><td>|||||-||[|||||||||--|]PAM</td><td>0</td><td>Exonic</td><td>Klra1</td><td>ENSMUSG00000079853</td></tr>
<tr><td>chr5</td><td>123235616</td><td>123235638</td><td>+</td><td>4</td><td>TCCCACCCTGAGTGTGCCAT</td><td>TGG</td><td>||||||--[||||||-||||-]PAM</td><td>920</td><td>Intronic</td><td>Psmd9</td><td>ENSMUSG00000029440</td></tr>
<tr><td>chr1</td><td>91087328</td><td>91087350</td><td>+</td><td>4</td><td>TCCTACCATGAGTGAACCAA</td><td>TGG</td><td>|||-||-|[|||||||-|||-]PAM</td><td>204</td><td>Intronic</td><td>Lrrfip1</td><td>ENSMUSG00000026305</td></tr>
<tr><td>chr6</td><td>129874892</td><td>129874914</td><td>-</td><td>4</td><td>ACCCATGATGAGTGAGCAGG</td><td>AGG</td><td>-||||-||[|||||||||--|]PAM</td><td>0</td><td>Exonic</td><td>Klra17</td><td>ENSMUSG00000014543</td></tr>
<tr><td>chr6</td><td>131245311</td><td>131245333</td><td>-</td><td>4</td><td>TTCCAAGATGAGTGAGCAGG</td><td>AGG</td><td>|-|||-||[|||||||||--|]PAM</td><td>0</td><td>Exonic</td><td>Klra2</td><td>ENSMUSG00000030187</td></tr>
<tr><td>chr18</td><td>9750381</td><td>9750403</td><td>+</td><td>4</td><td>TTCCAAGATGAGTGAGACAT</td><td>AGG</td><td>|-|||-||[||||||||-||-]PAM</td><td>657</td><td>Intronic</td><td>Gm7497</td><td>ENSMUSG00000098235</td></tr>
<tr><td>chr11</td><td>106531359</td><td>106531381</td><td>+</td><td>4</td><td>TGCCACCATGAGTGAGCCCA</td><td>GGG</td><td>|-||||-|[||||||||||--]PAM</td><td>1998</td><td>Intronic</td><td>Tex2</td><td>ENSMUSG00000040548</td></tr>
<tr><td>chr17</td><td>33484021</td><td>33484043</td><td>+</td><td>4</td><td>TCTCACAATGAGTGAGCCCC</td><td>AGG</td><td>||-|||-|[||||||||||--]PAM</td><td>40161</td><td>Intergenic</td><td>Adamts10</td><td>ENSMUSG00000024299</td></tr>
</table>

Species: Mouse (Mus musculus GRCm38/mm10)
Input: CCTGTCTCCTCAAGCCTCACCCG
PAM: NGG
Target site length: 20
Core length: 12
Core MM: 2
Total MM: 4

<table>
<tr><td>T1</td><td>CGGGTGAGGCTTGAGGAGACAGG</td><td>1000</td><td>CRISPRater score</td><td>0.7103423</td><td></td><td></td><td></td><td></td><td></td><td></td><td></td></tr>
<tr><th>Chromosome</th><th>start</th><th>end</th><th>strand</th><th>MM</th><th>target_seq</th><th>PAM</th><th>alignment</th><th>distance</th><th>position</th><th>gene name</th><th>gene id</th></tr>
<tr><td>chr6</td><td>130306342</td><td>130306364</td><td>-</td><td>0</td><td>CGGGTGAGGCTTGAGGAGAC</td><td>AGG</td><td>||||||||[|||||||||||||]PAM</td><td>0</td><td>Exonic</td><td>Klra13-ps</td><td>ENSMUSG00000030178</td></tr>
<tr><td>chr17</td><td>25483027</td><td>25483049</td><td>+</td><td>4</td><td>CTGTGGAAGCTTGAGGAGAC</td><td>AGG</td><td>|-|--||-[|||||||||||]PAM</td><td>6433</td><td>Intergenic</td><td>Tekt4</td><td>ENSMUSG00000024175</td></tr>
<tr><td>chr8</td><td>118975688</td><td>118975710</td><td>-</td><td>4</td><td>TGGCTGTGACTTGAGGAGAC</td><td>AGG</td><td>-||-||-|[-||||||||||]PAM</td><td>6820</td><td>Intronic</td><td>Cdh13</td><td>ENSMUSG00000031841</td></tr>
<tr><td>chr11</td><td>8670630</td><td>8670652</td><td>-</td><td>4</td><td>TGAGTGTGGTTTGAGGAGAC</td><td>TGG</td><td>-|-|||-|[|-|||||||||]PAM</td><td>6095</td><td>Intergenic</td><td>Tns3</td><td>ENSMUSG00000020422</td></tr>
<tr><td>chr16</td><td>87578815</td><td>87578837</td><td>+</td><td>4</td><td>GGGCTGACACTTGAGGAGAC</td><td>AGG</td><td>-||-|||-[-||||||||||]PAM</td><td>2293</td><td>Intronic</td><td>Map3k7cl</td><td>ENSMUSG00000025610</td></tr>
<tr><td>chr5</td><td>140683330</td><td>140683352</td><td>-</td><td>4</td><td>AGAGGGAGGCATGAGGAGAC</td><td>GGG</td><td>-|-|-|||[||-|||||||||]PAM</td><td>0</td><td>Exonic</td><td>Iqce</td><td>ENSMUSG00000036555</td></tr>
<tr><td>chr2</td><td>173578109</td><td>173578131</td><td>-</td><td>2</td><td>CGGGTGAAGCTGGAGGAGAC</td><td>TGG</td><td>|||||||-[|||-||||||||]PAM</td><td>0</td><td>Exonic</td><td>Ankrd60</td><td>ENSMUSG00000027517</td></tr>
<tr><td>chr4</td><td>136638205</td><td>136638227</td><td>-</td><td>4</td><td>GGGGGGGGGGTTGAGGAGAC</td><td>AGG</td><td>-|||-|-|[|-|||||||||]PAM</td><td>0</td><td>Exonic</td><td>Lactbl1</td><td>ENSMUSG00000070683</td></tr>
<tr><td>chr5</td><td>99764867</td><td>99764889</td><td>+</td><td>3</td><td>GGGGTGGGGCTGGAGGAGAC</td><td>TGG</td><td>-|||||-|[|||-||||||||]PAM</td><td>10894</td><td>Intergenic</td><td>Gm43248</td><td>ENSMUSG00000105218</td></tr>
<tr><td>chr8</td><td>89650899</td><td>89650921</td><td>-</td><td>4</td><td>GGCATGAGGCTGGAGGAGAC</td><td>AGG</td><td>-|--||||[|||-||||||||]PAM</td><td>16252</td><td>Intergenic</td><td>Gm24212</td><td>ENSMUSG00000088883</td></tr>
<tr><td>chr1</td><td>58338498</td><td>58338520</td><td>-</td><td>4</td><td>CAGGTCTGGGTTGAGGAGAC</td><td>AGG</td><td>|-|||--|[|-|||||||||]PAM</td><td>939</td><td>Intronic</td><td>Aox2</td><td>ENSMUSG00000079554</td></tr>
<tr><td>chr3</td><td>95629789</td><td>95629811</td><td>+</td><td>4</td><td>CTGGATAGGCCTGAGGAGAC</td><td>AGG</td><td>|-||--||[||-|||||||||]PAM</td><td>0</td><td>Exonic</td><td>Ensa</td><td>ENSMUSG00000038619</td></tr>
<tr><td>chr12</td><td>67983863</td><td>67983885</td><td>-</td><td>4</td><td>GGGATGAGCATTGAGGAGAC</td><td>AGG</td><td>-||-||||[--||||||||||]PAM</td><td>NA</td><td>Intergenic</td><td>NA</td><td>NA</td></tr>
<tr><td>chr19</td><td>34001419</td><td>34001441</td><td>-</td><td>4</td><td>CGGGACATCCTTGAGGAGAC</td><td>AGG</td><td>||||--|-[-||||||||||]PAM</td><td>6813</td><td>Intergenic</td><td>Lipk</td><td>ENSMUSG00000024771</td></tr>
<tr><td>chr5</td><td>106684904</td><td>106684926</td><td>-</td><td>4</td><td>GGGGTGGTGATTGAGGAGAC</td><td>AGG</td><td>-|||||--[|-|||||||||]PAM</td><td>2891</td><td>Intronic</td><td>Zfp644</td><td>ENSMUSG00000049606</td></tr>
<tr><td>chr1</td><td>19560362</td><td>19560384</td><td>-</td><td>4</td><td>CTGGAGAAGCATGAGGAGAC</td><td>AGG</td><td>|-||-||-[||-|||||||||]PAM</td><td>46443</td><td>Intergenic</td><td>Gm24633</td><td>ENSMUSG00000084668</td></tr>
<tr><td>chr16</td><td>4273096</td><td>4273118</td><td>-</td><td>4</td><td>CATGTGAGCCCTGAGGAGAC</td><td>AGG</td><td>|--|||||[-|-|||||||||]PAM</td><td>14411</td><td>Intergenic</td><td>Adcy9</td><td>ENSMUSG00000005580</td></tr>
<tr><td>chr8</td><td>76198998</td><td>76199020</td><td>+</td><td>4</td><td>TGAGTGAGGACTGAGGAGAC</td><td>TGG</td><td>-|-|||||[|--|||||||||]PAM</td><td>75324</td><td>Intergenic</td><td>Gm22509</td><td>ENSMUSG00000064709</td></tr>
<tr><td>chr12</td><td>110549211</td><td>110549233</td><td>-</td><td>4</td><td>CAGGTCAGTCGTGAGGAGAC</td><td>TGG</td><td>|-|||-||[-|-|||||||||]PAM</td><td>1829</td><td>Intronic</td><td>Ppp2r5c</td><td>ENSMUSG00000017843</td></tr>
<tr><td>chr19</td><td>42118200</td><td>42118222</td><td>+</td><td>4</td><td>GGGGTAAAGCTAGAGGAGAC</td><td>TGG</td><td>-||||-|-[|||-||||||||]PAM</td><td>1571</td><td>Intronic</td><td>Pi4k2a</td><td>ENSMUSG00000025178</td></tr>
<tr><td>chr1</td><td>42542810</td><td>42542832</td><td>-</td><td>3</td><td>AGGGTGTGGCTTGTGGAGAC</td><td>AGG</td><td>-|||||-|[|||||-||||||]PAM</td><td>47147</td><td>Intergenic</td><td>Gm37047</td><td>ENSMUSG00000103106</td></tr>
<tr><td>chr2</td><td>93082643</td><td>93082665</td><td>+</td><td>4</td><td>GGGGTGTGACTGGAGGAGAC</td><td>AGG</td><td>-|||||-|[-||-||||||||]PAM</td><td>3277</td><td>Intergenic</td><td>Gm13802</td><td>ENSMUSG00000087559</td></tr>
<tr><td>chr19</td><td>57979994</td><td>57980016</td><td>+</td><td>4</td><td>AGGGTGAACCTGGAGGAGAC</td><td>TGG</td><td>-||||||-[-||-||||||||]PAM</td><td>44290</td><td>Intronic</td><td>Atrnl1</td><td>ENSMUSG00000054843</td></tr>
<tr><td>chr9</td><td>111346273</td><td>111346295</td><td>-</td><td>4</td><td>CAGCTGAGGCAAGAGGAGAC</td><td>AGG</td><td>|-|-||||[||--||||||||]PAM</td><td>760</td><td>Intronic</td><td>Trank1</td><td>ENSMUSG00000062296</td></tr>
<tr><td>chr12</td><td>104875666</td><td>104875688</td><td>+</td><td>4</td><td>GGGGAGGGGCTTGGGGAGAC</td><td>AGG</td><td>-|||-|-|[|||||-||||||]PAM</td><td>4142</td><td>Intergenic</td><td>4930408O1</td><td>ENSMUSG00000021101</td></tr>
<tr><td>chr12</td><td>38149666</td><td>38149688</td><td>-</td><td>3</td><td>CTGCTGAGGCTTGAGTAGAC</td><td>AGG</td><td>|-|-||||[|||||||-||||]PAM</td><td>2065</td><td>Intronic</td><td>Dgkb</td><td>ENSMUSG00000036095</td></tr>
<tr><td>chr6</td><td>54186775</td><td>54186797</td><td>-</td><td>4</td><td>TGGATGAGGGTTGGGGAGAC</td><td>TGG</td><td>-||-||||[|-|||-||||||]PAM</td><td>7229</td><td>Intronic</td><td>Chn2</td><td>ENSMUSG00000004633</td></tr>
<tr><td>chr1</td><td>57031596</td><td>57031618</td><td>+</td><td>4</td><td>CCTGTGAGGCTGCAGGAGAC</td><td>GGG</td><td>|--|||||[|||--|||||||]PAM</td><td>8715</td><td>Intronic</td><td>9130024F1</td><td>ENSMUSG00000087022</td></tr>
<tr><td>chr1</td><td>81554788</td><td>81554810</td><td>+</td><td>4</td><td>CAGGAGAGGGTTGTGGAGAC</td><td>TGG</td><td>|-||-|||[|-|||-||||||]PAM</td><td>0</td><td>Exonic</td><td>Gm6198</td><td>ENSMUSG00000101986</td></tr>
<tr><td>chr8</td><td>25819530</td><td>25819552</td><td>-</td><td>4</td><td>CAGCTGAGGCTCAAGGAGAC</td><td>AGG</td><td>|-|-||||[|||--|||||||]PAM</td><td>74</td><td>Intronic</td><td>Ash2l</td><td>ENSMUSG00000031575</td></tr>
<tr><td>chr17</td><td>12295120</td><td>12295142</td><td>-</td><td>4</td><td>TGGCTGAGGCCTGTGGAGAC</td><td>AGG</td><td>-||-||||[||-||-||||||]PAM</td><td>5894</td><td>Intronic</td><td>Gm24794</td><td>ENSMUSG00000077476</td></tr>
<tr><td>chr5</td><td>142773847</td><td>142773869</td><td>+</td><td>3</td><td>CGGGGGAGGCTCGGGGAGAC</td><td>GGG</td><td>||||-|||[|||-|-||||||]PAM</td><td>0</td><td>Exonic</td><td>Tnrc18</td><td>ENSMUSG00000039477</td></tr>
<tr><td>chr3</td><td>102481888</td><td>102481910</td><td>+</td><td>3</td><td>GTGGTGAGGCTTGAGGTGAC</td><td>CGG</td><td>--||||||[||||||||-|||]PAM</td><td>892</td><td>Intronic</td><td>Ngf</td><td>ENSMUSG00000027859</td></tr>
<tr><td>chr7</td><td>99884229</td><td>99884251</td><td>-</td><td>4</td><td>CAGGAGAGGTTTGATGAGAC</td><td>TGG</td><td>|-||-|||[|-||||-|||||]PAM</td><td>1752</td><td>Intronic</td><td>Xrra1</td><td>ENSMUSG00000035211</td></tr>
<tr><td>chr2</td><td>180531779</td><td>180531801</td><td>-</td><td>4</td><td>CTGGGGAGGCTGGCGGAGAC</td><td>TGG</td><td>|-||-|||[|||-|-||||||]PAM</td><td>6816</td><td>Intronic</td><td>Ntsr1</td><td>ENSMUSG00000027568</td></tr>
<tr><td>chr7</td><td>64050030</td><td>64050052</td><td>-</td><td>4</td><td>AGGGAGAGGCATGAAGAGAC</td><td>AGG</td><td>-|||-|||[||-|||-|||||]PAM</td><td>7550</td><td>Intergenic</td><td>Gm20670</td><td>ENSMUSG00000108620</td></tr>
<tr><td>chr8</td><td>27114261</td><td>27114283</td><td>-</td><td>4</td><td>AGCATGAGGCTTGAGGTGAC</td><td>TGG</td><td>-|--||||[||||||||-|||]PAM</td><td>32</td><td>Intronic</td><td>Adgra2</td><td>ENSMUSG00000031486</td></tr>
<tr><td>chr15</td><td>41554935</td><td>41554957</td><td>+</td><td>4</td><td>GGGGGGAGGCTTAGGGAGAC</td><td>TGG</td><td>-|||-|||[||||--||||||]PAM</td><td>19208</td><td>Intronic</td><td>Oxr1</td><td>ENSMUSG00000022307</td></tr>
<tr><td>chr6</td><td>86161156</td><td>86161178</td><td>-</td><td>4</td><td>CTGGAGAGGCTGGAAGAGAC</td><td>AGG</td><td>|-||-|||[|||-||-|||||]PAM</td><td>12962</td><td>Intergenic</td><td>RP23-304D</td><td>ENSMUSG00000107857</td></tr>
<tr><td>chr9</td><td>45572849</td><td>45572871</td><td>+</td><td>4</td><td>CTGGGGAGGCTGGAAGAGAC</td><td>AGG</td><td>|-||-|||[|||-||-|||||]PAM</td><td>61835</td><td>Intronic</td><td>Gm22069</td><td>ENSMUSG00000096202</td></tr>
<tr><td>chr10</td><td>111955202</td><td>111955224</td><td>+</td><td>4</td><td>AGGGTGTGGCTTTGGGAGAC</td><td>AGG</td><td>-|||||-|[||||--||||||]PAM</td><td>17440</td><td>Intergenic</td><td>Krr1</td><td>ENSMUSG00000063334</td></tr>
<tr><td>chr12</td><td>55563755</td><td>55563777</td><td>+</td><td>4</td><td>CCTGTGAAGCTTGAGGGGAC</td><td>TGG</td><td>|--||||-[||||||||-|||]PAM</td><td>1462</td><td>Intergenic</td><td>Aldoart2</td><td>ENSMUSG00000063129</td></tr>
<tr><td>chr16</td><td>11089336</td><td>11089358</td><td>-</td><td>4</td><td>AGGGTGTGGCCTGAGAAGAC</td><td>TGG</td><td>-|||||-|[||-||||-||||]PAM</td><td>767</td><td>Intronic</td><td>Txndc11</td><td>ENSMUSG00000022498</td></tr>
<tr><td>chr17</td><td>14795907</td><td>14795929</td><td>-</td><td>4</td><td>AGAGTGAGGCTTTAGTAGAC</td><td>AGG</td><td>-|-|||||[||||-||-||||]PAM</td><td>3903</td><td>Intergenic</td><td>Gm23352</td><td>ENSMUSG00000089053</td></tr>
<tr><td>chrX</td><td>155520148</td><td>155520170</td><td>+</td><td>4</td><td>AGGGTGGGACTTGAGGTGAC</td><td>TGG</td><td>-|||||-|[-|||||||-|||]PAM</td><td>15224</td><td>Intergenic</td><td>Gm8606</td><td>ENSMUSG00000083362</td></tr>
<tr><td>chr15</td><td>87200434</td><td>87200456</td><td>-</td><td>4</td><td>AGGCTGAGGCCTGAGGGGAC</td><td>AGG</td><td>-||-||||[||-|||||-|||]PAM</td><td>31218</td><td>Intergenic</td><td>n-R5s42</td><td>ENSMUSG00000064631</td></tr>
<tr><td>chr1</td><td>93260010</td><td>93260032</td><td>+</td><td>4</td><td>GGGGTGGGGGTTGAGGGGAC</td><td>AGG</td><td>-|||||-|[|-||||||-|||]PAM</td><td>203</td><td>Intronic</td><td>Sned1</td><td>ENSMUSG00000047793</td></tr>
<tr><td>chr5</td><td>110499266</td><td>110499288</td><td>-</td><td>4</td><td>GGGGTGGGGTTTGAGGTGAC</td><td>AGG</td><td>-|||||-|[|-||||||-|||]PAM</td><td>15635</td><td>Intergenic</td><td>A630023P1</td><td>ENSMUSG00000048215</td></tr>
<tr><td>chr14</td><td>89757706</td><td>89757728</td><td>-</td><td>4</td><td>TGTGTGAAGCTTGAGGAAAC</td><td>AGG</td><td>-|-||||-[||||||||-||]PAM</td><td>50770</td><td>Intergenic</td><td>Gm25415</td><td>ENSMUSG00000077408</td></tr>
<tr><td>chr6</td><td>55609582</td><td>55609604</td><td>+</td><td>4</td><td>CTGGTGATGCTTGCTGAGAC</td><td>TGG</td><td>|-|||||-[|||||--|||||]PAM</td><td>68218</td><td>Intergenic</td><td>Neurod6</td><td>ENSMUSG00000037984</td></tr>
<tr><td>chrX</td><td>91826167</td><td>91826189</td><td>-</td><td>4</td><td>AAGGTGAGGTTTGAGGAAAC</td><td>AGG</td><td>--||||||[|-|||||||-||]PAM</td><td>33309</td><td>Intergenic</td><td>Gm14782</td><td>ENSMUSG00000082807</td></tr>
<tr><td>chr11</td><td>64325618</td><td>64325640</td><td>+</td><td>3</td><td>CGTGTGAGGCTTGTGGTGAC</td><td>AGG</td><td>||-|||||[|||||-||-|||]PAM</td><td>NA</td><td>Intergenic</td><td>NA</td><td>NA</td></tr>
<tr><td>chr4</td><td>89774968</td><td>89774990</td><td>+</td><td>4</td><td>GGGGAGAGGCCTGAGGACAC</td><td>TGG</td><td>-|||-|||[||-||||||-||]PAM</td><td>80196</td><td>Intergenic</td><td>Dmrta1</td><td>ENSMUSG00000043753</td></tr>
<tr><td>chr19</td><td>8587852</td><td>8587874</td><td>-</td><td>4</td><td>AGGGTCAGGCCTGAGGAAAC</td><td>AGG</td><td>-||||-||[||-||||||-||]PAM</td><td>533</td><td>Intergenic</td><td>Gm6425</td><td>ENSMUSG00000086174</td></tr>
<tr><td>chr7</td><td>44581845</td><td>44581867</td><td>+</td><td>4</td><td>AGGGAGGGGCTTGAGGAGGC</td><td>AGG</td><td>-|||-|-|[||||||||||-|]PAM</td><td>44</td><td>Intronic</td><td>Napsa</td><td>ENSMUSG00000002204</td></tr>
<tr><td>chr10</td><td>80312459</td><td>80312481</td><td>+</td><td>4</td><td>CTGGAGGGGCTTGAGGAGGC</td><td>TGG</td><td>|-||-|-|[||||||||||-|]PAM</td><td>0</td><td>Exonic</td><td>Apc2</td><td>ENSMUSG00000020135</td></tr>
<tr><td>chr19</td><td>55863374</td><td>55863396</td><td>-</td><td>2</td><td>AGGGTGAGGCTTGAGGAGAG</td><td>AGG</td><td>-|||||||[||||||||||||-]PAM</td><td>22</td><td>Intronic</td><td>Tcf7l2</td><td>ENSMUSG00000024985</td></tr>
<tr><td>chr19</td><td>52001443</td><td>52001465</td><td>-</td><td>4</td><td>TGGCTGAGGCATGAGGAGCC</td><td>TGG</td><td>-||-||||[||-|||||||-|]PAM</td><td>NA</td><td>Intergenic</td><td>NA</td><td>NA</td></tr>
<tr><td>chrX</td><td>11212846</td><td>11212868</td><td>-</td><td>4</td><td>AGTTTGAGGCTTGAGGAGAG</td><td>AGG</td><td>-|--||||[|||||||||||-]PAM</td><td>7739</td><td>Intergenic</td><td>Gm14481</td><td>ENSMUSG00000080948</td></tr>
<tr><td>chr5</td><td>30455527</td><td>30455549</td><td>-</td><td>4</td><td>CAGGTGTGGCTTGTGGACAC</td><td>AGG</td><td>|-||||-|[|||||-|||-||]PAM</td><td>6171</td><td>Intronic</td><td>Otof</td><td>ENSMUSG00000062372</td></tr>
<tr><td>chr6</td><td>127365391</td><td>127365413</td><td>+</td><td>4</td><td>TGGGAGAGGCTCGAGGAGGC</td><td>AGG</td><td>-|||-|||[|||-||||||-|]PAM</td><td>2534</td><td>Intergenic</td><td>Gm43631</td><td>ENSMUSG00000107243</td></tr>
<tr><td>chr5</td><td>149358755</td><td>149358777</td><td>-</td><td>4</td><td>GGGGAGAGGCTCGAGGAGCC</td><td>GGG</td><td>-|||-|||[|||-||||||-|]PAM</td><td>8859</td><td>Intergenic</td><td>BC028471</td><td>ENSMUSG00000107271</td></tr>
<tr><td>chr4</td><td>43461524</td><td>43461546</td><td>+</td><td>4</td><td>CGGGAGAAGGTTGAGGAGCC</td><td>TGG</td><td>||||-||-[|-|||||||-|]PAM</td><td>6896</td><td>Intergenic</td><td>Cd72</td><td>ENSMUSG00000028459</td></tr>
<tr><td>chr17</td><td>35802820</td><td>35802842</td><td>-</td><td>4</td><td>AGGGGTAGGCTTGAGGAGAA</td><td>AGG</td><td>-|||--||[|||||||||||-]PAM</td><td>18842</td><td>Intergenic</td><td>Ier3</td><td>ENSMUSG00000003541</td></tr>
<tr><td>chr11</td><td>93000248</td><td>93000270</td><td>+</td><td>4</td><td>CTGGAAAGGCTTGAGGAGAA</td><td>TGG</td><td>|-||--||[|||||||||||-]PAM</td><td>89809</td><td>Intergenic</td><td>Gm25113</td><td>ENSMUSG00000087810</td></tr>
<tr><td>chr18</td><td>20524390</td><td>20524412</td><td>-</td><td>4</td><td>CTGGTGGGGCTTGAGACGAC</td><td>TGG</td><td>|-||||-|[|||||||--|||]PAM</td><td>147</td><td>Intronic</td><td>Dsg3</td><td>ENSMUSG00000056632</td></tr>
<tr><td>chr11</td><td>79042542</td><td>79042564</td><td>-</td><td>3</td><td>GGGGTGAGGCCTGAGGAGAG</td><td>AGG</td><td>-|||||||[||-||||||||-]PAM</td><td>632</td><td>Intronic</td><td>Ksr1</td><td>ENSMUSG00000018334</td></tr>
<tr><td>chr5</td><td>114950896</td><td>114950918</td><td>+</td><td>4</td><td>GGGGTGGGGCTTCAGGAGGC</td><td>AGG</td><td>-|||||-|[||||-|||||-|]PAM</td><td>8</td><td>Intronic</td><td>Hnf1a</td><td>ENSMUSG00000029556</td></tr>
<tr><td>chr2</td><td>38357136</td><td>38357158</td><td>+</td><td>4</td><td>CGCGGGAGTCTTGAGGAGAT</td><td>CGG</td><td>||-|-|||[-||||||||||-]PAM</td><td>2284</td><td>Intronic</td><td>Lhx2</td><td>ENSMUSG00000000247</td></tr>
<tr><td>chr8</td><td>120309730</td><td>120309752</td><td>-</td><td>4</td><td>ATGGTGAGGCGTGAGGAGAG</td><td>GGG</td><td>--||||||[||-||||||||-]PAM</td><td>44238</td><td>Intronic</td><td>Gse1</td><td>ENSMUSG00000031822</td></tr>
<tr><td>chr13</td><td>8641366</td><td>8641388</td><td>+</td><td>4</td><td>AGGGGGAGGCTTGAGTAAAC</td><td>AGG</td><td>-|||-|||[|||||||-|-||]PAM</td><td>31017</td><td>Intronic</td><td>Adarb2</td><td>ENSMUSG00000052551</td></tr>
<tr><td>chr1</td><td>186795048</td><td>186795070</td><td>+</td><td>3</td><td>CGGGGGAGGCTGGAGGAGAG</td><td>AGG</td><td>||||-|||[|||-|||||||-]PAM</td><td>42127</td><td>Intergenic</td><td>A430105J0</td><td>ENSMUSG00000097784</td></tr>
<tr><td>chr12</td><td>108644816</td><td>108644838</td><td>+</td><td>4</td><td>CCAGTGAGGCTTGAAGAGGC</td><td>AGG</td><td>|--|||||[|||||||-|||-|]PAM</td><td>3477</td><td>Intronic</td><td>Evl</td><td>ENSMUSG00000021262</td></tr>
<tr><td>chr12</td><td>87089136</td><td>87089158</td><td>+</td><td>4</td><td>CAGGTGGGGCATGAGGAGAG</td><td>AGG</td><td>|-||||-|[||-||||||||-]PAM</td><td>68</td><td>Intronic</td><td>Tmem63c</td><td>ENSMUSG00000034145</td></tr>
<tr><td>chr7</td><td>64196394</td><td>64196416</td><td>-</td><td>4</td><td>CAGGTGATGCTTGAGAAAAC</td><td>AGG</td><td>|-|||||-[||||||-|-||]PAM</td><td>861</td><td>Intronic</td><td>Trpm1</td><td>ENSMUSG00000030523</td></tr>
<tr><td>chr8</td><td>84234845</td><td>84234867</td><td>-</td><td>4</td><td>TGGGGGAGGCTGGAGGAGAG</td><td>AGG</td><td>-|||-|||[|||-|||||||-]PAM</td><td>36</td><td>Intronic</td><td>Zswim4</td><td>ENSMUSG00000035671</td></tr>
<tr><td>chr8</td><td>11305000</td><td>11305022</td><td>-</td><td>4</td><td>CAGGTGGGGCTGGAGGAGAG</td><td>CGG</td><td>|-||||-|[|||-|||||||-]PAM</td><td>7494</td><td>Intronic</td><td>Col4a1</td><td>ENSMUSG00000031502</td></tr>
<tr><td>chr2</td><td>92950032</td><td>92950054</td><td>+</td><td>4</td><td>GGGGTGGGGCTTGAGGGCAC</td><td>TGG</td><td>-|||||-|[||||||||--||]PAM</td><td>1248</td><td>Intronic</td><td>Syt13</td><td>ENSMUSG00000027220</td></tr>
<tr><td>chr14</td><td>121798660</td><td>121798682</td><td>-</td><td>4</td><td>TGGGTGTGGCTTGAGTAGCC</td><td>GGG</td><td>-|||||-|[|||||||-||-|]PAM</td><td>926</td><td>Intergenic</td><td>Dock9</td><td>ENSMUSG00000025558</td></tr>
<tr><td>chr7</td><td>68173052</td><td>68173074</td><td>+</td><td>4</td><td>CCGGTGTGGCTTGCGGAGAG</td><td>TGG</td><td>|-||||-|[|||||-|||||-]PAM</td><td>157</td><td>Intronic</td><td>Igf1r</td><td>ENSMUSG00000005533</td></tr>
<tr><td>chr17</td><td>50129575</td><td>50129597</td><td>+</td><td>4</td><td>CAGGAGAGGCTTGAAGAGAG</td><td>AGG</td><td>|-||-|||[||||||-||||-]PAM</td><td>22319</td><td>Intronic</td><td>Rftn1</td><td>ENSMUSG00000039316</td></tr>
<tr><td>chr5</td><td>107338167</td><td>107338189</td><td>-</td><td>4</td><td>TGGGGGAGGCTTGAGGGGGC</td><td>AGG</td><td>-|||-|||[||||||||-|-|]PAM</td><td>3111</td><td>Intronic</td><td>Brdt</td><td>ENSMUSG00000029279</td></tr>
<tr><td>chr15</td><td>79308594</td><td>79308616</td><td>+</td><td>3</td><td>CTGGTGAGGCTTGAGGAAGC</td><td>AGG</td><td>|-||||||[||||||||||--|]PAM</td><td>53</td><td>Intronic</td><td>Pla2g6</td><td>ENSMUSG00000042632</td></tr>
<tr><td>chr12</td><td>112062945</td><td>112062967</td><td>+</td><td>4</td><td>CACGTGAGGCTTGAGGAAGC</td><td>AGG</td><td>|--|||||[||||||||||--|]PAM</td><td>187</td><td>Intronic</td><td>Tdrd9</td><td>ENSMUSG00000054003</td></tr>
<tr><td>chr2</td><td>132307308</td><td>132307330</td><td>+</td><td>4</td><td>CAGCTGAGGCTTGAGGAAGC</td><td>TGG</td><td>|-|-||||[||||||||||--|]PAM</td><td>0</td><td>Exonic</td><td>Cds2</td><td>ENSMUSG00000058793</td></tr>
<tr><td>chr10</td><td>23371599</td><td>23371621</td><td>+</td><td>4</td><td>CCAGTGAGGCTTGAGGTGAA</td><td>GGG</td><td>|--|||||[||||||||-||-]PAM</td><td>21704</td><td>Intergenic</td><td>Eya4</td><td>ENSMUSG00000010461</td></tr>
<tr><td>chr9</td><td>9518827</td><td>9518849</td><td>+</td><td>4</td><td>AGGATGAGGCTTGAGGACAT</td><td>GGG</td><td>-||-||||[||||||||||-|-]PAM</td><td>NA</td><td>Intergenic</td><td>NA</td><td>NA</td></tr>
<tr><td>chr13</td><td>6642548</td><td>6642570</td><td>-</td><td>4</td><td>CGATTGAGGCTTGAGGAGGT</td><td>GGG</td><td>||--||||[||||||||||||--]PAM</td><td>5898</td><td>Intronic</td><td>Pfkp</td><td>ENSMUSG00000021196</td></tr>
</table>

Species: Mouse (Mus musculus GRCm38/mm10)
Input: CCTCAGTCCTCACTAGTTTCTGC
PAM: NGG
Target site length: 20
Core length: 12
Core MM: 2
Total MM: 4

<table>
<tr><td>T1</td><td>GCAGAAACTAGTGAGGACTGAGG</td><td>1000</td><td>CRISPRater score</td><td>0.74922171</td><td></td><td></td><td></td><td></td><td></td><td></td><td></td></tr>
<tr><th>Chromosome</th><th>start</th><th>end</th><th>strand</th><th>MM</th><th>target_seq</th><th>PAM</th><th>alignment</th><th>distance</th><th>position</th><th>gene name</th><th>gene id</th></tr>
<tr><td>chr6</td><td>130231539</td><td>130231561</td><td>-</td><td>0</td><td>GCAGAAACTAGTGAGGACTG</td><td>AGG</td><td>||||||||[||||||||||||]PAM</td><td>0</td><td>Exonic</td><td>Klra7</td><td>ENSMUSG00000067599</td></tr>
<tr><td>chr3</td><td>157964562</td><td>157964584</td><td>+</td><td>4</td><td>AGGGAAAGTAGTGAGGACTG</td><td>TGG</td><td>---||||-[||||||||||||]PAM</td><td>2180</td><td>Intronic</td><td>Ankrd13c</td><td>ENSMUSG00000039988</td></tr>
<tr><td>chr2</td><td>134588585</td><td>134588607</td><td>+</td><td>4</td><td>CAAGCACCTAGTGAGGACTG</td><td>GGG</td><td>--||-|-|[||||||||||||]PAM</td><td>5578</td><td>Intergenic</td><td>Tmx4</td><td>ENSMUSG00000034723</td></tr>
<tr><td>chr4</td><td>59752148</td><td>59752170</td><td>+</td><td>4</td><td>ACAGCCCCTAGTGAGGACTG</td><td>GGG</td><td>-|||---|[||||||||||||]PAM</td><td>63</td><td>Intronic</td><td>E130308A1</td><td>ENSMUSG00000045071</td></tr>
<tr><td>chr5</td><td>43599323</td><td>43599345</td><td>+</td><td>4</td><td>AGAGAATGTAGTGAGGACTG</td><td>AGG</td><td>--||||--[||||||||||||]PAM</td><td>100</td><td>Intronic</td><td>Gm15866</td><td>ENSMUSG00000087290</td></tr>
<tr><td>chr9</td><td>117430529</td><td>117430551</td><td>+</td><td>4</td><td>TGAGAGACAAGTGAGGACTG</td><td>CGG</td><td>--|||-||[-|||||||||||]PAM</td><td>26187</td><td>Intronic</td><td>Gm20396</td><td>ENSMUSG00000092504</td></tr>
<tr><td>chr13</td><td>113100459</td><td>113100481</td><td>-</td><td>2</td><td>GCAGAAGCTAATGAGGACTG</td><td>TGG</td><td>||||||-|[||-|||||||||]PAM</td><td>401</td><td>Intronic</td><td>Gzma</td><td>ENSMUSG00000023132</td></tr>
<tr><td>chrX</td><td>165875798</td><td>165875820</td><td>+</td><td>4</td><td>GGTGATACCAGTGAGGACTG</td><td>AGG</td><td>|--||-||[-|||||||||||]PAM</td><td>NA</td><td>Intergenic</td><td>NA</td><td>NA</td></tr>
<tr><td>chr2</td><td>31299624</td><td>31299646</td><td>+</td><td>3</td><td>GCAGTAAGGAGTGAGGACTG</td><td>TGG</td><td>||||-||-[-|||||||||||]PAM</td><td>3635</td><td>Intergenic</td><td>Ncs1</td><td>ENSMUSG00000062661</td></tr>
<tr><td>chr19</td><td>6308245</td><td>6308267</td><td>-</td><td>4</td><td>GGAGTGACCAGTGAGGACTG</td><td>GGG</td><td>|-||--||[-|||||||||||]PAM</td><td>859</td><td>Intronic</td><td>Cdc42bpg</td><td>ENSMUSG00000024769</td></tr>
<tr><td>chr10</td><td>70165730</td><td>70165752</td><td>+</td><td>4</td><td>GTAGGCACCAGTGAGGACTG</td><td>GGG</td><td>|-||--||[-|||||||||||]PAM</td><td>1250</td><td>Intronic</td><td>Ccdc6</td><td>ENSMUSG00000048701</td></tr>
<tr><td>chr12</td><td>107033559</td><td>107033581</td><td>-</td><td>4</td><td>GCTGGAGATAGTGAGGACTG</td><td>AGG</td><td>||-|-|--[||||||||||||]PAM</td><td>45779</td><td>Intergenic</td><td>Gm16086</td><td>ENSMUSG00000085120</td></tr>
<tr><td>chr6</td><td>19453748</td><td>19453770</td><td>+</td><td>4</td><td>CCAAACACTTGTGAGGACTG</td><td>TGG</td><td>-||-|-||[|-||||||||||]PAM</td><td>NA</td><td>Intergenic</td><td>NA</td><td>NA</td></tr>
<tr><td>chr4</td><td>46849623</td><td>46849645</td><td>+</td><td>4</td><td>GGACAAAGCAGTGAGGACTG</td><td>AGG</td><td>|-|-|||-[-|||||||||||]PAM</td><td>3172</td><td>Intronic</td><td>Gabbr2</td><td>ENSMUSG00000039809</td></tr>
<tr><td>chr3</td><td>107018643</td><td>107018665</td><td>-</td><td>3</td><td>GGAGAGACTAGAGAGGACTG</td><td>GGG</td><td>|-|||-||[|||-||||||||]PAM</td><td>17504</td><td>Intronic</td><td>Kcna3</td><td>ENSMUSG00000047959</td></tr>
<tr><td>chr5</td><td>25155755</td><td>25155777</td><td>-</td><td>4</td><td>GCTGAGCCCAGTGAGGACTG</td><td>GGG</td><td>||-||--|[-|||||||||||]PAM</td><td>12595</td><td>Intergenic</td><td>Gm26051</td><td>ENSMUSG00000077832</td></tr>
<tr><td>chr7</td><td>138434591</td><td>138434613</td><td>+</td><td>4</td><td>CTAGAATCTAATGAGGACTG</td><td>AGG</td><td>--||||-|[||-|||||||||]PAM</td><td>36861</td><td>Intergenic</td><td>Tcerg1l</td><td>ENSMUSG00000091002</td></tr>
<tr><td>chr12</td><td>76136502</td><td>76136524</td><td>+</td><td>4</td><td>GTAGAATGAAGTGAGGACTG</td><td>AGG</td><td>|-||||--[-|||||||||||]PAM</td><td>2559</td><td>Intronic</td><td>Esr2</td><td>ENSMUSG00000021055</td></tr>
<tr><td>chr15</td><td>30502279</td><td>30502301</td><td>-</td><td>4</td><td>GCAGTGAATGGTGAGGACTG</td><td>TGG</td><td>||||--|-[|-||||||||||]PAM</td><td>21385</td><td>Intronic</td><td>Ctnnd2</td><td>ENSMUSG00000022240</td></tr>
<tr><td>chr18</td><td>73762143</td><td>73762165</td><td>+</td><td>4</td><td>GCAGAGGTGAGTGAGGACTG</td><td>AGG</td><td>|||||---[-|||||||||||]PAM</td><td>7664</td><td>Intergenic</td><td>Elac1</td><td>ENSMUSG00000036941</td></tr>
<tr><td>chr17</td><td>11869659</td><td>11869681</td><td>-</td><td>4</td><td>GTAGAGAGTATTGAGGACTG</td><td>AGG</td><td>|-|||-|-[||-|||||||||]PAM</td><td>2098</td><td>Intronic</td><td>Park2</td><td>ENSMUSG00000023826</td></tr>
<tr><td>chr17</td><td>5831739</td><td>5831761</td><td>+</td><td>4</td><td>GAAGAAGCAGGTGAGGACTG</td><td>TGG</td><td>|-||||-|[--||||||||||]PAM</td><td>6207</td><td>Intergenic</td><td>Gm26622</td><td>ENSMUSG00000096962</td></tr>
<tr><td>chr8</td><td>43363458</td><td>43363480</td><td>+</td><td>4</td><td>CCAGAGATTAGAGAGGACTG</td><td>AGG</td><td>-||||-|-[|||-||||||||]PAM</td><td>14495</td><td>Intergenic</td><td>Gm22986</td><td>ENSMUSG00000084702</td></tr>
<tr><td>chr6</td><td>71808972</td><td>71808994</td><td>+</td><td>4</td><td>GCAGGGTCTAGGGAGGACTG</td><td>AGG</td><td>||||---|[|||-||||||||]PAM</td><td>0</td><td>Exonic</td><td>Reep1</td><td>ENSMUSG00000052852</td></tr>
<tr><td>chr10</td><td>93958481</td><td>93958503</td><td>+</td><td>4</td><td>GAAGAAAGGACTGAGGACTG</td><td>TGG</td><td>|-|||||-[-|-|||||||||]PAM</td><td>3019</td><td>Intronic</td><td>Vezt</td><td>ENSMUSG00000036099</td></tr>
<tr><td>chr18</td><td>70711859</td><td>70711881</td><td>-</td><td>4</td><td>GAAAAAAATAGTCAGGACTG</td><td>TGG</td><td>|-|-|||-[||||-|||||||]PAM</td><td>85728</td><td>Intergenic</td><td>Mbd2</td><td>ENSMUSG00000024513</td></tr>
<tr><td>chr19</td><td>10058070</td><td>10058092</td><td>-</td><td>4</td><td>CCAGAAAATGCTGAGGACTG</td><td>AGG</td><td>-||||||-[|--|||||||||]PAM</td><td>0</td><td>Exonic</td><td>Fads3</td><td>ENSMUSG00000024664</td></tr>
<tr><td>chr15</td><td>30010386</td><td>30010408</td><td>-</td><td>4</td><td>GCAGCACGTAGGGAGGACTG</td><td>AGG</td><td>||||-|--[|||-||||||||]PAM</td><td>6123</td><td>Intergenic</td><td>Gm23327</td><td>ENSMUSG00000088045</td></tr>
<tr><td>chr5</td><td>46281295</td><td>46281317</td><td>+</td><td>4</td><td>GGAGAAATTTTTGAGGACTG</td><td>GGG</td><td>|-|||||-[|--|||||||||]PAM</td><td>NA</td><td>Intergenic</td><td>NA</td><td>NA</td></tr>
<tr><td>chr13</td><td>36099374</td><td>36099396</td><td>-</td><td>4</td><td>GAGGAAACTACAGAGGACTG</td><td>TGG</td><td>|--|||||[||--||||||||]PAM</td><td>6448</td><td>Intronic</td><td>Lyrm4</td><td>ENSMUSG00000046573</td></tr>
<tr><td>chr4</td><td>39397571</td><td>39397593</td><td>+</td><td>4</td><td>CCAGTAACAAGTAAGGACTG</td><td>AGG</td><td>-|||-|||[-|||-|||||||]PAM</td><td>452</td><td>Intergenic</td><td>Gm17094</td><td>ENSMUSG00000090839</td></tr>
<tr><td>chr18</td><td>34673142</td><td>34673164</td><td>+</td><td>4</td><td>GCAGAGGGTAGGGAGGACTG</td><td>AGG</td><td>|||||---[|||-||||||||]PAM</td><td>16739</td><td>Intergenic</td><td>Gfra3</td><td>ENSMUSG00000024366</td></tr>
<tr><td>chrX</td><td>81512471</td><td>81512493</td><td>+</td><td>4</td><td>GAAAAAACTAAGGAGGACTG</td><td>TGG</td><td>|-|-||||[||--||||||||]PAM</td><td>54350</td><td>Intergenic</td><td>4930595M</td><td>ENSMUSG00000060673</td></tr>
<tr><td>chr8</td><td>101658323</td><td>101658345</td><td>+</td><td>3</td><td>GCAAGAACTAGTGAAGACTG</td><td>AGG</td><td>|||--|||[||||||-|||||]PAM</td><td>NA</td><td>Intergenic</td><td>NA</td><td>NA</td></tr>
<tr><td>chr16</td><td>32285130</td><td>32285152</td><td>+</td><td>4</td><td>GTGGAAACTGGTAAGGACTG</td><td>AGG</td><td>|--|||||[|-||-|||||||]PAM</td><td>218</td><td>Intronic</td><td>Rnf168</td><td>ENSMUSG00000014074</td></tr>
<tr><td>chr3</td><td>122432637</td><td>122432659</td><td>+</td><td>4</td><td>GCGTAAACTATGGAGGACTG</td><td>AGG</td><td>||--||||[||--||||||||]PAM</td><td>5489</td><td>Intronic</td><td>Bcar3</td><td>ENSMUSG00000028121</td></tr>
<tr><td>chr6</td><td>119433610</td><td>119433632</td><td>+</td><td>4</td><td>GCAGGCAGTAGTCAGGACTG</td><td>GGG</td><td>||||--|-[||||-|||||||]PAM</td><td>0</td><td>Exonic</td><td>Wnt5b</td><td>ENSMUSG00000030170</td></tr>
<tr><td>chr7</td><td>74854728</td><td>74854750</td><td>+</td><td>4</td><td>GCAGGCACTTGGGAGGACTG</td><td>AGG</td><td>||||--||[|-|-||||||||]PAM</td><td>915</td><td>Intergenic</td><td>RP23-53N2</td><td>ENSMUSG00000108652</td></tr>
<tr><td>chr8</td><td>25597339</td><td>25597361</td><td>+</td><td>4</td><td>GCGGAAAGTGGGGAGGACTG</td><td>CGG</td><td>||-||||-[|-|-||||||||]PAM</td><td>0</td><td>Exonic</td><td>Letm2</td><td>ENSMUSG00000037363</td></tr>
<tr><td>chr9</td><td>22105927</td><td>22105949</td><td>-</td><td>4</td><td>TCAGGAACTACTTAGGACTG</td><td>TGG</td><td>-|||-|||[||-|-|||||||]PAM</td><td>147</td><td>Intronic</td><td>Cnn1</td><td>ENSMUSG00000001349</td></tr>
<tr><td>chr10</td><td>29747260</td><td>29747282</td><td>+</td><td>4</td><td>GTAAAGACTAGTGATGACTG</td><td>GGG</td><td>|-|-|-||[||||||-|||||]PAM</td><td>47880</td><td>Intergenic</td><td>Gm10275</td><td>ENSMUSG00000069682</td></tr>
<tr><td>chr14</td><td>30292534</td><td>30292556</td><td>+</td><td>4</td><td>GCAGAATGGAGAGAGGACTG</td><td>TGG</td><td>||||||--[-||-||||||||]PAM</td><td>54242</td><td>Intronic</td><td>Cacna1d</td><td>ENSMUSG00000015968</td></tr>
<tr><td>chr1</td><td>46398457</td><td>46398479</td><td>-</td><td>4</td><td>ACAGAAAGTGGTAAGGACTG</td><td>AGG</td><td>-||||||-[|-||-|||||||]PAM</td><td>16023</td><td>Intergenic</td><td>Gm8326</td><td>ENSMUSG00000101262</td></tr>
<tr><td>chr17</td><td>49967592</td><td>49967614</td><td>+</td><td>3</td><td>GCAGAAAGTATTCAGGACTG</td><td>GGG</td><td>|||||||-[||-|-|||||||]PAM</td><td>24643</td><td>Intergenic</td><td>Rftn1</td><td>ENSMUSG00000039316</td></tr>
<tr><td>chr19</td><td>5531258</td><td>5531280</td><td>+</td><td>4</td><td>CCAGAAGCCAGTGTGGACTG</td><td>TGG</td><td>-|||||-|[-||||-||||||]PAM</td><td>17857</td><td>Intergenic</td><td>Ovol1</td><td>ENSMUSG00000024922</td></tr>
<tr><td>chr11</td><td>3734138</td><td>3734160</td><td>+</td><td>4</td><td>GCAGCATGTAGTGTGGACTG</td><td>GGG</td><td>||||-|--[|||||-||||||]PAM</td><td>7563</td><td>Intronic</td><td>Osbp2</td><td>ENSMUSG00000020435</td></tr>
<tr><td>chr15</td><td>64751194</td><td>64751216</td><td>-</td><td>2</td><td>GCAGAAACAAGTGAGTACTG</td><td>AGG</td><td>||||||||[-||||||-||||]PAM</td><td>3568</td><td>Intronic</td><td>Adcy8</td><td>ENSMUSG00000022376</td></tr>
<tr><td>chr11</td><td>82383337</td><td>82383359</td><td>+</td><td>4</td><td>ACAGAGATTAGTGATGACTG</td><td>TGG</td><td>-||||-|-[||||||-|||||]PAM</td><td>5541</td><td>Intergenic</td><td>Tmem132e</td><td>ENSMUSG00000020701</td></tr>
<tr><td>chr10</td><td>79591834</td><td>79591856</td><td>-</td><td>4</td><td>GCCGAAAACAGTGTGGACTG</td><td>CGG</td><td>||-||||-[-||||-||||||]PAM</td><td>4503</td><td>Intergenic</td><td>Theg</td><td>ENSMUSG00000020317</td></tr>
<tr><td>chr6</td><td>130338238</td><td>130338260</td><td>-</td><td>3</td><td>ACAGAAACGAGTGAGAACTG</td><td>AGG</td><td>-|||||||[-||||||-||||]PAM</td><td>664</td><td>Intergenic</td><td>Klra3</td><td>ENSMUSG00000067591</td></tr>
<tr><td>chr16</td><td>65011768</td><td>65011790</td><td>-</td><td>3</td><td>TCAGAAACAAGTGAGAACTG</td><td>TGG</td><td>-|||||||[-||||||-||||]PAM</td><td>62332</td><td>Intronic</td><td>Htr1f</td><td>ENSMUSG00000050783</td></tr>
<tr><td>chr6</td><td>57610310</td><td>57610332</td><td>-</td><td>4</td><td>GAAGAAATTAGCTAGGACTG</td><td>AGG</td><td>|-|||||-[|||--|||||||]PAM</td><td>0</td><td>Exonic</td><td>Herc6</td><td>ENSMUSG00000029798</td></tr>
<tr><td>chr7</td><td>37429583</td><td>37429605</td><td>+</td><td>4</td><td>GCAGCAGCCAGTGAAGACTG</td><td>AGG</td><td>||||-|-|[-|||||-|||||]PAM</td><td>35515</td><td>Intergenic</td><td>RP23-291B</td><td>ENSMUSG00000108790</td></tr>
<tr><td>chr14</td><td>64553908</td><td>64553930</td><td>-</td><td>4</td><td>CCAGTAACTAGTAGGGACTG</td><td>AGG</td><td>-|||-|||[||||--||||||]PAM</td><td>33401</td><td>Intergenic</td><td>Mir124a-1h</td><td>ENSMUSG00000097545</td></tr>
<tr><td>chr6</td><td>130407860</td><td>130407882</td><td>-</td><td>3</td><td>GCAGAACCAAGTGAGTACTG</td><td>AGG</td><td>||||||-|[-||||||-||||]PAM</td><td>0</td><td>Exonic</td><td>Gm6584</td><td>ENSMUSG00000084204</td></tr>
<tr><td>chr4</td><td>82242632</td><td>82242654</td><td>-</td><td>4</td><td>ATAGAAACTGGTGAGAACTG</td><td>GGG</td><td>--||||||[|-|||||-||||]PAM</td><td>47519</td><td>Intergenic</td><td>Nfib</td><td>ENSMUSG00000008575</td></tr>
<tr><td>chr1</td><td>135533097</td><td>135533119</td><td>+</td><td>4</td><td>GAATGAACTAGTGAGGGCTG</td><td>GGG</td><td>|-|--|||[||||||||-|||]PAM</td><td>383</td><td>Intronic</td><td>Nav1</td><td>ENSMUSG00000009418</td></tr>
<tr><td>chr9</td><td>31553558</td><td>31553580</td><td>+</td><td>4</td><td>ACAGAAGCTAGTCTGGACTG</td><td>TGG</td><td>-|||||-|[||||--||||||]PAM</td><td>89320</td><td>Intergenic</td><td>Tmem45b</td><td>ENSMUSG00000041737</td></tr>
<tr><td>chr9</td><td>96677280</td><td>96677302</td><td>-</td><td>4</td><td>TGAGAATCTAGTGAGGTCTG</td><td>CGG</td><td>--||||-|[||||||||-|||]PAM</td><td>5468</td><td>Intergenic</td><td>Zbtb38</td><td>ENSMUSG00000040433</td></tr>
<tr><td>chr6</td><td>129913902</td><td>129913924</td><td>-</td><td>4</td><td>ACAGAAAAGAGTGAGAACTG</td><td>AGG</td><td>-||||||-[-||||||-||||]PAM</td><td>678</td><td>Intronic</td><td>Klra5</td><td>ENSMUSG00000030173</td></tr>
<tr><td>chr6</td><td>130027677</td><td>130027699</td><td>-</td><td>4</td><td>ACAGAAAAGAGTGAGAACTG</td><td>AGG</td><td>-||||||-[-||||||-||||]PAM</td><td>723</td><td>Intergenic</td><td>Klra6</td><td>ENSMUSG00000061769</td></tr>
<tr><td>chr8</td><td>48267818</td><td>48267840</td><td>-</td><td>4</td><td>CCAGAAATCAGTGAGAACTG</td><td>TGG</td><td>-||||||-[-||||||-||||]PAM</td><td>8075</td><td>Intronic</td><td>Tenm3</td><td>ENSMUSG00000031561</td></tr>
<tr><td>chr16</td><td>44536874</td><td>44536896</td><td>-</td><td>4</td><td>GCGGAGACTTGTGAGAACTG</td><td>GGG</td><td>||-||-||[|-|||||-||||]PAM</td><td>15539</td><td>Intronic</td><td>Boc</td><td>ENSMUSG00000022687</td></tr>
<tr><td>chr11</td><td>73480672</td><td>73480694</td><td>-</td><td>4</td><td>ACAGAAGCTGGTGAGCACTG</td><td>TGG</td><td>-|||||-|[|-|||||-||||]PAM</td><td>3080</td><td>Intergenic</td><td>RP23-103M</td><td>ENSMUSG00000108742</td></tr>
<tr><td>chr1</td><td>112744289</td><td>112744311</td><td>+</td><td>4</td><td>ACAGAAGCTGGTGAGTACTG</td><td>TGG</td><td>-|||||-|[|-|||||-||||]PAM</td><td>NA</td><td>Intergenic</td><td>NA</td><td>NA</td></tr>
<tr><td>chr2</td><td>125086160</td><td>125086182</td><td>+</td><td>4</td><td>GTAGAAAGTAGAGATGACTG</td><td>GGG</td><td>|-|||||-[|||-||-|||||]PAM</td><td>0</td><td>Exonic</td><td>Myef2</td><td>ENSMUSG00000027201</td></tr>
<tr><td>chr18</td><td>61109581</td><td>61109603</td><td>-</td><td>3</td><td>GCAGACACAAGTGAGGCCTG</td><td>GGG</td><td>|||||-||[-|||||||-|||]PAM</td><td>13</td><td>Intronic</td><td>Csf1r</td><td>ENSMUSG00000024621</td></tr>
<tr><td>chr15</td><td>24794874</td><td>24794896</td><td>-</td><td>4</td><td>GAAGAAATTTGTGAGCACTG</td><td>TGG</td><td>|-|||||-[|-|||||-||||]PAM</td><td>25649</td><td>Intergenic</td><td>Gm22810</td><td>ENSMUSG00000064675</td></tr>
<tr><td>chr6</td><td>129874829</td><td>129874851</td><td>-</td><td>3</td><td>GCAGAACCAAGTGAGGCCTG</td><td>AGG</td><td>||||||-|[-|||||||-|||]PAM</td><td>0</td><td>Exonic</td><td>Klra17</td><td>ENSMUSG00000014543</td></tr>
<tr><td>chr6</td><td>131245248</td><td>131245270</td><td>-</td><td>3</td><td>GCAGAACCCAGTGAGGCCTG</td><td>AGG</td><td>||||||-|[-|||||||-|||]PAM</td><td>0</td><td>Exonic</td><td>Klra2</td><td>ENSMUSG00000030187</td></tr>
<tr><td>chr11</td><td>103104166</td><td>103104188</td><td>+</td><td>4</td><td>ACAGAAGGTAGTGAGGGCTG</td><td>TGG</td><td>-|||||--[||||||||-|||]PAM</td><td>0</td><td>Exonic</td><td>Acbd4</td><td>ENSMUSG00000056938</td></tr>
<tr><td>chr7</td><td>11977186</td><td>11977208</td><td>+</td><td>4</td><td>TCAGAAAGTACTGAGTACTG</td><td>TGG</td><td>-||||||-[||-||||-||||]PAM</td><td>0</td><td>Exonic</td><td>Vmn1r-ps5</td><td>ENSMUSG00000092208</td></tr>
<tr><td>chr4</td><td>3172739</td><td>3172761</td><td>-</td><td>4</td><td>TCAGAAAGTATTGAGCACTG</td><td>TGG</td><td>-||||||-[||-||||-||||]PAM</td><td>0</td><td>Exonic</td><td>Vmn1r2</td><td>ENSMUSG00000094798</td></tr>
<tr><td>chr5</td><td>134523684</td><td>134523706</td><td>-</td><td>4</td><td>TCAGACACCAGTGAGGGCTG</td><td>GGG</td><td>-||||-||[-|||||||-|||]PAM</td><td>542</td><td>Intronic</td><td>Clip2</td><td>ENSMUSG00000063146</td></tr>
<tr><td>chr19</td><td>46390237</td><td>46390259</td><td>-</td><td>4</td><td>GCAGGTAATAGTGAGGGCTG</td><td>GGG</td><td>||||--|-[||||||||-|||]PAM</td><td>2721</td><td>Intronic</td><td>Actr1a</td><td>ENSMUSG00000025228</td></tr>
<tr><td>chr13</td><td>95948548</td><td>95948570</td><td>-</td><td>4</td><td>TCAGAACCTAGAGAGCACTG</td><td>GGG</td><td>-|||||-|[|||-|||-||||]PAM</td><td>6024</td><td>Intergenic</td><td>Sv2c</td><td>ENSMUSG00000051111</td></tr>
<tr><td>chr18</td><td>75967447</td><td>75967469</td><td>-</td><td>4</td><td>GAAGAAAGTAGAGAGTACTG</td><td>TGG</td><td>|-|||||-[|||-|||-||||]PAM</td><td>44160</td><td>Intronic</td><td>Zbtb7c</td><td>ENSMUSG00000044646</td></tr>
<tr><td>chr12</td><td>97741088</td><td>97741110</td><td>-</td><td>4</td><td>GAAGAAAGTAGAGAGCACTG</td><td>AGG</td><td>|-|||||-[|||-|||-||||]PAM</td><td>NA</td><td>Intergenic</td><td>NA</td><td>NA</td></tr>
<tr><td>chr18</td><td>21899035</td><td>21899057</td><td>+</td><td>4</td><td>GCAAATACTTGTGAGGTCTG</td><td>AGG</td><td>|||-|-||[|-||||||-|||]PAM</td><td>15890</td><td>Intronic</td><td>Ccdc178</td><td>ENSMUSG00000024306</td></tr>
<tr><td>chr11</td><td>53249890</td><td>53249912</td><td>-</td><td>4</td><td>GCAGGATCTAGGGAGTACTG</td><td>TGG</td><td>||||-|-|[|||-|||-||||]PAM</td><td>9902</td><td>Intergenic</td><td>Hspa4</td><td>ENSMUSG00000020361</td></tr>
<tr><td>chr10</td><td>90983105</td><td>90983127</td><td>+</td><td>4</td><td>ACAGATGCTAGTGAGGATTG</td><td>GGG</td><td>-||||--|[|||||||||-||]PAM</td><td>6184</td><td>Intergenic</td><td>Apaf1</td><td>ENSMUSG00000019979</td></tr>
<tr><td>chr19</td><td>12359999</td><td>12360021</td><td>+</td><td>4</td><td>GCTGAAGCTAGTTAGAACTG</td><td>GGG</td><td>||-|||-|[||||-||-||||]PAM</td><td>34244</td><td>Intergenic</td><td>Olfr1440</td><td>ENSMUSG00000046650</td></tr>
<tr><td>chr11</td><td>75138590</td><td>75138612</td><td>+</td><td>4</td><td>GGAGCAACTAGTGTGAACTG</td><td>TGG</td><td>|-||-|||[|||||-|-||||]PAM</td><td>581</td><td>Intronic</td><td>Smg6</td><td>ENSMUSG00000038290</td></tr>
<tr><td>chr8</td><td>92215012</td><td>92215034</td><td>+</td><td>4</td><td>TCAGACACTAGTGGGCACTG</td><td>GGG</td><td>-||||-||[|||||-|-||||]PAM</td><td>NA</td><td>Intergenic</td><td>NA</td><td>NA</td></tr>
<tr><td>chr6</td><td>125192778</td><td>125192800</td><td>+</td><td>4</td><td>GCGGAGAATAGTGAGGAGTG</td><td>CGG</td><td>||-||-|-[|||||||||-||]PAM</td><td>0</td><td>Exonic</td><td>Mrpl51</td><td>ENSMUSG00000030335</td></tr>
<tr><td>chr8</td><td>72403824</td><td>72403846</td><td>-</td><td>4</td><td>GCAGTGAATAGTGAGGAGTG</td><td>TGG</td><td>||||--|-[|||||||||-||]PAM</td><td>3591</td><td>Intronic</td><td>Eps15l1</td><td>ENSMUSG00000006276</td></tr>
<tr><td>chrX</td><td>168093533</td><td>168093555</td><td>-</td><td>4</td><td>ATAGAAACTAGTGAATACTG</td><td>AGG</td><td>--||||||[||||||--||||]PAM</td><td>18611</td><td>Intronic</td><td>Gm15237</td><td>ENSMUSG00000084180</td></tr>
<tr><td>chr4</td><td>136196303</td><td>136196325</td><td>-</td><td>4</td><td>GCACAGACCAGTGAGGAGTG</td><td>GGG</td><td>|||-|-||[-||||||||-||]PAM</td><td>246</td><td>Intergenic</td><td>E2f2</td><td>ENSMUSG00000018983</td></tr>
<tr><td>chr15</td><td>77113424</td><td>77113446</td><td>-</td><td>3</td><td>GGAGAAACTAGAGAGGAGTG</td><td>TGG</td><td>|-||||||[|||-|||||-||]PAM</td><td>159</td><td>Intronic</td><td>Rbfox2</td><td>ENSMUSG00000033565</td></tr>
<tr><td>chr18</td><td>12863878</td><td>12863900</td><td>+</td><td>4</td><td>GCAGAATTTAGTCAGAACTG</td><td>GGG</td><td>||||||--[||||-||-||||]PAM</td><td>498</td><td>Intronic</td><td>Osbpl1a</td><td>ENSMUSG00000044252</td></tr>
<tr><td>chr13</td><td>55731505</td><td>55731527</td><td>+</td><td>4</td><td>CCAGAAATGAGTGAGGAATG</td><td>AGG</td><td>-||||||-[-||||||||-||]PAM</td><td>1409</td><td>Intronic</td><td>Pcbd2</td><td>ENSMUSG00000021496</td></tr>
<tr><td>chr6</td><td>64038155</td><td>64038177</td><td>+</td><td>4</td><td>GGAGAAAGAAGTGAGGAGTG</td><td>TGG</td><td>|-|||||-[-||||||||-||]PAM</td><td>2252</td><td>Intronic</td><td>RP23-142H</td><td>ENSMUSG00000107864</td></tr>
<tr><td>chr15</td><td>13825084</td><td>13825106</td><td>+</td><td>4</td><td>ACAGGAACTAGTGATCACTG</td><td>GGG</td><td>-|||-|||[||||||--||||]PAM</td><td>NA</td><td>Intergenic</td><td>NA</td><td>NA</td></tr>
<tr><td>chr7</td><td>141273454</td><td>141273476</td><td>-</td><td>4</td><td>GCAAAAATCAGTGAGGAGTG</td><td>TGG</td><td>|||-|||-[-||||||||-||]PAM</td><td>212</td><td>Intronic</td><td>Cdhr5</td><td>ENSMUSG00000025497</td></tr>
<tr><td>chr11</td><td>71608435</td><td>71608457</td><td>-</td><td>3</td><td>GCAGAAATTAGTGATCACTG</td><td>AGG</td><td>|||||||-[||||||--||||]PAM</td><td>35252</td><td>Intergenic</td><td>9230020A</td><td>ENSMUSG00000072753</td></tr>
<tr><td>chr11</td><td>32771248</td><td>32771270</td><td>-</td><td>4</td><td>TCAGATACTACTGAGGAATG</td><td>CGG</td><td>-||||-||[||-||||||-||]PAM</td><td>3412</td><td>Intergenic</td><td>Gm8192</td><td>ENSMUSG00000098054</td></tr>
<tr><td>chr7</td><td>114214016</td><td>114214038</td><td>-</td><td>4</td><td>CCAGAAGCTAGTGACAACTG</td><td>GGG</td><td>-|||||-|[||||||--||||]PAM</td><td>1521</td><td>Intergenic</td><td>Copb1</td><td>ENSMUSG00000030754</td></tr>
</table>

<table>
<tr><td>chr5</td><td>149976887</td><td>149976909</td><td>+</td><td>4</td><td>TCAGAAGCTAATGAGGAGTG</td><td>GGG</td><td>-|||||-|[||-||||||-||]PAM</td><td>24896</td><td>Intergenic</td><td>Gm21048</td><td>ENSMUSG00000107056</td></tr>
<tr><td>chrX</td><td>98514782</td><td>98514804</td><td>+</td><td>4</td><td>GCAGAATGTAGGGAGGGCTG</td><td>GGG</td><td>||||||--[|||-||||-|||]PAM</td><td>395</td><td>Intergenic</td><td>Gm14815</td><td>ENSMUSG00000081397</td></tr>
<tr><td>chr5</td><td>67604957</td><td>67604979</td><td>+</td><td>4</td><td>GCAGATGCTGGTGAGGATTG</td><td>TGG</td><td>|||||--|[|-|||||||-||]PAM</td><td>2239</td><td>Intergenic</td><td>1700025A0</td><td>ENSMUSG00000106947</td></tr>
<tr><td>chr16</td><td>86248322</td><td>86248344</td><td>+</td><td>4</td><td>GCAGATGCTGGTGAGGAATG</td><td>TGG</td><td>|||||--|[|-|||||||-||]PAM</td><td>NA</td><td>Intergenic</td><td>NA</td><td>NA</td></tr>
<tr><td>chr4</td><td>137949833</td><td>137949855</td><td>-</td><td>3</td><td>GCAGGAACGAGTGAGGACGG</td><td>GGG</td><td>||||-|||[-|||||||||-|]PAM</td><td>288</td><td>Intronic</td><td>Ece1</td><td>ENSMUSG00000057530</td></tr>
<tr><td>chr12</td><td>117345920</td><td>117345942</td><td>-</td><td>4</td><td>GATGAAACAAGTGAGGACCG</td><td>AGG</td><td>|--|||||[-|||||||||-|]PAM</td><td>8898</td><td>Intergenic</td><td>Gm5441</td><td>ENSMUSG00000101930</td></tr>
<tr><td>chr4</td><td>139618169</td><td>139618191</td><td>-</td><td>4</td><td>GAAGAGAATAGTGAGGACAG</td><td>GGG</td><td>|-|||-|-[||||||||||-|]PAM</td><td>0</td><td>Exonic</td><td>Iffo2</td><td>ENSMUSG00000041025</td></tr>
<tr><td>chr6</td><td>121342781</td><td>121342803</td><td>+</td><td>4</td><td>GAAGACACTAGTCAGGAGTG</td><td>AGG</td><td>|-|||-||[||||-||||-||]PAM</td><td>273</td><td>Intergenic</td><td>Slc6a12</td><td>ENSMUSG00000030109</td></tr>
<tr><td>chr7</td><td>44880554</td><td>44880576</td><td>-</td><td>2</td><td>GCAGAATCTAGTGAGGACTC</td><td>GGG</td><td>||||||-|[|||||||||||-]PAM</td><td>0</td><td>Exonic</td><td>Med25</td><td>ENSMUSG00000002968</td></tr>
<tr><td>chr16</td><td>33087830</td><td>33087852</td><td>+</td><td>4</td><td>GAAGACACTAGTGAAGGCTG</td><td>GGG</td><td>|-|||-||[||||||-|-|||]PAM</td><td>250</td><td>Intronic</td><td>Lmln</td><td>ENSMUSG00000022802</td></tr>
<tr><td>chr7</td><td>103080072</td><td>103080094</td><td>-</td><td>4</td><td>GCAGTAAGTAGAGAGGAATG</td><td>GGG</td><td>||||-||-[|||-|||||-||]PAM</td><td>5426</td><td>Intergenic</td><td>Olfr584</td><td>ENSMUSG00000073959</td></tr>
<tr><td>chr11</td><td>74143414</td><td>74143436</td><td>+</td><td>4</td><td>GCAGAGATTAGGGAGGATTG</td><td>GGG</td><td>|||||-|-[|||-|||||-||]PAM</td><td>11679</td><td>Intergenic</td><td>Olfr402</td><td>ENSMUSG00000070379</td></tr>
<tr><td>chr15</td><td>10986141</td><td>10986163</td><td>+</td><td>4</td><td>GAAGAAAGAAGTGAGGACAG</td><td>AGG</td><td>|-|||||-[-|||||||||-|]PAM</td><td>1295</td><td>Intronic</td><td>Amacr</td><td>ENSMUSG00000022244</td></tr>
<tr><td>chr19</td><td>11796633</td><td>11796655</td><td>+</td><td>4</td><td>GTAGAAAGCAGTGAGGACAG</td><td>AGG</td><td>|-|||||-[-|||||||||-|]PAM</td><td>4788</td><td>Intronic</td><td>Stx3</td><td>ENSMUSG00000041488</td></tr>
<tr><td>chr13</td><td>51632424</td><td>51632446</td><td>+</td><td>4</td><td>CTAGAAACTAGTGAGAGCTG</td><td>TGG</td><td>--||||||[|||||||--|||]PAM</td><td>12786</td><td>Intergenic</td><td>Cks2</td><td>ENSMUSG00000062248</td></tr>
<tr><td>chr5</td><td>136700086</td><td>136700108</td><td>+</td><td>3</td><td>ACAGAAAGTAGTGAGGACTC</td><td>TGG</td><td>-||||||-[|||||||||||-]PAM</td><td>594</td><td>Intronic</td><td>Myl10</td><td>ENSMUSG00000005474</td></tr>
<tr><td>chr5</td><td>110754489</td><td>110754511</td><td>+</td><td>4</td><td>ACAGATGCTAGTGAGGACTC</td><td>TGG</td><td>-||||--|[|||||||||||-]PAM</td><td>575</td><td>Intronic</td><td>Ep400</td><td>ENSMUSG00000029505</td></tr>
<tr><td>chr13</td><td>34280022</td><td>34280044</td><td>+</td><td>4</td><td>GCTGGATCTAGTGAGGACTT</td><td>GGG</td><td>||-|-|-|[|||||||||||-]PAM</td><td>2770</td><td>Intronic</td><td>Gm24530</td><td>ENSMUSG00000088573</td></tr>
<tr><td>chr1</td><td>134515925</td><td>134515947</td><td>-</td><td>3</td><td>GCAGAATCAAGTGAGGACTA</td><td>AGG</td><td>||||||-|[-||||||||||-]PAM</td><td>2134</td><td>Intergenic</td><td>Gm15454</td><td>ENSMUSG00000096712</td></tr>
<tr><td>chr15</td><td>6569267</td><td>6569289</td><td>-</td><td>4</td><td>GAAGTAACTTGTGAGGACTT</td><td>TGG</td><td>|-||-|||[|-|||||||||-]PAM</td><td>10582</td><td>Intergenic</td><td>Fyb</td><td>ENSMUSG00000022148</td></tr>
<tr><td>chr5</td><td>8894619</td><td>8894641</td><td>-</td><td>4</td><td>GCAGAAGTTAGTGAAGAATG</td><td>AGG</td><td>||||||--[||||||-||-||]PAM</td><td>455</td><td>Intronic</td><td>Abcb4</td><td>ENSMUSG00000042476</td></tr>
<tr><td>chr3</td><td>113816163</td><td>113816185</td><td>+</td><td>4</td><td>AAAGAAACTAGTTAGGACTT</td><td>GGG</td><td>--||||||[||||-||||||-]PAM</td><td>38551</td><td>Intergenic</td><td>Gm18163</td><td>ENSMUSG00000106084</td></tr>
<tr><td>chr5</td><td>116757575</td><td>116757597</td><td>-</td><td>4</td><td>GCAGCAGCTACTGAGGACTC</td><td>CGG</td><td>||||-|-|[||-||||||||-]PAM</td><td>12794</td><td>Intronic</td><td>Gm43122</td><td>ENSMUSG00000106909</td></tr>
<tr><td>chr13</td><td>56100021</td><td>56100043</td><td>+</td><td>4</td><td>GCAGTGACTAGTGAAGACCG</td><td>TGG</td><td>||||--||[|||||||-|||-|]PAM</td><td>846</td><td>Intronic</td><td>H2afy</td><td>ENSMUSG00000015937</td></tr>
<tr><td>chr14</td><td>49265662</td><td>49265684</td><td>-</td><td>4</td><td>GCTGCAACTAGTGAGGGGTG</td><td>GGG</td><td>||-|-|||[|||||||||--||]PAM</td><td>20234</td><td>Intergenic</td><td>1700011H1</td><td>ENSMUSG00000021850</td></tr>
<tr><td>chr17</td><td>30828150</td><td>30828172</td><td>-</td><td>4</td><td>GCAAAAGCTAGTGAGGGTTG</td><td>GGG</td><td>|||-||-|[|||||||||--||]PAM</td><td>2601</td><td>Intronic</td><td>Dnah8</td><td>ENSMUSG00000033826</td></tr>
<tr><td>chr17</td><td>76327738</td><td>76327760</td><td>+</td><td>4</td><td>GGAGAAATTAGTGAGCACAG</td><td>TGG</td><td>|-|||||-[||||||||-||-|]PAM</td><td>42877</td><td>Intergenic</td><td>Gm16391</td><td>ENSMUSG00000066958</td></tr>
<tr><td>chr3</td><td>88073958</td><td>88073980</td><td>-</td><td>4</td><td>GCAGAAGGTAGTGAGCACAG</td><td>TGG</td><td>||||||--[||||||||-||-|]PAM</td><td>471</td><td>Intronic</td><td>Ttc24</td><td>ENSMUSG00000051036</td></tr>
<tr><td>chr3</td><td>86546728</td><td>86546750</td><td>+</td><td>4</td><td>GCAAAAAGTAGTGAGGGCAG</td><td>CGG</td><td>|||-|||-[|||||||||-|-|]PAM</td><td>0</td><td>Exonic</td><td>Mab21l2</td><td>ENSMUSG00000057777</td></tr>
<tr><td>chr8</td><td>20879654</td><td>20879676</td><td>+</td><td>4</td><td>ACAGACACTAGTGAGGAAGG</td><td>AGG</td><td>-||||-||[||||||||||--|]PAM</td><td>2704</td><td>Intergenic</td><td>Gm20796</td><td>ENSMUSG00000095803</td></tr>
</table>

Species: Mouse (Mus musculus GRCm38/mm10)
Input: CCTTTAGTCTCCTCAGGTCTCAC
PAM: NGG
Target site length: 20
Core length: 12
Core MM: 2
Total MM: 4

T1 GTGAGACCTGAGGAGACTAAAGG 1000 CRISPRater score 0.72113809

<table>
<tr><th>Chromosome</th><th>start</th><th>end</th><th>strand</th><th>MM</th><th>target_seq</th><th>PAM</th><th>alignment</th><th>distance</th><th>position</th><th>gene name</th><th>gene id</th></tr>
<tr><td>chr6</td><td>130380574</td><td>130380596</td><td>-</td><td>0</td><td>GTGAGACCTGAGGAGACTAA</td><td>AGG</td><td>||||||||[||||||||||||]PAM</td><td>0</td><td>Exonic</td><td>Klra1</td><td>ENSMUSG00000079853</td></tr>
<tr><td>chr13</td><td>59436908</td><td>59436930</td><td>+</td><td>4</td><td>GCAGGGCCTGAGGAGACTAA</td><td>GGG</td><td>|---|-||[||||||||||||]PAM</td><td>12607</td><td>Intergenic</td><td>Agtpbp1</td><td>ENSMUSG00000021557</td></tr>
<tr><td>chr19</td><td>51602618</td><td>51602640</td><td>+</td><td>3</td><td>CTGAGCCTTGAGGAGACTAA</td><td>GGG</td><td>-||||-|-[||||||||||||]PAM</td><td>NA</td><td>Intergenic</td><td>NA</td><td>NA</td></tr>
<tr><td>chr10</td><td>64573994</td><td>64574016</td><td>-</td><td>3</td><td>GGGAGAGATGAGGAGACTAA</td><td>AGG</td><td>|-||||--[||||||||||||]PAM</td><td>11944</td><td>Intronic</td><td>Ctnna3</td><td>ENSMUSG00000060843</td></tr>
<tr><td>chr7</td><td>139224833</td><td>139224855</td><td>+</td><td>4</td><td>TCGAGGCCGGAGGAGACTAA</td><td>CGG</td><td>--|||-||[-|||||||||||]PAM</td><td>1092</td><td>Intronic</td><td>Lrrc27</td><td>ENSMUSG00000015980</td></tr>
<tr><td>chr1</td><td>159875062</td><td>159875084</td><td>+</td><td>4</td><td>GCTATACCAGAGGAGACTAA</td><td>GGG</td><td>|--|-|||[-|||||||||||]PAM</td><td>769</td><td>Intronic</td><td>Tnr</td><td>ENSMUSG00000015829</td></tr>
<tr><td>chr18</td><td>34497401</td><td>34497423</td><td>+</td><td>4</td><td>GAAAAACCTAAGGAGACTAA</td><td>TGG</td><td>|--|-|||[|-||||||||||]PAM</td><td>0</td><td>Exonic</td><td>Fam13b</td><td>ENSMUSG00000036501</td></tr>
<tr><td>chr18</td><td>61370228</td><td>61370250</td><td>+</td><td>4</td><td>GTTACAGATGAGGAGACTAA</td><td>AGG</td><td>||-|-|--[||||||||||||]PAM</td><td>12195</td><td>Intronic</td><td>Ppargc1b</td><td>ENSMUSG00000033871</td></tr>
<tr><td>chr1</td><td>100956794</td><td>100956816</td><td>-</td><td>4</td><td>CTGGGACTTAAGGAGACTAA</td><td>AGG</td><td>-||-|||-[|-||||||||||]PAM</td><td>94893</td><td>Intergenic</td><td>Gm25537</td><td>ENSMUSG00000077039</td></tr>
<tr><td>chr8</td><td>6835519</td><td>6835541</td><td>-</td><td>4</td><td>TTGAGTACTTAGGAGACTAA</td><td>TGG</td><td>-||||--|[|-||||||||||]PAM</td><td>NA</td><td>Intergenic</td><td>NA</td><td>NA</td></tr>
<tr><td>chr8</td><td>23176436</td><td>23176458</td><td>-</td><td>2</td><td>TTGAGACCTGAGGACACTAA</td><td>AGG</td><td>-|||||||[||||||-|||||]PAM</td><td>800</td><td>Intronic</td><td>Gpat4</td><td>ENSMUSG00000031545</td></tr>
<tr><td>chr2</td><td>170423716</td><td>170423738</td><td>+</td><td>4</td><td>TTGTGGCCTGAGCAGACTAA</td><td>AGG</td><td>-||-|-||[||||-|||||||]PAM</td><td>3049</td><td>Intronic</td><td>Bcas1</td><td>ENSMUSG00000013523</td></tr>
<tr><td>chr7</td><td>27688502</td><td>27688524</td><td>+</td><td>4</td><td>GGGTGTCCTGAGCAGACTAA</td><td>GGG</td><td>|-|-|-||[||||-|||||||]PAM</td><td>816</td><td>Intergenic</td><td>C030039L03Rik</td><td>ENSMUSG00000057093</td></tr>
<tr><td>chr16</td><td>41683858</td><td>41683880</td><td>+</td><td>4</td><td>TTGAAACCTATGGAGACTAA</td><td>GGG</td><td>-|||-|||[|--|||||||||]PAM</td><td>NA</td><td>Intronic</td><td>NA</td><td>NA</td></tr>
<tr><td>chr16</td><td>11990537</td><td>11990559</td><td>-</td><td>4</td><td>TTGGGATCTGAGAAGACTAA</td><td>AGG</td><td>-||-||-|[||||-|||||||]PAM</td><td>453</td><td>Intronic</td><td>Shisa9</td><td>ENSMUSG00000022494</td></tr>
<tr><td>chr12</td><td>107739235</td><td>107739257</td><td>+</td><td>4</td><td>CAGACACCTGAGGGGACTAA</td><td>GGG</td><td>--||-|||[|||||-||||||]PAM</td><td>3739</td><td>Intergenic</td><td>4930465M20Rik</td><td>ENSMUSG00000085641</td></tr>
<tr><td>chr16</td><td>87538399</td><td>87538421</td><td>-</td><td>4</td><td>TTAACACCTGAGGTGACTAA</td><td>CGG</td><td>-|-|-|||[|||||-||||||]PAM</td><td>10234</td><td>Intergenic</td><td>Gm24891</td><td>ENSMUSG00000077579</td></tr>
<tr><td>chr13</td><td>35167904</td><td>35167926</td><td>+</td><td>3</td><td>GTGATAGCTGAGGTGACTAA</td><td>TGG</td><td>||||-|-|[|||||-||||||]PAM</td><td>12341</td><td>Intergenic</td><td>1700011B04Rik</td><td>ENSMUSG00000102099</td></tr>
<tr><td>chr13</td><td>70148182</td><td>70148204</td><td>-</td><td>4</td><td>GTAAGCACTGAGTAGACTAA</td><td>AGG</td><td>||-||--|[||||-|||||||]PAM</td><td>20628</td><td>Intergenic</td><td>Gm26018</td><td>ENSMUSG00000077594</td></tr>
<tr><td>chr5</td><td>82751462</td><td>82751484</td><td>-</td><td>4</td><td>GAGAGATCAGAAGAGACTAA</td><td>AGG</td><td>|-||||-|[-||-||||||||]PAM</td><td>47460</td><td>Intergenic</td><td>Mir1187</td><td>ENSMUSG00000080586</td></tr>
<tr><td>chr4</td><td>134744133</td><td>134744155</td><td>+</td><td>4</td><td>TTGAGACATCCGGAGACTAA</td><td>AGG</td><td>-||||||-[|--|||||||||]PAM</td><td>0</td><td>Exonic</td><td>Ldlrap1</td><td>ENSMUSG00000037295</td></tr>
<tr><td>chr8</td><td>125481334</td><td>125481356</td><td>-</td><td>4</td><td>GATAGACCTGTGCAGACTAA</td><td>AGG</td><td>|--|||||[||-|-|||||||]PAM</td><td>692</td><td>Intronic</td><td>Sipa1l2</td><td>ENSMUSG00000001995</td></tr>
<tr><td>chr10</td><td>31534480</td><td>31534502</td><td>-</td><td>4</td><td>ATGGGGCCTGAGGATACTAA</td><td>AGG</td><td>-||-|-||[||||||-|||||]PAM</td><td>161</td><td>Intronic</td><td>Rnf217</td><td>ENSMUSG00000063760</td></tr>
<tr><td>chr4</td><td>54942634</td><td>54942656</td><td>+</td><td>4</td><td>CTGAGAGCTGTGCAGACTAA</td><td>GGG</td><td>-|||||-|[||-|-|||||||]PAM</td><td>2392</td><td>Intergenic</td><td>Zfp462</td><td>ENSMUSG00000060206</td></tr>
<tr><td>chr7</td><td>104260002</td><td>104260024</td><td>-</td><td>4</td><td>TTCATACCTGAGGAGCCTAA</td><td>AGG</td><td>-|-|-|||[|||||||-||||]PAM</td><td>94</td><td>Intronic</td><td>Trim34a</td><td>ENSMUSG00000056144</td></tr>
<tr><td>chrX</td><td>98918960</td><td>98918982</td><td>-</td><td>4</td><td>GTGAAAACGGAGGGGACTAA</td><td>TGG</td><td>||||-|-|[-||||-||||||]PAM</td><td>17334</td><td>Intergenic</td><td>Yipf6</td><td>ENSMUSG00000047694</td></tr>
<tr><td>chr6</td><td>85584293</td><td>85584315</td><td>-</td><td>4</td><td>CTGGTACCTGAGGAGGCTAA</td><td>AGG</td><td>-||--|||[|||||||-||||]PAM</td><td>3216</td><td>Intergenic</td><td>Alms1</td><td>ENSMUSG00000063810</td></tr>
<tr><td>chr4</td><td>28717333</td><td>28717355</td><td>-</td><td>4</td><td>TTGAAACCAGAGGACACTAA</td><td>AGG</td><td>-|||-|||[-|||||-|||||]PAM</td><td>95776</td><td>Intergenic</td><td>Epha7</td><td>ENSMUSG00000028289</td></tr>
<tr><td>chr12</td><td>80032211</td><td>80032233</td><td>-</td><td>4</td><td>AGGAGAGCTGAGGAGCCTAA</td><td>AGG</td><td>--||||-|[|||||||-||||]PAM</td><td>68339</td><td>Intergenic</td><td>Gm28056</td><td>ENSMUSG00000101696</td></tr>
<tr><td>chr3</td><td>28537233</td><td>28537255</td><td>-</td><td>4</td><td>GTGAGTGCTTAGGGGACTAA</td><td>AGG</td><td>|||||--|[|-|||-||||||]PAM</td><td>1023</td><td>Intronic</td><td>Tnik</td><td>ENSMUSG00000027692</td></tr>
<tr><td>chr2</td><td>128865814</td><td>128865836</td><td>+</td><td>3</td><td>GTGAAACCAGAGGAGGCTAA</td><td>AGG</td><td>||||-|||[-||||||-||||]PAM</td><td>1667</td><td>Intronic</td><td>Fbln7</td><td>ENSMUSG00000027386</td></tr>
<tr><td>chr6</td><td>117646837</td><td>117646859</td><td>-</td><td>4</td><td>GCCAGACCTGAGCTGACTAA</td><td>AGG</td><td>|--|||||[||||--||||||]PAM</td><td>0</td><td>Exonic</td><td>1700030F04Rik</td><td>ENSMUSG00000099869</td></tr>
<tr><td>chr5</td><td>43164751</td><td>43164773</td><td>-</td><td>4</td><td>GTGGCACCTGTGGACACTAA</td><td>CGG</td><td>|||--|||[||-|||-|||||]PAM</td><td>1146</td><td>Intronic</td><td>Gm7854</td><td>ENSMUSG00000085720</td></tr>
<tr><td>chr17</td><td>7958364</td><td>7958386</td><td>+</td><td>4</td><td>CTGAGCCCTGAGCGGACTAA</td><td>TGG</td><td>-||||-||[||||--||||||]PAM</td><td>5728</td><td>Intronic</td><td>Rsph3a</td><td>ENSMUSG00000073471</td></tr>
<tr><td>chr1</td><td>39351153</td><td>39351175</td><td>-</td><td>4</td><td>GGGAAACCTGAAGACACTAA</td><td>AGG</td><td>|-||-|||[|||-||-|||||]PAM</td><td>2629</td><td>Intronic</td><td>Npas2</td><td>ENSMUSG00000026077</td></tr>
<tr><td>chr9</td><td>69121261</td><td>69121283</td><td>+</td><td>4</td><td>TTGAGAACAGAGGAGTCTAA</td><td>AGG</td><td>-|||||-|[-||||||-||||]PAM</td><td>74258</td><td>Intronic</td><td>Rora</td><td>ENSMUSG00000032238</td></tr>
<tr><td>chr7</td><td>35468059</td><td>35468081</td><td>-</td><td>3</td><td>GTGAGTCCTGAGAACACTAA</td><td>AGG</td><td>|||||-||[||||-|-|||||]PAM</td><td>1017</td><td>Intergenic</td><td>Tdrd12</td><td>ENSMUSG00000030491</td></tr>
<tr><td>chr14</td><td>57532569</td><td>57532591</td><td>+</td><td>4</td><td>CTGTGACCTGAGAATACTAA</td><td>AGG</td><td>-||-||||[||||-|-|||||]PAM</td><td>6970</td><td>Intronic</td><td>Il17d</td><td>ENSMUSG00000050222</td></tr>
<tr><td>chr14</td><td>5161527</td><td>5161549</td><td>-</td><td>4</td><td>CTTAGACTTGAGGAGAATAA</td><td>TGG</td><td>-|-||||-[||||||||-|||]PAM</td><td>0</td><td>Exonic</td><td>Gm26552</td><td>ENSMUSG00000097218</td></tr>
<tr><td>chr14</td><td>5514168</td><td>5514190</td><td>-</td><td>4</td><td>CTTAGACTTGAGGAGAATAA</td><td>TGG</td><td>-|-||||-[||||||||-|||]PAM</td><td>0</td><td>Exonic</td><td>Gm26893</td><td>ENSMUSG00000097450</td></tr>
<tr><td>chr14</td><td>3067074</td><td>3067096</td><td>-</td><td>4</td><td>CTTAGACTTGAGGAGAATAA</td><td>TGG</td><td>-|-||||-[||||||||-|||]PAM</td><td>846</td><td>Intronic</td><td>Gm2897</td><td>ENSMUSG00000079410</td></tr>
<tr><td>chr14</td><td>5800299</td><td>5800321</td><td>+</td><td>4</td><td>CTTAGACTTGAGGAGAATAA</td><td>TGG</td><td>-|-||||-[||||||||-|||]PAM</td><td>1116</td><td>Intronic</td><td>Gm3383</td><td>ENSMUSG00000096629</td></tr>
<tr><td>chr14</td><td>5447808</td><td>5447830</td><td>+</td><td>4</td><td>CTTAGACTTGAGGAGAATAA</td><td>TGG</td><td>-|-||||-[||||||||-|||]PAM</td><td>1116</td><td>Intronic</td><td>Gm3194</td><td>ENSMUSG00000094825</td></tr>
<tr><td>chr14</td><td>5048570</td><td>5048592</td><td>-</td><td>4</td><td>CTTAGACTTGAGGAGAATAA</td><td>TGG</td><td>-|-||||-[||||||||-|||]PAM</td><td>2037</td><td>Intergenic</td><td>4930555G01Rik</td><td>ENSMUSG00000096197</td></tr>
<tr><td>chr14</td><td>3996335</td><td>3996357</td><td>-</td><td>4</td><td>CTTAGACTTGAGGAGAATAA</td><td>TGG</td><td>-|-||||-[||||||||-|||]PAM</td><td>2037</td><td>Intergenic</td><td>Gm3099</td><td>ENSMUSG00000095686</td></tr>
<tr><td>chr14</td><td>7196585</td><td>7196607</td><td>+</td><td>4</td><td>CTTAGACTTGAGGAGAATAA</td><td>TGG</td><td>-|-||||-[||||||||-|||]PAM</td><td>2045</td><td>Intergenic</td><td>Gm16434</td><td>ENSMUSG00000095015</td></tr>
<tr><td>chr14</td><td>7127457</td><td>7127479</td><td>+</td><td>4</td><td>CTTAGACTTGAGGAGAATAA</td><td>TGG</td><td>-|-||||-[||||||||-|||]PAM</td><td>2045</td><td>Intergenic</td><td>Gm6676</td><td>ENSMUSG00000096869</td></tr>
<tr><td>chr14</td><td>4774256</td><td>4774278</td><td>-</td><td>4</td><td>CTTAGACTTGAGGAGAATAA</td><td>TGG</td><td>-|-||||-[||||||||-|||]PAM</td><td>2059</td><td>Intergenic</td><td>Gm9602</td><td>ENSMUSG00000090539</td></tr>
<tr><td>chr14</td><td>6025487</td><td>6025509</td><td>+</td><td>4</td><td>CTTAGACTTGAGGAGAATAA</td><td>TGG</td><td>-|-||||-[||||||||-|||]PAM</td><td>2082</td><td>Intronic</td><td>Gm8206</td><td>ENSMUSG00000091568</td></tr>
<tr><td>chr14</td><td>5337016</td><td>5337038</td><td>+</td><td>4</td><td>CTTAGACTTGAGGAGAATAA</td><td>TGG</td><td>-|-||||-[||||||||-|||]PAM</td><td>2088</td><td>Intergenic</td><td>Gm3542</td><td>ENSMUSG00000096023</td></tr>
<tr><td>chr14</td><td>6221381</td><td>6221403</td><td>+</td><td>4</td><td>CTTAGACTTGAGGAGAATAA</td><td>TGG</td><td>-|-||||-[||||||||-|||]PAM</td><td>2089</td><td>Intergenic</td><td>Gm21560</td><td>ENSMUSG00000094460</td></tr>
<tr><td>chr14</td><td>6539380</td><td>6539402</td><td>+</td><td>4</td><td>CTTAGACTTGAGGAGAATAA</td><td>TGG</td><td>-|-||||-[||||||||-|||]PAM</td><td>2090</td><td>Intergenic</td><td>Gm8356</td><td>ENSMUSG00000090627</td></tr>
<tr><td>chr14</td><td>5689551</td><td>5689573</td><td>+</td><td>4</td><td>CTTAGACTTGAGGAGAATAA</td><td>TGG</td><td>-|-||||-[||||||||-|||]PAM</td><td>2094</td><td>Intronic</td><td>Gm9603</td><td>ENSMUSG00000096470</td></tr>
<tr><td>chr14</td><td>6592726</td><td>6592748</td><td>+</td><td>4</td><td>CTTAGACTTGAGGAGAATAA</td><td>TGG</td><td>-|-||||-[||||||||-|||]PAM</td><td>2113</td><td>Intergenic</td><td>Gm3629</td><td>ENSMUSG00000093898</td></tr>
<tr><td>chr14</td><td>6842425</td><td>6842447</td><td>+</td><td>4</td><td>CTTAGACTTGAGGAGAATAA</td><td>TGG</td><td>-|-||||-[||||||||-|||]PAM</td><td>2114</td><td>Intergenic</td><td>Gm3642</td><td>ENSMUSG00000096685</td></tr>
<tr><td>chr14</td><td>7554897</td><td>7554919</td><td>+</td><td>4</td><td>CTTAGACTTGAGGAGAATAA</td><td>TGG</td><td>-|-||||-[||||||||-|||]PAM</td><td>2136</td><td>Intronic</td><td>Gm3558</td><td>ENSMUSG00000079364</td></tr>
<tr><td>chr14</td><td>3961389</td><td>3961411</td><td>-</td><td>4</td><td>CTTAGACTTGAGGAGAATAA</td><td>TGG</td><td>-|-||||-[||||||||-|||]PAM</td><td>2136</td><td>Intronic</td><td>Gm3095</td><td>ENSMUSG00000091756</td></tr>
<tr><td>chr14</td><td>4738676</td><td>4738698</td><td>-</td><td>4</td><td>CTTAGACTTGAGGAGAATAA</td><td>TGG</td><td>-|-||||-[||||||||-|||]PAM</td><td>2137</td><td>Intronic</td><td>Gm3252</td><td>ENSMUSG00000094628</td></tr>
<tr><td>chr14</td><td>6174141</td><td>6174163</td><td>+</td><td>4</td><td>CTTAGACTTGAGGAGAATAA</td><td>TGG</td><td>-|-||||-[||||||||-|||]PAM</td><td>2140</td><td>Intergenic</td><td>Gm26991</td><td>ENSMUSG00000097199</td></tr>
<tr><td>chr14</td><td>6096090</td><td>6096112</td><td>+</td><td>4</td><td>CTTAGACTTGAGGAGAATAA</td><td>TGG</td><td>-|-||||-[||||||||-|||]PAM</td><td>2143</td><td>Intronic</td><td>Gm3468</td><td>ENSMUSG00000094634</td></tr>
<tr><td>chr14</td><td>5948060</td><td>5948082</td><td>+</td><td>4</td><td>CTTAGACTTGAGGAGAATAA</td><td>TGG</td><td>-|-||||-[||||||||-|||]PAM</td><td>2143</td><td>Intronic</td><td>Gm3248</td><td>ENSMUSG00000091275</td></tr>
<tr><td>chr14</td><td>7231282</td><td>7231304</td><td>+</td><td>4</td><td>CTTAGACTTGAGGAGAATAA</td><td>TGG</td><td>-|-||||-[||||||||-|||]PAM</td><td>2144</td><td>Intergenic</td><td>Gm3715</td><td>ENSMUSG00000072731</td></tr>
<tr><td>chr14</td><td>7092962</td><td>7092984</td><td>+</td><td>4</td><td>CTTAGACTTGAGGAGAATAA</td><td>TGG</td><td>-|-||||-[||||||||-|||]PAM</td><td>2145</td><td>Intronic</td><td>Gm3696</td><td>ENSMUSG00000092167</td></tr>
<tr><td>chr14</td><td>5083539</td><td>5083561</td><td>-</td><td>4</td><td>CTTAGACTTGAGGAGAATAA</td><td>TGG</td><td>-|-||||-[||||||||-|||]PAM</td><td>2148</td><td>Intronic</td><td>Gm8281</td><td>ENSMUSG00000095681</td></tr>
<tr><td>chr14</td><td>4031581</td><td>4031603</td><td>-</td><td>4</td><td>CTTAGACTTGAGGAGAATAA</td><td>TGG</td><td>-|-||||-[||||||||-|||]PAM</td><td>2151</td><td>Intronic</td><td>Gm5796</td><td>ENSMUSG00000096775</td></tr>
<tr><td>chr14</td><td>3131823</td><td>3131845</td><td>-</td><td>4</td><td>CTTAGACTTGAGGAGAATAA</td><td>TGG</td><td>-|-||||-[||||||||-|||]PAM</td><td>2152</td><td>Intergenic</td><td>Gm10340</td><td>ENSMUSG00000071613</td></tr>
<tr><td>chr14</td><td>5728101</td><td>5728123</td><td>+</td><td>4</td><td>CTTAGACTTGAGGAGAATAA</td><td>TGG</td><td>-|-||||-[||||||||-|||]PAM</td><td>2153</td><td>Intronic</td><td>Gm3373</td><td>ENSMUSG00000094370</td></tr>
<tr><td>chr14</td><td>4662016</td><td>4662038</td><td>-</td><td>4</td><td>CTTAGACTTGAGGAGAATAA</td><td>TGG</td><td>-|-||||-[||||||||-|||]PAM</td><td>2154</td><td>Intergenic</td><td>Gm3239</td><td>ENSMUSG00000090872</td></tr>
<tr><td>chr14</td><td>5866722</td><td>5866744</td><td>+</td><td>4</td><td>CTTAGACTTGAGGAGAATAA</td><td>TGG</td><td>-|-||||-[||||||||-|||]PAM</td><td>2157</td><td>Intronic</td><td>Gm8237</td><td>ENSMUSG00000090707</td></tr>
<tr><td>chr14</td><td>5375565</td><td>5375587</td><td>+</td><td>4</td><td>CTTAGACTTGAGGAGAATAA</td><td>TGG</td><td>-|-||||-[||||||||-|||]PAM</td><td>2159</td><td>Intronic</td><td>Gm3500</td><td>ENSMUSG00000096003</td></tr>
<tr><td>chr14</td><td>4247450</td><td>4247472</td><td>-</td><td>4</td><td>CTTAGACTTGAGGAGAATAA</td><td>TGG</td><td>-|-||||-[||||||||-|||]PAM</td><td>2159</td><td>Intergenic</td><td>Gm3138</td><td>ENSMUSG00000091114</td></tr>
<tr><td>chr14</td><td>7014931</td><td>7014953</td><td>+</td><td>4</td><td>CTTAGACTTGAGGAGAATAA</td><td>TGG</td><td>-|-||||-[||||||||-|||]PAM</td><td>2160</td><td>Intronic</td><td>Gm10406</td><td>ENSMUSG00000093985</td></tr>
<tr><td>chr14</td><td>5013693</td><td>5013715</td><td>-</td><td>4</td><td>CTTAGACTTGAGGAGAATAA</td><td>TGG</td><td>-|-||||-[||||||||-|||]PAM</td><td>2176</td><td>Intergenic</td><td>Gm3298</td><td>ENSMUSG00000079383</td></tr>
<tr><td>chr14</td><td>7302545</td><td>7302567</td><td>+</td><td>4</td><td>CTTAGACTTGAGGAGAATAA</td><td>TGG</td><td>-|-||||-[||||||||-|||]PAM</td><td>2188</td><td>Intronic</td><td>Gm3739</td><td>ENSMUSG00000091472</td></tr>
<tr><td>chr14</td><td>6877350</td><td>6877372</td><td>+</td><td>4</td><td>CTTAGACTTGAGGAGAATAA</td><td>TGG</td><td>-|-||||-[||||||||-|||]PAM</td><td>2190</td><td>Intronic</td><td>Gm3667</td><td>ENSMUSG00000090691</td></tr>
<tr><td>chr14</td><td>7266967</td><td>7266989</td><td>+</td><td>4</td><td>CTTAGACTTGAGGAGAATAA</td><td>TGG</td><td>-|-||||-[||||||||-|||]PAM</td><td>2253</td><td>Intergenic</td><td>Gm3727</td><td>ENSMUSG00000096183</td></tr>
<tr><td>chr14</td><td>6060521</td><td>6060543</td><td>+</td><td>4</td><td>CTTAGACTTGAGGAGAATAA</td><td>TGG</td><td>-|-||||-[||||||||-|||]PAM</td><td>2254</td><td>Intronic</td><td>Gm6337</td><td>ENSMUSG00000091882</td></tr>
<tr><td>chr14</td><td>4081839</td><td>4081861</td><td>-</td><td>4</td><td>CTTAGACTTGAGGAGAATAA</td><td>TGG</td><td>-|-||||-[||||||||-|||]PAM</td><td>2259</td><td>Intergenic</td><td>Gm3115</td><td>ENSMUSG00000091022</td></tr>
<tr><td>chr14</td><td>5282940</td><td>5282962</td><td>+</td><td>4</td><td>CTTAGACTTGAGGAGAATAA</td><td>TGG</td><td>-|-||||-[||||||||-|||]PAM</td><td>2418</td><td>Intergenic</td><td>Gm3339</td><td>ENSMUSG00000091995</td></tr>
<tr><td>chr14</td><td>5635446</td><td>5635468</td><td>+</td><td>4</td><td>CTTAGACTTGAGGAGAATAA</td><td>TGG</td><td>-|-||||-[||||||||-|||]PAM</td><td>2418</td><td>Intronic</td><td>Gm8265</td><td>ENSMUSG00000090547</td></tr>
<tr><td>chr14</td><td>7162289</td><td>7162311</td><td>+</td><td>4</td><td>CTTAGACTTGAGGAGAATAA</td><td>TGG</td><td>-|-||||-[||||||||-|||]PAM</td><td>2903</td><td>Intronic</td><td>Gm3512</td><td>ENSMUSG00000090363</td></tr>
<tr><td>chr14</td><td>6269655</td><td>6269677</td><td>+</td><td>4</td><td>CTTAGACTTGAGGAGAATAA</td><td>TGG</td><td>-|-||||-[||||||||-|||]PAM</td><td>3000</td><td>Intronic</td><td>Gm3411</td><td>ENSMUSG00000079396</td></tr>
<tr><td>chr14</td><td>3062584</td><td>3062606</td><td>-</td><td>4</td><td>CTTAGACTTGAGGAGAATAA</td><td>TGG</td><td>-|-||||-[||||||||-|||]PAM</td><td>3460</td><td>Intronic</td><td>Gm2897</td><td>ENSMUSG00000079410</td></tr>
<tr><td>chr1</td><td>72142980</td><td>72143002</td><td>-</td><td>3</td><td>GTGTGACCCGAGGAGAGTAA</td><td>GGG</td><td>|||-||||[-|||||||-|||]PAM</td><td>16440</td><td>Intergenic</td><td>Mreg</td><td>ENSMUSG00000039395</td></tr>
<tr><td>chr17</td><td>63548024</td><td>63548046</td><td>+</td><td>4</td><td>GTCAGAACTGATGAAACTAA</td><td>AGG</td><td>||-|||-|[|||-||-|||||]PAM</td><td>48007</td><td>Intergenic</td><td>Fbxl17</td><td>ENSMUSG00000023965</td></tr>
<tr><td>chr6</td><td>35086386</td><td>35086408</td><td>-</td><td>4</td><td>TGGAGACCAGAGGAGAGTAA</td><td>TGG</td><td>--||||||[-|||||||-|||]PAM</td><td>6160</td><td>Intronic</td><td>Cnot4</td><td>ENSMUSG00000038784</td></tr>
<tr><td>chrX</td><td>100578121</td><td>100578143</td><td>-</td><td>4</td><td>GTGTGACAAGAGGAGTCTAA</td><td>GGG</td><td>|||-|||-[-||||||-||||]PAM</td><td>0</td><td>Exonic</td><td>Awat1</td><td>ENSMUSG00000015665</td></tr>
<tr><td>chr4</td><td>11253491</td><td>11253513</td><td>+</td><td>4</td><td>ATGAGAACTGAGCAAACTAA</td><td>AGG</td><td>-|||||-|[||||-|-|||||]PAM</td><td>468</td><td>Intronic</td><td>Ints8</td><td>ENSMUSG00000040738</td></tr>
<tr><td>chr2</td><td>25995838</td><td>25995860</td><td>-</td><td>3</td><td>ATGAGAGCTGAGGAGACCAA</td><td>TGG</td><td>-|||||-|[|||||||||-||]PAM</td><td>655</td><td>Intergenic</td><td>Gm13542</td><td>ENSMUSG00000082357</td></tr>
<tr><td>chr19</td><td>56329893</td><td>56329915</td><td>-</td><td>4</td><td>CTGAGGCCCGAGGAGATTAA</td><td>AGG</td><td>-||||-||[-|||||||-|||]PAM</td><td>793</td><td>Intronic</td><td>Nrap</td><td>ENSMUSG00000049134</td></tr>
<tr><td>chr14</td><td>26302469</td><td>26302491</td><td>+</td><td>3</td><td>GAGAGAACTGAGGAGACCAA</td><td>AGG</td><td>|-||||-|[|||||||||-||]PAM</td><td>33033</td><td>Intergenic</td><td>Duxbl3</td><td>ENSMUSG00000072672</td></tr>
<tr><td>chr5</td><td>91623780</td><td>91623802</td><td>+</td><td>4</td><td>GAGAGACATGCGGAGTCTAA</td><td>GGG</td><td>|-|||||-[||-||||-||||]PAM</td><td>0</td><td>Exonic</td><td>Parm1</td><td>ENSMUSG00000034981</td></tr>
<tr><td>chr3</td><td>33666045</td><td>33666067</td><td>-</td><td>4</td><td>TGGAGACCTGAGTAGCCTAA</td><td>GGG</td><td>--||||||[||||-||-||||]PAM</td><td>18441</td><td>Intergenic</td><td>Gm42694</td><td>ENSMUSG00000104954</td></tr>
<tr><td>chr11</td><td>56625280</td><td>56625302</td><td>+</td><td>4</td><td>GGGAGAGCAGAGGAGAGTAA</td><td>TGG</td><td>|-||||-|[-|||||||-|||]PAM</td><td>NA</td><td>Intergenic</td><td>NA</td><td>NA</td></tr>
<tr><td>chr5</td><td>14826936</td><td>14826958</td><td>+</td><td>4</td><td>GTGACATGTGAGGAGAATAA</td><td>AGG</td><td>||||-|--[||||||||-|||]PAM</td><td>9602</td><td>Intronic</td><td>Pclo</td><td>ENSMUSG00000061601</td></tr>
<tr><td>chr5</td><td>12696583</td><td>12696605</td><td>-</td><td>4</td><td>GGGAGGCCTGAGGTAACTAA</td><td>AGG</td><td>|-|||-||[|||||--|||||]PAM</td><td>25855</td><td>Intergenic</td><td>Gm42580</td><td>ENSMUSG00000105237</td></tr>
<tr><td>chr2</td><td>10674978</td><td>10675000</td><td>+</td><td>4</td><td>GTGTGAACGGAGGAGAATAA</td><td>AGG</td><td>|||-||-|[-|||||||-|||]PAM</td><td>79725</td><td>Intergenic</td><td>Sfmbt2</td><td>ENSMUSG00000061186</td></tr>
<tr><td>chr3</td><td>107581911</td><td>107581933</td><td>-</td><td>4</td><td>AGGAGACTTGAGGAGACAAA</td><td>AGG</td><td>--|||||-[|||||||||-||]PAM</td><td>13098</td><td>Intergenic</td><td>Alx3</td><td>ENSMUSG00000014603</td></tr>
<tr><td>chr14</td><td>86680929</td><td>86680951</td><td>+</td><td>4</td><td>GAGGGATCTGAGGAGACCAA</td><td>TGG</td><td>|-|-||-|[|||||||||-||]PAM</td><td>24347</td><td>Intronic</td><td>Diap3</td><td>ENSMUSG00000022021</td></tr>
</table>

<table>
<tr><td>chr11</td><td>3532597</td><td>3532619</td><td>-</td><td>4</td><td>GTGCGAGCTCAGGAGATTAA</td><td>GGG</td><td>|||-||-|[|-||||||-|||]PAM</td><td>0</td><td>Exonic</td><td>Smtn</td><td>ENSMUSG00000020439</td></tr>
<tr><td>chr15</td><td>68535087</td><td>68535109</td><td>-</td><td>4</td><td>GTGACAGCTGAGGTCACTAA</td><td>AGG</td><td>||||-|-|[|||||--|||||]PAM</td><td>NA</td><td>Intergenic</td><td>NA</td><td>NA</td></tr>
<tr><td>chr9</td><td>51388321</td><td>51388343</td><td>+</td><td>4</td><td>CTGAGAACTGGGGAGAGTAA</td><td>TGG</td><td>-|||||-|[||-|||||-|||]PAM</td><td>59404</td><td>Intergenic</td><td>1810046K07Rik</td><td>ENSMUSG00000036027</td></tr>
<tr><td>chr17</td><td>73382442</td><td>73382464</td><td>+</td><td>4</td><td>CTGACACCTGAAGAGAATAA</td><td>AGG</td><td>-|||-|||[|||-||||-|||]PAM</td><td>359</td><td>Intronic</td><td>Capn13</td><td>ENSMUSG00000043705</td></tr>
<tr><td>chr18</td><td>31935326</td><td>31935348</td><td>-</td><td>4</td><td>CTGAGTCCAGAGGAGACAAA</td><td>GGG</td><td>-||||-||[-||||||||-||]PAM</td><td>3776</td><td>Intronic</td><td>Lims2</td><td>ENSMUSG00000024395</td></tr>
<tr><td>chr6</td><td>131199347</td><td>131199369</td><td>-</td><td>3</td><td>GTCAGGCCTGAGGAGACTCA</td><td>GGG</td><td>||-||-||[||||||||||-|]PAM</td><td>16968</td><td>Intergenic</td><td>Gm5581</td><td>ENSMUSG00000061969</td></tr>
<tr><td>chr15</td><td>34338100</td><td>34338122</td><td>-</td><td>3</td><td>ATGAGATCTGAGGAGACTGA</td><td>AGG</td><td>-|||||-|[||||||||||-|]PAM</td><td>6869</td><td>Intronic</td><td>Matn2</td><td>ENSMUSG00000022324</td></tr>
<tr><td>chr1</td><td>172221686</td><td>172221708</td><td>-</td><td>4</td><td>GTGGGAGCTGAAGAGAGTAA</td><td>GGG</td><td>|||-||-|[|||-||||-|||]PAM</td><td>1805</td><td>Intergenic</td><td>Atp1a4</td><td>ENSMUSG00000007107</td></tr>
<tr><td>chr12</td><td>12267469</td><td>12267491</td><td>+</td><td>3</td><td>GAGAGATCTGAGGAGACTCA</td><td>AGG</td><td>|-||||-|[||||||||||-|]PAM</td><td>5231</td><td>Intronic</td><td>Fam49a</td><td>ENSMUSG00000020589</td></tr>
<tr><td>chr9</td><td>112251216</td><td>112251238</td><td>+</td><td>4</td><td>GGAAGGCCTGAGGAGACTTA</td><td>AGG</td><td>|--||-||[||||||||||-|]PAM</td><td>5336</td><td>Intronic</td><td>2900079G21Rik</td><td>ENSMUSG00000087038</td></tr>
<tr><td>chr6</td><td>130407850</td><td>130407872</td><td>-</td><td>3</td><td>GTGAGTACTGAGGAGACTCA</td><td>AGG</td><td>|||||--|[||||||||||-|]PAM</td><td>0</td><td>Exonic</td><td>Gm6584</td><td>ENSMUSG00000084204</td></tr>
<tr><td>chr13</td><td>54451490</td><td>54451512</td><td>+</td><td>3</td><td>CTGAGACCTGAGAAGACCAA</td><td>AGG</td><td>-|||||||[||||-||||-||]PAM</td><td>7325</td><td>Intergenic</td><td>Thoc3</td><td>ENSMUSG00000025872</td></tr>
<tr><td>chr15</td><td>84199816</td><td>84199838</td><td>+</td><td>4</td><td>GACAGAGCTGAGGAGACTGA</td><td>CGG</td><td>|--|||-|[||||||||||-|]PAM</td><td>0</td><td>Exonic</td><td>Samm50</td><td>ENSMUSG00000022437</td></tr>
<tr><td>chr7</td><td>28090275</td><td>28090297</td><td>+</td><td>4</td><td>GGAAGAGCTGAGGAGACTGA</td><td>AGG</td><td>|--|||-|[||||||||||-|]PAM</td><td>116</td><td>Intronic</td><td>Fcgbp</td><td>ENSMUSG00000047730</td></tr>
<tr><td>chr1</td><td>152810119</td><td>152810141</td><td>+</td><td>4</td><td>CTGAGAGCTGTGGAGACAAA</td><td>CGG</td><td>-|||||-|[||-||||||-||]PAM</td><td>0</td><td>Exonic</td><td>Ncf2</td><td>ENSMUSG00000026480</td></tr>
<tr><td>chr10</td><td>122178953</td><td>122178975</td><td>-</td><td>4</td><td>GTGAAACTTTAGGAGACAAA</td><td>TGG</td><td>||||-||-[|-|||||||-||]PAM</td><td>8590</td><td>Intergenic</td><td>Gm9081</td><td>ENSMUSG00000098138</td></tr>
<tr><td>chr3</td><td>118495998</td><td>118496020</td><td>-</td><td>4</td><td>GGGAGAGCTGAAGAGACCAA</td><td>GGG</td><td>|-||||-|[|||-|||||-||]PAM</td><td>18118</td><td>Intronic</td><td>Gm26871</td><td>ENSMUSG00000097311</td></tr>
<tr><td>chr1</td><td>157408304</td><td>157408326</td><td>+</td><td>4</td><td>GCTAGACCTGAGGACAATAA</td><td>AGG</td><td>|--|||||[||||||-|-|||]PAM</td><td>1583</td><td>Intronic</td><td>Gm37060</td><td>ENSMUSG00000103808</td></tr>
<tr><td>chr16</td><td>50205925</td><td>50205947</td><td>+</td><td>4</td><td>GGAAGACCTGAGGACAGTAA</td><td>GGG</td><td>|--|||||[||||||-|-|||]PAM</td><td>3138</td><td>Intronic</td><td>Bbx</td><td>ENSMUSG00000022641</td></tr>
<tr><td>chr17</td><td>19354929</td><td>19354951</td><td>-</td><td>4</td><td>TTGGGACCTGAGGAAAGTAA</td><td>GGG</td><td>-||-||||[||||||-|-|||]PAM</td><td>7184</td><td>Intergenic</td><td>Vmn2r99</td><td>ENSMUSG00000090304</td></tr>
<tr><td>chr10</td><td>35173361</td><td>35173383</td><td>-</td><td>4</td><td>TTGGGACCTGAGGAAAGTAA</td><td>GGG</td><td>-||-||||[||||||-|-|||]PAM</td><td>61030</td><td>Intergenic</td><td>Gm26341</td><td>ENSMUSG00000077749</td></tr>
<tr><td>chr6</td><td>130335711</td><td>130335733</td><td>-</td><td>4</td><td>GTAAGGCATGAGGAGACTCA</td><td>AGG</td><td>||-||-|-[||||||||||-|]PAM</td><td>0</td><td>Exonic</td><td>Klra3</td><td>ENSMUSG00000067591</td></tr>
<tr><td>chr6</td><td>130025131</td><td>130025153</td><td>-</td><td>4</td><td>GTAAGGCATGAGGAGACTCA</td><td>AGG</td><td>||-||-|-[||||||||||-|]PAM</td><td>0</td><td>Exonic</td><td>Klra6</td><td>ENSMUSG00000061769</td></tr>
<tr><td>chr6</td><td>130191213</td><td>130191235</td><td>-</td><td>4</td><td>GTAAGGCATGAGGAGACTCA</td><td>AGG</td><td>||-||-|-[||||||||||-|]PAM</td><td>0</td><td>Exonic</td><td>Klra9</td><td>ENSMUSG00000033024</td></tr>
<tr><td>chr6</td><td>130281834</td><td>130281856</td><td>-</td><td>4</td><td>GTAAGGCATGAGGAGACTCA</td><td>AGG</td><td>||-||-|-[||||||||||-|]PAM</td><td>0</td><td>Exonic</td><td>Klra10</td><td>ENSMUSG00000072718</td></tr>
<tr><td>chr5</td><td>127632065</td><td>127632087</td><td>-</td><td>4</td><td>AGGGGACCTGAGGAGACTAG</td><td>CGG</td><td>--|-||||[|||||||||||-]PAM</td><td>175</td><td>Intronic</td><td>Glt1d1</td><td>ENSMUSG00000049971</td></tr>
<tr><td>chr8</td><td>39011657</td><td>39011679</td><td>-</td><td>4</td><td>CTGAGACATGAGGAAAATAA</td><td>AGG</td><td>-||||||-[||||||-|-|||]PAM</td><td>5457</td><td>Intronic</td><td>Tusc3</td><td>ENSMUSG00000039530</td></tr>
<tr><td>chr16</td><td>38494470</td><td>38494492</td><td>-</td><td>3</td><td>ATGAGACCTGAGGAAACAAA</td><td>AGG</td><td>-|||||||[||||||-||-||]PAM</td><td>3855</td><td>Intergenic</td><td>Timmdc1</td><td>ENSMUSG00000002846</td></tr>
<tr><td>chr6</td><td>94043003</td><td>94043025</td><td>-</td><td>4</td><td>AAGAGACCTGAGGAGGTTAA</td><td>TGG</td><td>--||||||[|||||||--|||]PAM</td><td>89898</td><td>Intronic</td><td>Magi1</td><td>ENSMUSG00000045095</td></tr>
<tr><td>chr11</td><td>77522525</td><td>77522547</td><td>-</td><td>4</td><td>GTGAGAAATGAAGAGACCAA</td><td>CGG</td><td>||||||--[|||-|||||-||]PAM</td><td>2480</td><td>Intronic</td><td>Abhd15</td><td>ENSMUSG00000000686</td></tr>
<tr><td>chr16</td><td>37140280</td><td>37140302</td><td>-</td><td>4</td><td>GTTAGAACTCAGGAGACTGA</td><td>AGG</td><td>||-|||-|[|-||||||||-|]PAM</td><td>197</td><td>Intronic</td><td>Stxbp5l</td><td>ENSMUSG00000022829</td></tr>
<tr><td>chr16</td><td>36677736</td><td>36677758</td><td>-</td><td>4</td><td>GTGTGTCCTGTGGAGACTGA</td><td>AGG</td><td>|||-|-||[||-|||||||-|]PAM</td><td>11655</td><td>Intergenic</td><td>Cd86</td><td>ENSMUSG00000022901</td></tr>
<tr><td>chr14</td><td>32309064</td><td>32309086</td><td>-</td><td>4</td><td>AAGAGACCTGAGAAGACTGA</td><td>AGG</td><td>--||||||[||||-|||||-|]PAM</td><td>11514</td><td>Intergenic</td><td>Parg</td><td>ENSMUSG00000021911</td></tr>
<tr><td>chr13</td><td>35532501</td><td>35532523</td><td>+</td><td>4</td><td>GTCAGACATGAGGGGACAAA</td><td>GGG</td><td>||-||||-[|||||-|||-||]PAM</td><td>NA</td><td>Intergenic</td><td>NA</td><td>NA</td></tr>
<tr><td>chr13</td><td>69971320</td><td>69971342</td><td>-</td><td>4</td><td>GAAAGACCTGAGCAGACTGA</td><td>AGG</td><td>|--|||||[||||-|||||-|]PAM</td><td>32708</td><td>Intergenic</td><td>1700084F23Rik</td><td>ENSMUSG00000099639</td></tr>
<tr><td>chr5</td><td>128370984</td><td>128371006</td><td>-</td><td>3</td><td>GAGAGACCTGTGGAGACTAG</td><td>GGG</td><td>|-||||||[||-||||||||-]PAM</td><td>28206</td><td>Intronic</td><td>Gm42817</td><td>ENSMUSG00000105857</td></tr>
<tr><td>chr13</td><td>43304687</td><td>43304709</td><td>+</td><td>4</td><td>GTGAGCTCTGAGGAGGTTAA</td><td>GGG</td><td>|||||--|[|||||||--|||]PAM</td><td>515</td><td>Intergenic</td><td>Gfod1</td><td>ENSMUSG00000051335</td></tr>
<tr><td>chr16</td><td>86895007</td><td>86895029</td><td>-</td><td>4</td><td>GTTAGAAGTGAGGAGACTAT</td><td>GGG</td><td>||-|||--[|||||||||||-]PAM</td><td>49413</td><td>Intergenic</td><td>Gm25715</td><td>ENSMUSG00000089252</td></tr>
<tr><td>chr7</td><td>87396888</td><td>87396910</td><td>+</td><td>4</td><td>CTGAAACCTTAGGAGACTAC</td><td>TGG</td><td>-|||-|||[|-|||||||||-]PAM</td><td>0</td><td>Exonic</td><td>Nox4</td><td>ENSMUSG00000030562</td></tr>
<tr><td>chr1</td><td>85749380</td><td>85749402</td><td>+</td><td>4</td><td>GGAAGACCTGAGGAGTCCAA</td><td>AGG</td><td>|--|||||[|||||||-|-||]PAM</td><td>2546</td><td>Intronic</td><td>Mir8096</td><td>ENSMUSG00000098634</td></tr>
<tr><td>chr4</td><td>61891846</td><td>61891868</td><td>+</td><td>4</td><td>GTGTGACATGAGGACACAAA</td><td>AGG</td><td>|||-|||-[||||||-||-||]PAM</td><td>8550</td><td>Intergenic</td><td>Mup-ps18</td><td>ENSMUSG00000082363</td></tr>
<tr><td>chr12</td><td>31406970</td><td>31406992</td><td>+</td><td>4</td><td>ATGAAACCTGAGGAGGCAAA</td><td>AGG</td><td>-|||-|||[|||||||-|-||]PAM</td><td>11856</td><td>Intronic</td><td>Gm9359</td><td>ENSMUSG00000091040</td></tr>
<tr><td>chr5</td><td>140217862</td><td>140217884</td><td>-</td><td>4</td><td>CTGAGAACTGAGGAGGCCAA</td><td>AGG</td><td>-|||||-|[|||||||-|-||]PAM</td><td>7111</td><td>Intronic</td><td>Gm16120</td><td>ENSMUSG00000087248</td></tr>
<tr><td>chr12</td><td>20955983</td><td>20956005</td><td>-</td><td>4</td><td>TTGGGACCTGAGGAAACTGA</td><td>GGG</td><td>-||-||||[||||||-|||-|]PAM</td><td>34454</td><td>Intergenic</td><td>Gm21989</td><td>ENSMUSG00000090370</td></tr>
<tr><td>chr12</td><td>24131949</td><td>24131971</td><td>+</td><td>4</td><td>TTGGGACCTGAGGAAACTGA</td><td>GGG</td><td>-||-||||[||||||-|||-|]PAM</td><td>34981</td><td>Intergenic</td><td>9030624G23Rik</td><td>ENSMUSG00000073158</td></tr>
<tr><td>chr12</td><td>18817925</td><td>18817947</td><td>-</td><td>4</td><td>TTGGGACCTGAGGAAACTGA</td><td>GGG</td><td>-||-||||[||||||-|||-|]PAM</td><td>35925</td><td>Intergenic</td><td>Gm3944</td><td>ENSMUSG00000094947</td></tr>
<tr><td>chr12</td><td>22782474</td><td>22782496</td><td>+</td><td>4</td><td>TTGGGACCTGAGGAAACTGA</td><td>GGG</td><td>-||-||||[||||||-|||-|]PAM</td><td>NA</td><td>Intergenic</td><td>NA</td><td>NA</td></tr>
<tr><td>chr12</td><td>20578066</td><td>20578088</td><td>-</td><td>4</td><td>TTGGGACCTGAGGAAACTGA</td><td>GGG</td><td>-||-||||[||||||-|||-|]PAM</td><td>NA</td><td>Intergenic</td><td>NA</td><td>NA</td></tr>
<tr><td>chr1</td><td>187036689</td><td>187036711</td><td>+</td><td>4</td><td>GGGAGACTTGAGGAGCCCAA</td><td>AGG</td><td>|-|||||-[|||||||-|-||]PAM</td><td>7937</td><td>Intergenic</td><td>Spata17</td><td>ENSMUSG00000026611</td></tr>
<tr><td>chr13</td><td>84165301</td><td>84165323</td><td>-</td><td>4</td><td>GCAAGACCTGAGGAGAGAAA</td><td>TGG</td><td>|--|||||[||||||||--||]PAM</td><td>16955</td><td>Intergenic</td><td>Gm24331</td><td>ENSMUSG00000092840</td></tr>
<tr><td>chr3</td><td>84180874</td><td>84180896</td><td>+</td><td>4</td><td>GAGACACCTGAGGAGAGCAA</td><td>AGG</td><td>|-||-|||[||||||||--||]PAM</td><td>2696</td><td>Intronic</td><td>Trim2</td><td>ENSMUSG00000027993</td></tr>
<tr><td>chr16</td><td>85272004</td><td>85272026</td><td>-</td><td>4</td><td>GTGGGCCCTGAGGAGAAGAA</td><td>GGG</td><td>|||-|-||[||||||||--||]PAM</td><td>98297</td><td>Intergenic</td><td>App</td><td>ENSMUSG00000022892</td></tr>
<tr><td>chr7</td><td>103730238</td><td>103730260</td><td>+</td><td>4</td><td>GTGATACGTGAAGAGACTAT</td><td>GGG</td><td>||||-||-[|||-|||||||-]PAM</td><td>1668</td><td>Intergenic</td><td>Olfr628</td><td>ENSMUSG00000096516</td></tr>
<tr><td>chr17</td><td>4518337</td><td>4518359</td><td>+</td><td>4</td><td>TTGAGACATGAGGAGAAGAA</td><td>TGG</td><td>-||||||-[||||||||--||]PAM</td><td>NA</td><td>Intergenic</td><td>NA</td><td>NA</td></tr>
<tr><td>chr10</td><td>46268660</td><td>46268682</td><td>-</td><td>4</td><td>CTGAGACATGAGGAGAGCAA</td><td>AGG</td><td>-||||||-[||||||||--||]PAM</td><td>NA</td><td>Intergenic</td><td>NA</td><td>NA</td></tr>
<tr><td>chr9</td><td>86764791</td><td>86764813</td><td>+</td><td>4</td><td>ATGTGACCTGAGGGGACTAG</td><td>AGG</td><td>-||-||||[|||||-|||||-]PAM</td><td>1110</td><td>Intergenic</td><td>Snap91</td><td>ENSMUSG00000033419</td></tr>
<tr><td>chr11</td><td>84914210</td><td>84914232</td><td>-</td><td>4</td><td>GTGAGGTCTGAGGAGAGCAA</td><td>AGG</td><td>|||||--|[||||||||--||]PAM</td><td>0</td><td>Exonic</td><td>Znhit3</td><td>ENSMUSG00000020526</td></tr>
<tr><td>chr10</td><td>116946597</td><td>116946619</td><td>-</td><td>4</td><td>ATGAGAGCTGAGGGGACTAC</td><td>GGG</td><td>-|||||-|[|||||-|||||-]PAM</td><td>3641</td><td>Intronic</td><td>Rab3ip</td><td>ENSMUSG00000064181</td></tr>
<tr><td>chr6</td><td>32027303</td><td>32027325</td><td>-</td><td>4</td><td>CTGAGGCCTGAGGAGATTCA</td><td>AGG</td><td>-||||-||[||||||||-|-|]PAM</td><td>22921</td><td>Intergenic</td><td>1700012A03Rik</td><td>ENSMUSG00000029766</td></tr>
<tr><td>chr8</td><td>104203710</td><td>104203732</td><td>-</td><td>4</td><td>GTGTGCCCTGAGGAGAATGA</td><td>AGG</td><td>|||-|-||[||||||||-|-|]PAM</td><td>7186</td><td>Intronic</td><td>Bean1</td><td>ENSMUSG00000031872</td></tr>
<tr><td>chr11</td><td>71126363</td><td>71126385</td><td>+</td><td>4</td><td>GTTAGAACTGAGGAGATTTA</td><td>AGG</td><td>||-|||-|[||||||||-|-|]PAM</td><td>2092</td><td>Intronic</td><td>Nlrp1a</td><td>ENSMUSG00000069830</td></tr>
<tr><td>chr6</td><td>129911373</td><td>129911395</td><td>-</td><td>4</td><td>GTGAGCCATGAGGAGATTCA</td><td>AGG</td><td>|||||-|-[||||||||-|-|]PAM</td><td>0</td><td>Exonic</td><td>Klra5</td><td>ENSMUSG00000030173</td></tr>
<tr><td>chr17</td><td>20843007</td><td>20843029</td><td>-</td><td>4</td><td>GGAAGACCTGAGGAGAATAT</td><td>AGG</td><td>|--|||||[||||||||-||-]PAM</td><td>3522</td><td>Intergenic</td><td>Vmn1r230</td><td>ENSMUSG00000045417</td></tr>
<tr><td>chr11</td><td>120870647</td><td>120870669</td><td>+</td><td>4</td><td>GTCAGACATGAGGAGACCCA</td><td>AGG</td><td>||-||||-[|||||||||--|]PAM</td><td>379</td><td>Intronic</td><td>Ccdc57</td><td>ENSMUSG00000048445</td></tr>
<tr><td>chr14</td><td>64170929</td><td>64170951</td><td>-</td><td>4</td><td>GTGAGGACTGAGGAGACAGA</td><td>TGG</td><td>|||||--|[|||||||||--|]PAM</td><td>39477</td><td>Intronic</td><td>Msra</td><td>ENSMUSG00000054733</td></tr>
<tr><td>chr17</td><td>23778766</td><td>23778788</td><td>+</td><td>3</td><td>GGGAGACCTGAGGAGACCAC</td><td>AGG</td><td>|-||||||[|||||||||-|-]PAM</td><td>653</td><td>Intronic</td><td>Flywch2</td><td>ENSMUSG00000023911</td></tr>
<tr><td>chr1</td><td>62808403</td><td>62808425</td><td>+</td><td>3</td><td>GGGAGACCTGAGGAGACAAG</td><td>GGG</td><td>|-||||||[|||||||||-|-]PAM</td><td>877</td><td>Intronic</td><td>Gm37121</td><td>ENSMUSG00000103032</td></tr>
<tr><td>chr4</td><td>137564754</td><td>137564776</td><td>-</td><td>4</td><td>GCCAGACCTGAGGAGACCAT</td><td>AGG</td><td>|--|||||[|||||||||-|-]PAM</td><td>0</td><td>Exonic</td><td>Hspg2</td><td>ENSMUSG00000028763</td></tr>
<tr><td>chr16</td><td>64365597</td><td>64365619</td><td>+</td><td>4</td><td>CTCAGACCTGAGGAGACTCC</td><td>AGG</td><td>-|-|||||[||||||||||--]PAM</td><td>NA</td><td>Intergenic</td><td>NA</td><td>NA</td></tr>
<tr><td>chr3</td><td>158091937</td><td>158091959</td><td>-</td><td>4</td><td>GGGAGAACTGAGGAGACTCC</td><td>AGG</td><td>|-||||-|[||||||||||--]PAM</td><td>7391</td><td>Intronic</td><td>Lrrc7</td><td>ENSMUSG00000028176</td></tr>
<tr><td>chr5</td><td>100679827</td><td>100679849</td><td>+</td><td>4</td><td>GTAAGACTTGAGGAGACTTG</td><td>AGG</td><td>||-||||-[||||||||||--]PAM</td><td>0</td><td>Exonic</td><td>Hpse</td><td>ENSMUSG00000035273</td></tr>
</table>

Species: Mouse (Mus musculus GRCm38/mm10)
Input: CCTTGTGAGCTCTATGCTGTACC
PAM: NGG
Target site length: 20
Core length: 12
Core MM: 2
Total MM: 4

<table>
<tr><td>T1</td><td>GGTACAGCATAGAGCTCACAAGG</td><td>1000</td><td>CRISPRater score</td><td>0.54546908</td><td></td><td></td><td></td><td></td><td></td><td></td><td></td></tr>
<tr><td>Chromosome</td><td>start</td><td>end</td><td>strand</td><td>MM</td><td>target_seq</td><td>PAM</td><td>alignment</td><td>distance</td><td>position</td><td>gene name</td><td>gene id</td></tr>
<tr><td>chr7</td><td>4344697</td><td>4344719</td><td>-</td><td>0</td><td>GGTACAGCATAGAGCTCACA</td><td>AGG</td><td>||||||||[||||||||||||]PAM</td><td>0</td><td>Exonic</td><td>Ncr1</td><td>ENSMUSG00000062524</td></tr>
<tr><td>chr11</td><td>101117137</td><td>101117159</td><td>-</td><td>3</td><td>TGAACAGCACAGAGCTCACA</td><td>GGG</td><td>-|-|||||[|-||||||||||]PAM</td><td>27</td><td>Intronic</td><td>Fam134c</td><td>ENSMUSG00000017802</td></tr>
<tr><td>chr10</td><td>43153574</td><td>43153596</td><td>+</td><td>4</td><td>CATAAAGCTTAGAGCTCACA</td><td>TGG</td><td>--||-|||[-|||||||||||]PAM</td><td>620</td><td>Intronic</td><td>Gm29245</td><td>ENSMUSG00000099893</td></tr>
<tr><td>chr8</td><td>26221668</td><td>26221690</td><td>-</td><td>4</td><td>GTTGAAGGATAGAGCTCACA</td><td>CGG</td><td>|-|--||-[||||||||||||]PAM</td><td>2119</td><td>Intergenic</td><td>AC154102.</td><td>ENSMUSG00000105292</td></tr>
<tr><td>chr12</td><td>109718242</td><td>109718264</td><td>-</td><td>4</td><td>AATACAACAAAGAGCTCACA</td><td>GGG</td><td>--||||-|[|-||||||||||]PAM</td><td>490</td><td>Intergenic</td><td>Mir495</td><td>ENSMUSG00000070105</td></tr>
<tr><td>chr11</td><td>79097824</td><td>79097846</td><td>+</td><td>4</td><td>GGAGGAGCACAGAGCTCACA</td><td>GGG</td><td>||---|||[|-||||||||||]PAM</td><td>23338</td><td>Intronic</td><td>Ksr1</td><td>ENSMUSG00000018334</td></tr>
<tr><td>chr11</td><td>79097845</td><td>79097867</td><td>+</td><td>4</td><td>GGAGGAGCACAGAGCTCACA</td><td>GGG</td><td>||---|||[|-||||||||||]PAM</td><td>23359</td><td>Intronic</td><td>Ksr1</td><td>ENSMUSG00000018334</td></tr>
<tr><td>chr11</td><td>79097866</td><td>79097888</td><td>+</td><td>4</td><td>GGAGGAGCACAGAGCTCACA</td><td>GGG</td><td>||---|||[|-||||||||||]PAM</td><td>23380</td><td>Intronic</td><td>Ksr1</td><td>ENSMUSG00000018334</td></tr>
<tr><td>chr11</td><td>79097887</td><td>79097909</td><td>+</td><td>4</td><td>GGAGGAGCACAGAGCTCACA</td><td>GGG</td><td>||---|||[|-||||||||||]PAM</td><td>23401</td><td>Intronic</td><td>Ksr1</td><td>ENSMUSG00000018334</td></tr>
<tr><td>chr14</td><td>70469614</td><td>70469636</td><td>+</td><td>4</td><td>TGTTCACCAGAGAGCTCACA</td><td>TGG</td><td>-||-||-|[|-||||||||||]PAM</td><td>782</td><td>Intergenic</td><td>Phyhip</td><td>ENSMUSG00000003469</td></tr>
<tr><td>chr14</td><td>121341693</td><td>121341715</td><td>-</td><td>3</td><td>GGGACAGCTGAGAGCTCACA</td><td>GGG</td><td>||-|||||[--||||||||||]PAM</td><td>4041</td><td>Intronic</td><td>Stk24</td><td>ENSMUSG00000063410</td></tr>
<tr><td>chr13</td><td>74296414</td><td>74296436</td><td>-</td><td>3</td><td>GGCAAAGCATAAAGCTCACA</td><td>AGG</td><td>||-|-|||[|||-||||||||]PAM</td><td>4083</td><td>Intergenic</td><td>Ahrr</td><td>ENSMUSG00000021575</td></tr>
<tr><td>chr12</td><td>97408340</td><td>97408362</td><td>-</td><td>4</td><td>GTTTCAACAAAGAGCTCACA</td><td>TGG</td><td>|-|-||-|[|-||||||||||]PAM</td><td>37227</td><td>Intergenic</td><td>Gm23558</td><td>ENSMUSG00000088448</td></tr>
<tr><td>chrX</td><td>17664394</td><td>17664416</td><td>+</td><td>4</td><td>GGAGCAGGAAAGAGCTCACA</td><td>AGG</td><td>||--|||-[|-||||||||||]PAM</td><td>10014</td><td>Intergenic</td><td>Gm9436</td><td>ENSMUSG00000081614</td></tr>
<tr><td>chr7</td><td>79717687</td><td>79717709</td><td>+</td><td>4</td><td>GGAGCTGCATAAAGCTCACA</td><td>CGG</td><td>||--|-||[|||-||||||||]PAM</td><td>1967</td><td>Intergenic</td><td>Kif7</td><td>ENSMUSG00000050382</td></tr>
<tr><td>chr15</td><td>54535435</td><td>54535457</td><td>+</td><td>4</td><td>GGGACAAAAGAGAGCTCACA</td><td>GGG</td><td>||-|||--[|-||||||||||]PAM</td><td>35909</td><td>Intergenic</td><td>Mal2</td><td>ENSMUSG00000024479</td></tr>
<tr><td>chr9</td><td>63081791</td><td>63081813</td><td>-</td><td>4</td><td>AGCACAGCACTGAGCTCACA</td><td>AGG</td><td>-|-|||||[|--|||||||||]PAM</td><td>51470</td><td>Intergenic</td><td>Gm25064</td><td>ENSMUSG00000094201</td></tr>
<tr><td>chrX</td><td>140139848</td><td>140139870</td><td>+</td><td>4</td><td>TGGACAGCAGAAAGCTCACA</td><td>AGG</td><td>-|-|||||[|-|-||||||||]PAM</td><td>33051</td><td>Intergenic</td><td>Pih1h3b</td><td>ENSMUSG00000042433</td></tr>
<tr><td>chrX</td><td>140140945</td><td>140140967</td><td>+</td><td>4</td><td>TGGACAGCAGAAAGCTCACA</td><td>AGG</td><td>-|-|||||[|-|-||||||||]PAM</td><td>34148</td><td>Intergenic</td><td>Pih1h3b</td><td>ENSMUSG00000042433</td></tr>
<tr><td>chr12</td><td>116552441</td><td>116552463</td><td>-</td><td>4</td><td>GTTGCAGCAGAAAGCTCACA</td><td>AGG</td><td>|-|-||||[|-|-||||||||]PAM</td><td>27936</td><td>Intronic</td><td>Ptprn2</td><td>ENSMUSG00000056553</td></tr>
<tr><td>chr18</td><td>73096631</td><td>73096653</td><td>+</td><td>4</td><td>TGTACAGTGTACAGCTCACA</td><td>AGG</td><td>-||||||-[-||-||||||||]PAM</td><td>NA</td><td>Intergenic</td><td>NA</td><td>NA</td></tr>
<tr><td>chr9</td><td>36428417</td><td>36428439</td><td>-</td><td>4</td><td>GGTTCAGATTATAGCTCACA</td><td>TGG</td><td>|||-|||-[-||-||||||||]PAM</td><td>3445</td><td>Intergenic</td><td>Gm7257</td><td>ENSMUSG00000023093</td></tr>
<tr><td>chr14</td><td>95813416</td><td>95813438</td><td>+</td><td>4</td><td>GCTTCAGCAAAGATCTCACA</td><td>GGG</td><td>|-|-||||[|-|||-||||||]PAM</td><td>67828</td><td>Intergenic</td><td>4921530L2</td><td>ENSMUSG00000034689</td></tr>
<tr><td>chr2</td><td>165352728</td><td>165352750</td><td>-</td><td>3</td><td>GGTACACCAGAGAACTCACA</td><td>CGG</td><td>||||||-|[|-|||-||||||]PAM</td><td>0</td><td>Exonic</td><td>Zfp663</td><td>ENSMUSG00000056824</td></tr>
<tr><td>chr5</td><td>93302124</td><td>93302146</td><td>+</td><td>4</td><td>GGTGCAGGATGCAGCTCACA</td><td>GGG</td><td>|||-|||-[||--||||||||]PAM</td><td>25893</td><td>Intergenic</td><td>Ccng2</td><td>ENSMUSG00000029385</td></tr>
<tr><td>chr5</td><td>17705106</td><td>17705128</td><td>-</td><td>3</td><td>AGTACAGCACAGAGATCACA</td><td>CGG</td><td>-|||||||[|-||||-|||||]PAM</td><td>6258</td><td>Intronic</td><td>Sema3c</td><td>ENSMUSG00000028780</td></tr>
<tr><td>chr5</td><td>59945436</td><td>59945458</td><td>-</td><td>4</td><td>TGTACATCATTGTGCTCACA</td><td>TGG</td><td>-|||||-|[||-|-|||||||]PAM</td><td>8053</td><td>Intergenic</td><td>Gm43041</td><td>ENSMUSG00000104563</td></tr>
<tr><td>chr12</td><td>117500749</td><td>117500771</td><td>+</td><td>4</td><td>GGCATAGCACAGACCTCACA</td><td>TGG</td><td>||-|-|||[|-|||-||||||]PAM</td><td>15708</td><td>Intergenic</td><td>Rapgef5</td><td>ENSMUSG00000041992</td></tr>
<tr><td>chr16</td><td>8701917</td><td>8701939</td><td>+</td><td>4</td><td>AGCACAGCACAGAGGTCACA</td><td>CGG</td><td>-|-|||||[|-||||-|||||]PAM</td><td>23</td><td>Intronic</td><td>Usp7</td><td>ENSMUSG00000022710</td></tr>
<tr><td>chr5</td><td>35843848</td><td>35843870</td><td>+</td><td>4</td><td>TGTAGAGCAGAGAGGTCACA</td><td>AGG</td><td>-|||-|||[|-||||-|||||]PAM</td><td>582</td><td>Intronic</td><td>Ablim2</td><td>ENSMUSG00000029095</td></tr>
<tr><td>chr18</td><td>65593161</td><td>65593183</td><td>+</td><td>4</td><td>GGCCCAGCAGAGAGGTCACA</td><td>GGG</td><td>||--||||[|-||||-|||||]PAM</td><td>6537</td><td>Intronic</td><td>Zfp532</td><td>ENSMUSG00000042439</td></tr>
<tr><td>chr19</td><td>54151325</td><td>54151347</td><td>-</td><td>3</td><td>GGTAAAGCATACAGGTCACA</td><td>TGG</td><td>||||-|||[|||-||-|||||]PAM</td><td>NA</td><td>Intergenic</td><td>NA</td><td>NA</td></tr>
<tr><td>chr19</td><td>54151281</td><td>54151303</td><td>-</td><td>3</td><td>GGTAAAGCATACAGGTCACA</td><td>TGG</td><td>||||-|||[|||-||-|||||]PAM</td><td>NA</td><td>Intergenic</td><td>NA</td><td>NA</td></tr>
<tr><td>chr19</td><td>54150644</td><td>54150666</td><td>-</td><td>3</td><td>GGTAAAGCATACAGGTCACA</td><td>TGG</td><td>||||-|||[|||-||-|||||]PAM</td><td>NA</td><td>Intergenic</td><td>NA</td><td>NA</td></tr>
<tr><td>chr19</td><td>54150753</td><td>54150775</td><td>-</td><td>3</td><td>GGTAAAGCATACAGGTCACA</td><td>TGG</td><td>||||-|||[|||-||-|||||]PAM</td><td>NA</td><td>Intergenic</td><td>NA</td><td>NA</td></tr>
<tr><td>chr19</td><td>54151611</td><td>54151633</td><td>-</td><td>3</td><td>GGTAAAGCATACAGGTCACA</td><td>TGG</td><td>||||-|||[|||-||-|||||]PAM</td><td>NA</td><td>Intergenic</td><td>NA</td><td>NA</td></tr>
<tr><td>chr19</td><td>54150885</td><td>54150907</td><td>-</td><td>3</td><td>GGTAAAGCATACAGGTCACA</td><td>TGG</td><td>||||-|||[|||-||-|||||]PAM</td><td>NA</td><td>Intergenic</td><td>NA</td><td>NA</td></tr>
<tr><td>chr19</td><td>54151017</td><td>54151039</td><td>-</td><td>3</td><td>GGTAAAGCATACAGGTCACA</td><td>TGG</td><td>||||-|||[|||-||-|||||]PAM</td><td>NA</td><td>Intergenic</td><td>NA</td><td>NA</td></tr>
<tr><td>chr19</td><td>54151721</td><td>54151743</td><td>-</td><td>3</td><td>GGTAAAGCATACAGGTCACA</td><td>TGG</td><td>||||-|||[|||-||-|||||]PAM</td><td>NA</td><td>Intergenic</td><td>NA</td><td>NA</td></tr>
<tr><td>chr19</td><td>54151699</td><td>54151721</td><td>-</td><td>3</td><td>GGTAAAGCATACAGGTCACA</td><td>TGG</td><td>||||-|||[|||-||-|||||]PAM</td><td>NA</td><td>Intergenic</td><td>NA</td><td>NA</td></tr>
<tr><td>chr19</td><td>54151677</td><td>54151699</td><td>-</td><td>3</td><td>GGTAAAGCATACAGGTCACA</td><td>TGG</td><td>||||-|||[|||-||-|||||]PAM</td><td>NA</td><td>Intergenic</td><td>NA</td><td>NA</td></tr>
<tr><td>chr19</td><td>54151655</td><td>54151677</td><td>-</td><td>3</td><td>GGTAAAGCATACAGGTCACA</td><td>TGG</td><td>||||-|||[|||-||-|||||]PAM</td><td>NA</td><td>Intergenic</td><td>NA</td><td>NA</td></tr>
<tr><td>chr19</td><td>54151633</td><td>54151655</td><td>-</td><td>3</td><td>GGTAAAGCATACAGGTCACA</td><td>TGG</td><td>||||-|||[|||-||-|||||]PAM</td><td>NA</td><td>Intergenic</td><td>NA</td><td>NA</td></tr>
<tr><td>chr19</td><td>54151215</td><td>54151237</td><td>-</td><td>3</td><td>GGTAAAGCATACAGGTCACA</td><td>TGG</td><td>||||-|||[|||-||-|||||]PAM</td><td>NA</td><td>Intergenic</td><td>NA</td><td>NA</td></tr>
<tr><td>chr19</td><td>54151127</td><td>54151149</td><td>-</td><td>3</td><td>GGTAAAGCATACAGATCACA</td><td>TGG</td><td>||||-|||[|||-||-|||||]PAM</td><td>NA</td><td>Intergenic</td><td>NA</td><td>NA</td></tr>
<tr><td>chr19</td><td>54151347</td><td>54151369</td><td>-</td><td>3</td><td>GGTAAAGCATACAGATCACA</td><td>TGG</td><td>||||-|||[|||-||-|||||]PAM</td><td>NA</td><td>Intergenic</td><td>NA</td><td>NA</td></tr>
<tr><td>chr19</td><td>54151039</td><td>54151061</td><td>-</td><td>3</td><td>GGTAAAGCATACAGATCACA</td><td>TGG</td><td>||||-|||[|||-||-|||||]PAM</td><td>NA</td><td>Intergenic</td><td>NA</td><td>NA</td></tr>
<tr><td>chr19</td><td>54150819</td><td>54150841</td><td>-</td><td>3</td><td>GGTAAAGCATATAGATCACA</td><td>TGG</td><td>||||-|||[|||-||-|||||]PAM</td><td>NA</td><td>Intergenic</td><td>NA</td><td>NA</td></tr>
<tr><td>chr19</td><td>54150687</td><td>54150709</td><td>-</td><td>3</td><td>GGTAAAGCATACAGTTCACA</td><td>TGG</td><td>||||-|||[|||-||-|||||]PAM</td><td>NA</td><td>Intergenic</td><td>NA</td><td>NA</td></tr>
<tr><td>chr19</td><td>54150973</td><td>54150995</td><td>-</td><td>3</td><td>GGTAAAGCATACAGGTCACA</td><td>CGG</td><td>||||-|||[|||-||-|||||]PAM</td><td>NA</td><td>Intergenic</td><td>NA</td><td>NA</td></tr>
<tr><td>chr19</td><td>54151501</td><td>54151523</td><td>-</td><td>3</td><td>GGTAAAGCATACAGGTCACA</td><td>CGG</td><td>||||-|||[|||-||-|||||]PAM</td><td>NA</td><td>Intergenic</td><td>NA</td><td>NA</td></tr>
<tr><td>chr10</td><td>82060609</td><td>82060631</td><td>+</td><td>4</td><td>GGTACACAAAAGAACTCACA</td><td>CGG</td><td>||||||--[|-|||-||||||]PAM</td><td>0</td><td>Exonic</td><td>Zfp873</td><td>ENSMUSG00000061371</td></tr>
<tr><td>chr8</td><td>107089592</td><td>107089614</td><td>-</td><td>3</td><td>GGTAAAGCACAGAGCCCACA</td><td>GGG</td><td>||||-|||[|-|||||-||||]PAM</td><td>5208</td><td>Intronic</td><td>Terf2</td><td>ENSMUSG00000031921</td></tr>
<tr><td>chr19</td><td>54150929</td><td>54150951</td><td>-</td><td>4</td><td>GATAAAGCATACAGGTCACA</td><td>TGG</td><td>|-||-|||[|||-||-|||||]PAM</td><td>NA</td><td>Intergenic</td><td>NA</td><td>NA</td></tr>
<tr><td>chr19</td><td>54151259</td><td>54151281</td><td>-</td><td>4</td><td>GATAAAGCATACAGGTCACA</td><td>TGG</td><td>|-||-|||[|||-||-|||||]PAM</td><td>NA</td><td>Intergenic</td><td>NA</td><td>NA</td></tr>
<tr><td>chr19</td><td>54151391</td><td>54151413</td><td>-</td><td>4</td><td>GATAAAGCATACAGGTCACA</td><td>TGG</td><td>|-||-|||[|||-||-|||||]PAM</td><td>NA</td><td>Intergenic</td><td>NA</td><td>NA</td></tr>
<tr><td>chr19</td><td>54151567</td><td>54151589</td><td>-</td><td>4</td><td>GATAAAGCATACAGGTCACA</td><td>TGG</td><td>|-||-|||[|||-||-|||||]PAM</td><td>NA</td><td>Intergenic</td><td>NA</td><td>NA</td></tr>
<tr><td>chr19</td><td>54151545</td><td>54151567</td><td>-</td><td>4</td><td>GATAAAGCATACAGGTCACA</td><td>TGG</td><td>|-||-|||[|||-||-|||||]PAM</td><td>NA</td><td>Intergenic</td><td>NA</td><td>NA</td></tr>
<tr><td>chr19</td><td>54151479</td><td>54151501</td><td>-</td><td>4</td><td>GATAAAGCATACAGGTCACA</td><td>TGG</td><td>|-||-|||[|||-||-|||||]PAM</td><td>NA</td><td>Intergenic</td><td>NA</td><td>NA</td></tr>
<tr><td>chr19</td><td>54151237</td><td>54151259</td><td>-</td><td>4</td><td>GATAAAGCATACAGGTCACA</td><td>TGG</td><td>|-||-|||[|||-||-|||||]PAM</td><td>NA</td><td>Intergenic</td><td>NA</td><td>NA</td></tr>
<tr><td>chr19</td><td>54151303</td><td>54151325</td><td>-</td><td>4</td><td>GATAAAGCATACAGGTCACA</td><td>TGG</td><td>|-||-|||[|||-||-|||||]PAM</td><td>NA</td><td>Intergenic</td><td>NA</td><td>NA</td></tr>
<tr><td>chr19</td><td>54151369</td><td>54151391</td><td>-</td><td>4</td><td>GATAAAGCATACAGGTCACA</td><td>TGG</td><td>|-||-|||[|||-||-|||||]PAM</td><td>NA</td><td>Intergenic</td><td>NA</td><td>NA</td></tr>
<tr><td>chr19</td><td>54150731</td><td>54150753</td><td>-</td><td>4</td><td>GATAAAGCATACAGATCACA</td><td>TGG</td><td>|-||-|||[|||-||-|||||]PAM</td><td>NA</td><td>Intergenic</td><td>NA</td><td>NA</td></tr>
<tr><td>chr19</td><td>54150622</td><td>54150644</td><td>-</td><td>4</td><td>GATAAAGCATACAGATCACA</td><td>TGG</td><td>|-||-|||[|||-||-|||||]PAM</td><td>NA</td><td>Intergenic</td><td>NA</td><td>NA</td></tr>
<tr><td>chr19</td><td>54151061</td><td>54151083</td><td>-</td><td>4</td><td>GATAAAGCATACAGATCACA</td><td>TGG</td><td>|-||-|||[|||-||-|||||]PAM</td><td>NA</td><td>Intergenic</td><td>NA</td><td>NA</td></tr>
<tr><td>chr19</td><td>54150907</td><td>54150929</td><td>-</td><td>4</td><td>GATAAAGCATATAGATCACA</td><td>TGG</td><td>|-||-|||[|||-||-|||||]PAM</td><td>NA</td><td>Intergenic</td><td>NA</td><td>NA</td></tr>
<tr><td>chr19</td><td>54150951</td><td>54150973</td><td>-</td><td>4</td><td>GATAAAGCATACAGATCACA</td><td>CGG</td><td>|-||-|||[|||-||-|||||]PAM</td><td>NA</td><td>Intergenic</td><td>NA</td><td>NA</td></tr>
<tr><td>chr19</td><td>54151413</td><td>54151435</td><td>-</td><td>4</td><td>GATAAAGCATACAGATCACA</td><td>CGG</td><td>|-||-|||[|||-||-|||||]PAM</td><td>NA</td><td>Intergenic</td><td>NA</td><td>NA</td></tr>
<tr><td>chr19</td><td>54151589</td><td>54151611</td><td>-</td><td>4</td><td>GATAAAGCATACAGATCACA</td><td>CGG</td><td>|-||-|||[|||-||-|||||]PAM</td><td>NA</td><td>Intergenic</td><td>NA</td><td>NA</td></tr>
<tr><td>chr19</td><td>54151193</td><td>54151215</td><td>-</td><td>4</td><td>GGCAAAGCATACAGATCACA</td><td>TGG</td><td>||-|-|||[|||-||-|||||]PAM</td><td>NA</td><td>Intergenic</td><td>NA</td><td>NA</td></tr>
<tr><td>chr14</td><td>61337907</td><td>61337929</td><td>-</td><td>3</td><td>AGTACAGCAGAGAGCTGACA</td><td>GGG</td><td>-|||||||[|-|||||-|||]PAM</td><td>1834</td><td>Intergenic</td><td>Ebpl</td><td>ENSMUSG00000021928</td></tr>
<tr><td>chr11</td><td>16091670</td><td>16091692</td><td>+</td><td>4</td><td>GGAAGAGCATTGAGCACACA</td><td>GGG</td><td>||-|-|||[||-||||-||||]PAM</td><td>53242</td><td>Intergenic</td><td>Gm37319</td><td>ENSMUSG00000104323</td></tr>
<tr><td>chr19</td><td>54150797</td><td>54150819</td><td>-</td><td>4</td><td>GGTAAAACATACAGGTCACA</td><td>TGG</td><td>||||-|-|[|||-||-|||||]PAM</td><td>NA</td><td>Intergenic</td><td>NA</td><td>NA</td></tr>
<tr><td>chr15</td><td>72859933</td><td>72859955</td><td>+</td><td>2</td><td>GGTACAGCATACAGCTGACA</td><td>CGG</td><td>||||||||[|||-||||-|||]PAM</td><td>7420</td><td>Intronic</td><td>Trappc9</td><td>ENSMUSG00000047921</td></tr>
<tr><td>chr2</td><td>60185827</td><td>60185849</td><td>+</td><td>4</td><td>GGTACAAAACAGAGCACACA</td><td>GGG</td><td>||||||--[|-|||||-||||]PAM</td><td>20277</td><td>Intronic</td><td>Baz2b</td><td>ENSMUSG00000026987</td></tr>
<tr><td>chr6</td><td>141177374</td><td>141177396</td><td>-</td><td>4</td><td>GGTACAAAATAGGGTTCACA</td><td>TGG</td><td>||||||--[||||-|-|||||]PAM</td><td>38798</td><td>Intergenic</td><td>RP23-97L1</td><td>ENSMUSG00000107462</td></tr>
<tr><td>chr11</td><td>28383621</td><td>28383643</td><td>-</td><td>4</td><td>TGTACCCCATAGAGCTCTCA</td><td>GGG</td><td>-||||--|[|||||||||-||]PAM</td><td>2042</td><td>Intergenic</td><td>Ccdc85a</td><td>ENSMUSG00000032878</td></tr>
<tr><td>chr1</td><td>184482806</td><td>184482828</td><td>-</td><td>4</td><td>GGTAGCTCATAGAGCTCGCA</td><td>GGG</td><td>||||---|[|||||||||-||]PAM</td><td>39072</td><td>Intergenic</td><td>2900092O1</td><td>ENSMUSG00000104419</td></tr>
<tr><td>chr16</td><td>44679251</td><td>44679273</td><td>+</td><td>4</td><td>GGTACAATATAGATTTCACA</td><td>GGG</td><td>||||||--[|||||--|||||]PAM</td><td>45028</td><td>Intergenic</td><td>BC027231</td><td>ENSMUSG00000036208</td></tr>
<tr><td>chr7</td><td>36982017</td><td>36982039</td><td>+</td><td>4</td><td>GGTGGAGCCTAGAGCTCTCA</td><td>GGG</td><td>|||--|||[-||||||||-||]PAM</td><td>NA</td><td>Intergenic</td><td>NA</td><td>NA</td></tr>
<tr><td>chr1</td><td>91377818</td><td>91377840</td><td>-</td><td>4</td><td>GGCAGAGCATAGAGGCCACA</td><td>CGG</td><td>||-|-|||[||||||--||||]PAM</td><td>626</td><td>Intronic</td><td>Ilkap</td><td>ENSMUSG00000026309</td></tr>
<tr><td>chr13</td><td>115225522</td><td>115225544</td><td>-</td><td>4</td><td>GGTACCCCACAGAGCTCCCA</td><td>GGG</td><td>|||||--|[|-|||||||-||]PAM</td><td>NA</td><td>Intergenic</td><td>NA</td><td>NA</td></tr>
<tr><td>chr10</td><td>45163510</td><td>45163532</td><td>+</td><td>4</td><td>AGTACAGAATAGAGGACACA</td><td>TGG</td><td>-||||||-[||||||--||||]PAM</td><td>4513</td><td>Intergenic</td><td>Prep</td><td>ENSMUSG00000019849</td></tr>
<tr><td>chr19</td><td>23977003</td><td>23977025</td><td>+</td><td>4</td><td>GGGACAGAATAGAGACCACA</td><td>AGG</td><td>||-||||-[||||||--||||]PAM</td><td>937</td><td>Intronic</td><td>Fam189a2</td><td>ENSMUSG00000071604</td></tr>
<tr><td>chr1</td><td>161195181</td><td>161195203</td><td>-</td><td>4</td><td>GTTAGAGAATAGAGCTCAAA</td><td>GGG</td><td>|-||-||-[||||||||||-|]PAM</td><td>224</td><td>Intronic</td><td>Gm6185</td><td>ENSMUSG00000102817</td></tr>
<tr><td>chr13</td><td>45577979</td><td>45578001</td><td>-</td><td>4</td><td>GATACAGGATACAGCTCTCA</td><td>GGG</td><td>|-|||||-[|||-|||||-||]PAM</td><td>9399</td><td>Intronic</td><td>Atxn1</td><td>ENSMUSG00000046876</td></tr>
<tr><td>chr14</td><td>31278800</td><td>31278822</td><td>-</td><td>4</td><td>GGACCAGCAGAGAGCTCAGA</td><td>TGG</td><td>||--||||[|-||||||||-|]PAM</td><td>233</td><td>Intronic</td><td>Dnah1</td><td>ENSMUSG00000019027</td></tr>
<tr><td>chr13</td><td>63300095</td><td>63300117</td><td>-</td><td>4</td><td>GGCATAGCACAGAGCTCATA</td><td>GGG</td><td>||-|-|||[|-||||||||-|]PAM</td><td>367</td><td>Intronic</td><td>Mir23b</td><td>ENSMUSG00000065599</td></tr>
<tr><td>chr7</td><td>117250975</td><td>117250997</td><td>-</td><td>4</td><td>GGGACTGCAGAGAGCTCAGA</td><td>AGG</td><td>||-||-||[|-||||||||-|]PAM</td><td>21193</td><td>Intronic</td><td>RP23-174K</td><td>ENSMUSG00000108303</td></tr>
<tr><td>chr5</td><td>103072951</td><td>103072973</td><td>+</td><td>4</td><td>CATACAGCATAGAGCGAACA</td><td>GGG</td><td>--||||||[|||||||--|||]PAM</td><td>14559</td><td>Intronic</td><td>Mapk10</td><td>ENSMUSG00000046709</td></tr>
<tr><td>chr15</td><td>31218750</td><td>31218772</td><td>+</td><td>4</td><td>GGCACAACACAGAGCTCAGA</td><td>AGG</td><td>||-|||-|[|-||||||||-|]PAM</td><td>5542</td><td>Intergenic</td><td>Dap</td><td>ENSMUSG00000039168</td></tr>
<tr><td>chr7</td><td>12721234</td><td>12721256</td><td>+</td><td>4</td><td>GGTATAGAAGAGAGCTCAAA</td><td>GGG</td><td>||||-||-[|-||||||||-|]PAM</td><td>5210</td><td>Intronic</td><td>Vmn2r56</td><td>ENSMUSG00000090762</td></tr>
<tr><td>chr5</td><td>5212943</td><td>5212965</td><td>+</td><td>4</td><td>GGTAACGCATAGAACTCCCA</td><td>TGG</td><td>||||--||[|||||-|||-||]PAM</td><td>14199</td><td>Intronic</td><td>Cdk14</td><td>ENSMUSG00000028926</td></tr>
<tr><td>chr9</td><td>99766190</td><td>99766212</td><td>-</td><td>3</td><td>GGAACAGCCTAGAGCTCACT</td><td>TGG</td><td>||-|||||[-||||||||||-]PAM</td><td>10147</td><td>Intergenic</td><td>Gm16004</td><td>ENSMUSG00000089655</td></tr>
<tr><td>chr5</td><td>129268165</td><td>129268187</td><td>-</td><td>4</td><td>GGTGAGGCATAGAGCTCACC</td><td>TGG</td><td>|||---||[|||||||||||-]PAM</td><td>63566</td><td>Intergenic</td><td>Adgrd1</td><td>ENSMUSG00000044017</td></tr>
<tr><td>chr7</td><td>112937138</td><td>112937160</td><td>+</td><td>3</td><td>GGGACAGCATGGAGCTCACT</td><td>CGG</td><td>||-|||||[||-||||||||-]PAM</td><td>16802</td><td>Intergenic</td><td>Rassf10</td><td>ENSMUSG00000098132</td></tr>
<tr><td>chr16</td><td>11938768</td><td>11938790</td><td>-</td><td>4</td><td>AGTACAGTATAGAGGTCTCA</td><td>GGG</td><td>-||||||-[||||||-||-||]PAM</td><td>8168</td><td>Intergenic</td><td>Gm9961</td><td>ENSMUSG00000055015</td></tr>
</table>

<table>
<tr><td>chr7</td><td>74729747</td><td>74729769</td><td>+</td><td>4</td><td>TGTTCAGCATAGATCTCAGA</td><td>GGG</td><td>-||-||||[|||||-||||-|]PAM</td><td>29661</td><td>Intergenic</td><td>RP23-53N2</td><td>ENSMUSG00000108570</td></tr>
<tr><td>chr1</td><td>36791823</td><td>36791845</td><td>-</td><td>4</td><td>GTGACAGCATAGAGCCCTCA</td><td>GGG</td><td>|--|||||[|||||||-|-||]PAM</td><td>346</td><td>Intergenic</td><td>Tmem131</td><td>ENSMUSG00000026116</td></tr>
<tr><td>chr13</td><td>58937499</td><td>58937521</td><td>+</td><td>4</td><td>GGTCCAGGATAGAGATCTCA</td><td>AGG</td><td>|||-|||-[||||||-||-||]PAM</td><td>469</td><td>Intronic</td><td>Ntrk2</td><td>ENSMUSG00000055254</td></tr>
<tr><td>chr2</td><td>145865815</td><td>145865837</td><td>+</td><td>4</td><td>TGTACAGTCTAGAGCTCACT</td><td>AGG</td><td>-||||||-[-||||||||||-]PAM</td><td>3118</td><td>Intronic</td><td>Rin2</td><td>ENSMUSG00000001768</td></tr>
<tr><td>chr3</td><td>148998056</td><td>148998078</td><td>+</td><td>4</td><td>AATACAGCATAGAGTTCAGA</td><td>TGG</td><td>--||||||[||||||-|||-|]PAM</td><td>2893</td><td>Intronic</td><td>Gm43573</td><td>ENSMUSG00000104786</td></tr>
<tr><td>chr9</td><td>48905225</td><td>48905247</td><td>+</td><td>4</td><td>GGAAGAGCATAGAGTTCATA</td><td>AGG</td><td>||-|-|||[||||||-|||-|]PAM</td><td>479</td><td>Intronic</td><td>Htr3a</td><td>ENSMUSG00000032269</td></tr>
<tr><td>chr4</td><td>141000365</td><td>141000387</td><td>-</td><td>4</td><td>GGTGCAGGATGGAGCTCACC</td><td>AGG</td><td>|||-|||-[||-||||||||-]PAM</td><td>0</td><td>Exonic</td><td>Atp13a2</td><td>ENSMUSG00000036622</td></tr>
<tr><td>chr3</td><td>20271049</td><td>20271071</td><td>+</td><td>4</td><td>GGTAATGCATAGAGCTGCCA</td><td>AGG</td><td>||||--||[||||||||--||]PAM</td><td>682</td><td>Intronic</td><td>Cpb1</td><td>ENSMUSG00000011463</td></tr>
<tr><td>chr19</td><td>42278658</td><td>42278680</td><td>-</td><td>4</td><td>GGTTCAACATAGTGCTCACG</td><td>GGG</td><td>|||-||-|[||||-||||||-]PAM</td><td>1402</td><td>Intergenic</td><td>Mir3085</td><td>ENSMUSG00000093234</td></tr>
<tr><td>chr17</td><td>9581184</td><td>9581206</td><td>+</td><td>4</td><td>GAGACAGCATAGAGGTCACC</td><td>AGG</td><td>|--|||||[||||||-||||-]PAM</td><td>85291</td><td>Intergenic</td><td>Pabpc6</td><td>ENSMUSG00000046173</td></tr>
<tr><td>chr3</td><td>103598025</td><td>103598047</td><td>+</td><td>4</td><td>GCTGCAGCATAGAGCTGACT</td><td>GGG</td><td>|-|-||||[||||||||-||-]PAM</td><td>3896</td><td>Intronic</td><td>Gm43066</td><td>ENSMUSG00000104764</td></tr>
<tr><td>chr12</td><td>54063919</td><td>54063941</td><td>+</td><td>4</td><td>GGTGCATCATAGAGCTCTGA</td><td>GGG</td><td>|||-||-|[|||||||||--|]PAM</td><td>1672</td><td>Intronic</td><td>1700060O[illegible]</td><td>ENSMUSG00000099407</td></tr>
<tr><td>chr2</td><td>112564742</td><td>112564764</td><td>-</td><td>4</td><td>GATACTGCATAGAGCTGACT</td><td>TGG</td><td>|-|||-||[||||||||-||-]PAM</td><td>4955</td><td>Intronic</td><td>Aven</td><td>ENSMUSG00000003604</td></tr>
<tr><td>chr4</td><td>107121872</td><td>107121894</td><td>+</td><td>4</td><td>GGAACAGGATAGAGCTAACG</td><td>AGG</td><td>||-||||-[||||||||-||-]PAM</td><td>4719</td><td>Intergenic</td><td>Gm12802</td><td>ENSMUSG00000085304</td></tr>
<tr><td>chr11</td><td>97830789</td><td>97830811</td><td>+</td><td>4</td><td>GGTCCCGCATAGAGCTCTCC</td><td>AGG</td><td>|||-|-||[|||||||||-|-]PAM</td><td>2372</td><td>Intronic</td><td>Lasp1</td><td>ENSMUSG00000038366</td></tr>
</table>

| | |
|---|---|
| Species: | Mouse (Mus musculus GRCm38/mm10) |
| Input: | GTGAGCTCTATGCTGTACCCTGG |
| PAM: | NGG |
| Target site length: | 20 |
| Core length: | 12 |
| Core MM: | 2 |
| Total MM: | 4 |

<table>
<tr><td>T1</td><td>GTGAGCTCTATGCTGTACCCTGG</td><td>1000</td><td>CRISPRater score</td><td>0.706960705</td><td></td><td></td><td></td><td></td><td></td><td></td><td></td></tr>
<tr><td>Chromosome</td><td>start</td><td>end</td><td>strand</td><td>MM</td><td>target_seq</td><td>PAM</td><td>alignment</td><td>distance</td><td>position</td><td>gene name</td><td>gene id</td></tr>
<tr><td>chr7</td><td>4344701</td><td>4344723</td><td>+</td><td>0</td><td>GTGAGCTCTATGCTGTACCC</td><td>TGG</td><td>||||||||[||||||||||||]PAM</td><td>0</td><td>Exonic</td><td>Ncr1</td><td>ENSMUSG00000062524</td></tr>
<tr><td>chr8</td><td>122711124</td><td>122711146</td><td>-</td><td>3</td><td>TGCAGCTCTATGCTGTACCC</td><td>AGG</td><td>---|||||[||||||||||||]PAM</td><td>10964</td><td>Intronic</td><td>Gm10612</td><td>ENSMUSG00000090174</td></tr>
<tr><td>chr15</td><td>72859929</td><td>72859951</td><td>-</td><td>2</td><td>GTCAGCTGTATGCTGTACCC</td><td>AGG</td><td>||-||||-[||||||||||||]PAM</td><td>7416</td><td>Intronic</td><td>Trappc9</td><td>ENSMUSG00000047921</td></tr>
<tr><td>chr4</td><td>15472870</td><td>15472892</td><td>+</td><td>4</td><td>CCTAGCTTTATGCTGTACCC</td><td>TGG</td><td>---||||-[||||||||||||]PAM</td><td>13474</td><td>Intergenic</td><td>Gm11856</td><td>ENSMUSG00000080869</td></tr>
<tr><td>chr11</td><td>107890198</td><td>107890220</td><td>-</td><td>3</td><td>GTGAGGCCTCTGCTGTACCC</td><td>TGG</td><td>|||||--|[|-||||||||||]PAM</td><td>5617</td><td>Intronic</td><td>Cacng5</td><td>ENSMUSG00000040373</td></tr>
<tr><td>chr14</td><td>70220786</td><td>70220808</td><td>-</td><td>4</td><td>GGGATCTCATTGCTGTACCC</td><td>TGG</td><td>|-||-|||[--||||||||||]PAM</td><td>740</td><td>Intronic</td><td>Ppp3cc</td><td>ENSMUSG00000022092</td></tr>
<tr><td>chr5</td><td>111435286</td><td>111435308</td><td>-</td><td>4</td><td>CTGAGGTTTATCCTGTACCC</td><td>AGG</td><td>-||||-|-[|||-||||||||]PAM</td><td>9576</td><td>Intronic</td><td>Gm43119</td><td>ENSMUSG00000106560</td></tr>
<tr><td>chr10</td><td>62201705</td><td>62201727</td><td>-</td><td>4</td><td>GTATGCTCTGGGCTGTACCC</td><td>AGG</td><td>||--||||[|--||||||||]PAM</td><td>51</td><td>Intronic</td><td>Tspan15</td><td>ENSMUSG00000037031</td></tr>
<tr><td>chr3</td><td>53452692</td><td>53452714</td><td>+</td><td>4</td><td>GTGTGCCCTTGGCTGTACCC</td><td>TGG</td><td>|||-||-|[|--||||||||]PAM</td><td>0</td><td>Exonic</td><td>Gm16206</td><td>ENSMUSG00000085174</td></tr>
<tr><td>chr9</td><td>84253809</td><td>84253831</td><td>-</td><td>4</td><td>GTAAGCTATAGTCTGTACCC</td><td>AGG</td><td>||-||||-[||--||||||||]PAM</td><td>65270</td><td>Intergenic</td><td>Gm26146</td><td>ENSMUSG00000077666</td></tr>
<tr><td>chr4</td><td>70789699</td><td>70789721</td><td>-</td><td>4</td><td>GTGAACTGTATCATGTACCC</td><td>TGG</td><td>||||-||-[|||--|||||||]PAM</td><td>NA</td><td>Intergenic</td><td>NA</td><td>NA</td></tr>
<tr><td>chr14</td><td>99809384</td><td>99809406</td><td>-</td><td>4</td><td>GTGAGAACGATGCTCTACCC</td><td>AGG</td><td>|||||--|[-|||||-|||||]PAM</td><td>16611</td><td>Intergenic</td><td>Gm37093</td><td>ENSMUSG00000103188</td></tr>
<tr><td>chr3</td><td>129559477</td><td>129559499</td><td>-</td><td>4</td><td>CTGAGCCCTAAGCTCTACCC</td><td>TGG</td><td>-|||||-|[||-|||-|||||]PAM</td><td>6894</td><td>Intronic</td><td>Elovl6</td><td>ENSMUSG00000041220</td></tr>
<tr><td>chr19</td><td>44312148</td><td>44312170</td><td>-</td><td>2</td><td>GTGAGCTCTCTGCTGTTCCC</td><td>CGG</td><td>||||||||[|-||||||-|||]PAM</td><td>5284</td><td>Intergenic</td><td>Scd2</td><td>ENSMUSG00000025203</td></tr>
<tr><td>chr3</td><td>128154554</td><td>128154576</td><td>-</td><td>4</td><td>TTGAGTTCTATTCTTTACCC</td><td>TGG</td><td>-||||-||[|||-||-|||||]PAM</td><td>4657</td><td>Intronic</td><td>D030025E(</td><td>ENSMUSG00000105816</td></tr>
<tr><td>chr2</td><td>20251004</td><td>20251026</td><td>-</td><td>4</td><td>CTGAGATCTCTGCTGCACCC</td><td>AGG</td><td>-||||-||[|-|||||-||||]PAM</td><td>14940</td><td>Intronic</td><td>Gm13328</td><td>ENSMUSG00000081128</td></tr>
<tr><td>chr12</td><td>117757465</td><td>117757487</td><td>+</td><td>4</td><td>CTGAGAGCTATGCTGTTCCC</td><td>TGG</td><td>-||||--|[||||||||-|||]PAM</td><td>487</td><td>Intergenic</td><td>Rapgef5</td><td>ENSMUSG00000041992</td></tr>
<tr><td>chr4</td><td>154386676</td><td>154386698</td><td>+</td><td>4</td><td>GCAAGCTCTTTGCTGTCCCC</td><td>AGG</td><td>|--|||||[|-||||||-|||]PAM</td><td>19361</td><td>Intronic</td><td>Prdm16</td><td>ENSMUSG00000039410</td></tr>
<tr><td>chr6</td><td>89585855</td><td>89585877</td><td>-</td><td>4</td><td>CTGTGCTCTGTGCTGTTCCC</td><td>TGG</td><td>-||-||||[|-||||||-|||]PAM</td><td>9582</td><td>Intronic</td><td>Chchd6</td><td>ENSMUSG00000030086</td></tr>
<tr><td>chr3</td><td>127123731</td><td>127123753</td><td>-</td><td>4</td><td>CTGTGCTCTCTGCTGTTCCC</td><td>AGG</td><td>-||-||||[|-||||||-|||]PAM</td><td>14084</td><td>Intronic</td><td>Gm42974</td><td>ENSMUSG00000104963</td></tr>
<tr><td>chr12</td><td>27484930</td><td>27484952</td><td>+</td><td>4</td><td>GTGTGCTTTATGATATACCC</td><td>AGG</td><td>|||-|||-[||||-|-|||||]PAM</td><td>NA</td><td>Intergenic</td><td>NA</td><td>NA</td></tr>
<tr><td>chr11</td><td>98382171</td><td>98382193</td><td>+</td><td>4</td><td>GTGCTCTCTCTGCTGTCCCC</td><td>AGG</td><td>|||--|||[|-||||||-|||]PAM</td><td>1059</td><td>Intergenic</td><td>Stard3</td><td>ENSMUSG00000018167</td></tr>
<tr><td>chr12</td><td>99069758</td><td>99069780</td><td>+</td><td>4</td><td>CTGACCTCTAGGCTGTCCCC</td><td>TGG</td><td>-|||-|||[||-|||||-|||]PAM</td><td>22181</td><td>Intergenic</td><td>Rpl30-ps8</td><td>ENSMUSG00000062582</td></tr>
<tr><td>chr14</td><td>100576873</td><td>100576895</td><td>+</td><td>4</td><td>CAGAGCTCTATTCTGTTCCC</td><td>TGG</td><td>--||||||[|||-||||-|||]PAM</td><td>51506</td><td>Intergenic</td><td>Gm24345</td><td>ENSMUSG00000089331</td></tr>
<tr><td>chr4</td><td>65331551</td><td>65331573</td><td>+</td><td>4</td><td>GTGAGGTAAATGCTGTTCCC</td><td>TGG</td><td>|||||-|-[-|||||||-|||]PAM</td><td>4134</td><td>Intronic</td><td>Pappa</td><td>ENSMUSG00000028370</td></tr>
<tr><td>chr15</td><td>10118082</td><td>10118104</td><td>+</td><td>4</td><td>GTGAGCAGAATGCTGTTCCC</td><td>TGG</td><td>||||||--[-|||||||-|||]PAM</td><td>59134</td><td>Intergenic</td><td>Prlr</td><td>ENSMUSG00000005268</td></tr>
<tr><td>chr11</td><td>112892969</td><td>112892991</td><td>-</td><td>4</td><td>GAGAGATTTATGCTGTATCC</td><td>AGG</td><td>|-|||-|-[|||||||||-||]PAM</td><td>60500</td><td>Intergenic</td><td>4933434M</td><td>ENSMUSG00000020624</td></tr>
<tr><td>chr11</td><td>120840465</td><td>120840487</td><td>-</td><td>4</td><td>TTGAGCTATATCCTGTCCCC</td><td>AGG</td><td>-||||||-[|||-||||-|||]PAM</td><td>6221</td><td>Intronic</td><td>Gm11773</td><td>ENSMUSG00000087638</td></tr>
<tr><td>chr16</td><td>3948036</td><td>3948058</td><td>+</td><td>4</td><td>TTGACCTCTAAGCTGTAACC</td><td>AGG</td><td>-|||-|||[||-||||||-||]PAM</td><td>127</td><td>Intronic</td><td>Nlrc3</td><td>ENSMUSG00000049871</td></tr>
<tr><td>chr10</td><td>77056341</td><td>77056363</td><td>+</td><td>4</td><td>GTGCTCTCTATGCTTGACCC</td><td>AGG</td><td>|||--|||[|||||||--||||]PAM</td><td>514</td><td>Intronic</td><td>Col18a1</td><td>ENSMUSG00000001435</td></tr>
<tr><td>chr9</td><td>19919351</td><td>19919373</td><td>+</td><td>3</td><td>GTAAGCTCTGTGCTGTACAC</td><td>AGG</td><td>||-|||||[|-||||||||-|]PAM</td><td>527</td><td>Intronic</td><td>Olfr77</td><td>ENSMUSG00000051118</td></tr>
<tr><td>chr12</td><td>109012861</td><td>109012883</td><td>+</td><td>3</td><td>GTGAGGTCAATGCTGTACTC</td><td>AGG</td><td>|||||-||[-||||||||||-|]PAM</td><td>12080</td><td>Intronic</td><td>Wdr25</td><td>ENSMUSG00000040877</td></tr>
<tr><td>chr2</td><td>118692329</td><td>118692351</td><td>-</td><td>4</td><td>GCCAGCTCTATGCTATTCCC</td><td>AGG</td><td>|--|||||[||||||-|-|||]PAM</td><td>876</td><td>Intronic</td><td>Pak6</td><td>ENSMUSG00000074923</td></tr>
<tr><td>chr14</td><td>70668990</td><td>70669012</td><td>+</td><td>4</td><td>CTCAGCTCTGTGCTGTACTC</td><td>TGG</td><td>-|-|||||[|-||||||||-|]PAM</td><td>232</td><td>Intronic</td><td>Xpo7</td><td>ENSMUSG00000022100</td></tr>
<tr><td>chr12</td><td>98147122</td><td>98147144</td><td>-</td><td>4</td><td>TAGAGCTCTAAGCTGTACTC</td><td>TGG</td><td>--||||||[||-|||||||-|]PAM</td><td>55160</td><td>Intergenic</td><td>Galc</td><td>ENSMUSG00000021003</td></tr>
<tr><td>chr5</td><td>51576009</td><td>51576031</td><td>+</td><td>4</td><td>CTGAGCACTATGCTCTGCCC</td><td>TGG</td><td>-|||||-|[||||||-|-|||]PAM</td><td>8283</td><td>Intergenic</td><td>Ppargc1a</td><td>ENSMUSG00000029167</td></tr>
<tr><td>chr18</td><td>28698149</td><td>28698171</td><td>-</td><td>4</td><td>GTGTGATCTATGGTGTATCC</td><td>AGG</td><td>|||-|-||[||||-||||-||]PAM</td><td>NA</td><td>Intergenic</td><td>NA</td><td>NA</td></tr>
<tr><td>chr7</td><td>57238632</td><td>57238654</td><td>-</td><td>3</td><td>CTGAGCTCTATGCTTTAGCC</td><td>AGG</td><td>-|||||||[||||||-||-||]PAM</td><td>58895</td><td>Intronic</td><td>Gabrg3</td><td>ENSMUSG00000055026</td></tr>
<tr><td>chr18</td><td>80599559</td><td>80599581</td><td>-</td><td>4</td><td>ATGAACTCTATCCTGTACAC</td><td>TGG</td><td>-|||-|||[|||-||||||-|]PAM</td><td>6624</td><td>Intergenic</td><td>Nfatc1</td><td>ENSMUSG00000033016</td></tr>
<tr><td>chr4</td><td>124944990</td><td>124945012</td><td>+</td><td>4</td><td>CTGAGCCCTATGCTGAGCCC</td><td>TGG</td><td>-|||||-|[|||||||--|||]PAM</td><td>334</td><td>Intergenic</td><td>9930104L0</td><td>ENSMUSG00000044730</td></tr>
<tr><td>chr14</td><td>60010419</td><td>60010441</td><td>+</td><td>4</td><td>GCGAACTTTATGCTGTACCA</td><td>TGG</td><td>|-||-||-[||||||||||||-]PAM</td><td>532</td><td>Intronic</td><td>Atp8a2</td><td>ENSMUSG00000021983</td></tr>
<tr><td>chr4</td><td>56450364</td><td>56450386</td><td>-</td><td>4</td><td>GTGAAGTCTATGCTATAGCC</td><td>AGG</td><td>||||--||[||||||-||-||]PAM</td><td>61784</td><td>Intergenic</td><td>Gm12518</td><td>ENSMUSG00000082613</td></tr>
<tr><td>chr3</td><td>125935842</td><td>125935864</td><td>-</td><td>4</td><td>TTGTGCTCTTTGCTGTACCA</td><td>TGG</td><td>-||-||||[|-||||||||||-]PAM</td><td>2589</td><td>Intronic</td><td>Ugt8a</td><td>ENSMUSG00000032854</td></tr>
<tr><td>chr7</td><td>110865049</td><td>110865071</td><td>+</td><td>4</td><td>GAGAGCTGTTTGCTGTACCA</td><td>TGG</td><td>|-|||||-[|-|||||||||-]PAM</td><td>2096</td><td>Intergenic</td><td>Lyve1</td><td>ENSMUSG00000030787</td></tr>
<tr><td>chr1</td><td>36488533</td><td>36488555</td><td>+</td><td>4</td><td>CTGTGCTCTATGCTGTCTCC</td><td>AGG</td><td>-||-||||[||||||||||--||]PAM</td><td>9435</td><td>Intronic</td><td>Cnnm4</td><td>ENSMUSG00000037408</td></tr>
<tr><td>chr11</td><td>8668626</td><td>8668648</td><td>+</td><td>4</td><td>GTGGGCACTATGCTGTGCCG</td><td>GGG</td><td>|||-||-|[||||||||-||-]PAM</td><td>4091</td><td>Intergenic</td><td>Tns3</td><td>ENSMUSG00000020422</td></tr>
</table>
