## Supplemental Table S3 for "Major histocompatibility complex class I-restricted protection against murine cytomegalovirus requires missing-self recognition by the natural killer cell inhibitory Ly49 receptors"

**Table S3: sgRNAs specific for MCMV *m06* or *m152***

| **Guide ID** | **Target** | **sgRNA Sequence + PAM** | **Forward Oligo (5' to 3')** | **Reverse Oligo (5' to 3')** |
|---|---|---|---|---|
| T303 | *m06* | AGAGTCTTACGTTAAGACAGAGG | caccgAGAGTCTTACGTTAAGACAG | aaacCTGTGTTAACGTAAGACTCTc |
| T18561 | *m152* | TATGGACGTGCGCATATTCGAGG | caccgTATGGACGTGCGCATATTCG | aaacCGAATATGCGCACGTCCATAc |

The names of the sgRNA guides (Guide ID) are provided (see also **Table_S2**) along with the intended MCMV ORF targets, and the sgRNA sequence including the "NGG" protospacer adjacent motif (PAM) site underlined. Two oligos were used to generate duplex DNA used for cloning into the modified px330 *Cas9*-encoding vector (see **Methods**). An additional guanine residue was appended to the forward oligo (and complementary cytosine to the reverse oligo) to enhance sgRNA production from the *U6* promoter in cells.
