## Supplemental Table S4 for "Major histocompatibility complex class I-restricted protection against murine cytomegalovirus requires missing-self recognition by the natural killer cell inhibitory Ly49 receptors"

**Table S4: The primers used for *Ly49* gene sequence analysis**

| Gene | Fwd Primer Sequence (5' to 3') | Rev Primer Sequence (5' to 3') | Amplicon size (nt) |
|---|---|---|---|
| *Klra1/Ly49a* | TCTTCCCTCCCATCTTTGTTCA | TGGGTCAGTCCATGTCAGTG | 439 |
| *Klra2/Ly49b* | ATTGTTCTGCTCTGCGCATC | AGAGTCAGGGTGTTTGGACC | 409 |
| *Klra3/Ly49c* | TCTTCCCTCCCGTCCTTGTA | TGCATGTCAGGGTGTTTGGA | 428 |
| *Klra4/Ly49d* | TCACCCTCATGCATAACTAAGG | CAGTCCATGCTGCAGTGTTT | 421 |
| *Klra5/Ly49e* | CTTCTCCGGGCCCTTGAATC | GGCTGTATCAATGGTAGAATGGC | 414 |
| *Klra6/Ly49f* | CTCCCATACTTGTGCATAATCAAA | GGATCAGTCCATGTCAGGGTG | 427 |
| *Klra7/Ly49g* | AACCAAGCCCCAATGAGATC | GGTCAGTCCATGTCAGGGTG | 411 |
| *Klra8/Ly49h* | GGAACATTTTACTTTTCAATGAAAGCCT | CTGTATCATCAGATCCAGGTACCTTT | 324 |
| *Klra9/Ly49i* | CAAGCCCCGATGAGATGGAT | GGATCAGTCCATGTCAGGGTT | 409 |
| *Klra10/Ly49j* | CAAGCCCCGATGAGATGGAC | GGATCAGTCCATGTCAGGGTA | 409 |
| *Klra13-ps/Ly49m* | CTTTTCTTCTCTCTCACCCTTATGCATAAC | CCCAAGATGAGTGAGCAGGAGGT | 278 |
| *Klra14-ps/Ly49n* | ACTTCTTGTTTCCCAAGAAATGTTTTCTTACTG | CACCCTTTCTCAACCTTCTGTATCACT | 591 |
| *Klra17/Ly49q* | GCCCATCTGGCTTCCTTTCT | TGAGTCCCAGTCAGGGTCAT | 533 |
| *Gm15854/Ly49x* | TTCTCTCTTACCCTCGTGCATAAC | GTGAGGTCAGTCCATGCTGAG | 438 |
| *Gm6584* | AGCCAAGCCCAGATGAAATGA | ACAGGGTTTCTCCCCTGAAA | 428 |

This table provides the primer sequences and amplicon lengths used for amplification and sequencing the *Ly49* region of the murine NKC locus on chromosome 6. See **Methods** for sequencing conditions and **Fig S1** for results.
