## Supplemental Table S5 for "Major histocompatibility complex class I-restricted protection against murine cytomegalovirus requires missing-self recognition by the natural killer cell inhibitory Ly49 receptors"

**Table S5: The primers used for MCMV *m06* and *m152* gene sequence analysis**

| **ORF** | **Fwd Sequence Primer (5' to 3')** | **Rev Sequence Primer (5' to 3')** | **Amplicon size (nt)** |
|---|---|---|---|
| *m06* | CACGCCCAAAATCACGCAAAC | GGCGTAGTCGAATGGTACA | 853 |
| *m152* | CGATGTCATCCTCGGATA | GGCTACTCCCGAAAGAGTAA | 578 |

This table provides the primer sequences and amplicon lengths used for amplification and sequencing the viral ORFs *m06* and *m152* from MCMV. See **Methods** for sequencing conditions and **Fig S5** for results.
